# Optimization of 3-Cyano-7-cyclopropylamino-pyrazolo[1,5-a]pyrimidines Toward the Development of an In Vivo Chemical Probe for CSNK2A

**DOI:** 10.1101/2023.05.15.540828

**Authors:** Xuan Yang, Han Wee Ong, Rebekah J. Dickmander, Jeffery L. Smith, Jason W. Brown, William Tao, Edcon Chang, Nathaniel J. Moorman, Alison D. Axtman, Timothy M. Willson

## Abstract

3-cyano-7-cyclopropylamino-pyrazolo[1,5-a]pyrimidines, including the chemical probe SGC-CK2-1, are potent and selective inhibitors of CSNK2A in cells but have limited utility in animal models due to their poor pharmacokinetic properties. While developing analogs with reduced intrinsic clearance and the potential for sustained exposure in mice, we discovered that Phase II conjugation by GST enzymes was a major metabolic transformation in hepatocytes. A protocol for co-dosing with ethacrynic acid, a covalent reversible GST inhibitor, was developed to improve the exposure of analog **2h** in mice. A double co-dosing protocol, using a combination of ethacrynic acid and irreversible P450 inhibitor 1-aminobenzotriazole increased the blood level of **2h** by 40-fold at a 5 h time point.

**Graphical Abstract:** 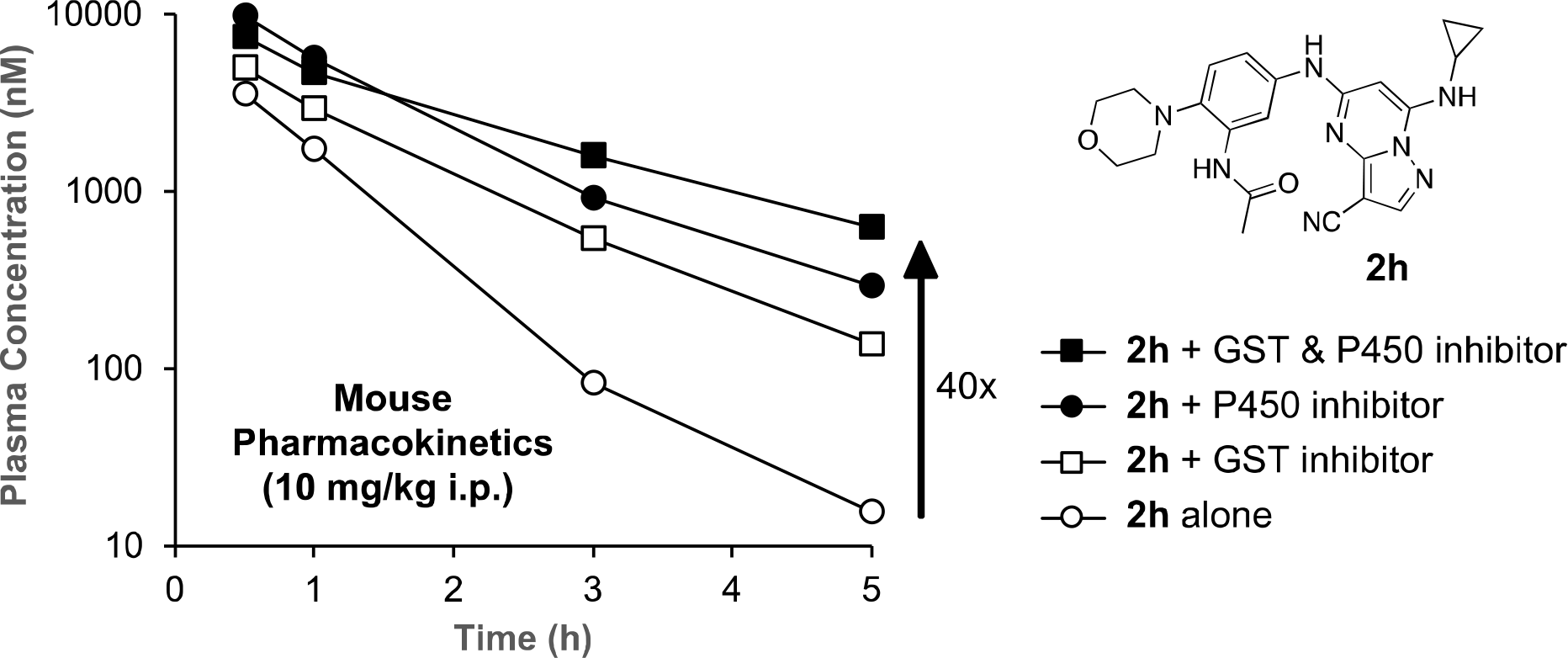

## Introduction

The appearance of a novel coronavirus SARS-CoV-2 and the resulting pandemic have highlighted the need for effective treatments against COVID-19.^1^ Host-directed therapies that target cellular pathways required for virus replication have emerged as a promising approach to combat viral infections.^2^ Protein kinases have been proposed as potential targets for development of antiviral drugs.^3^ Among these kinases, Casein Kinase 2α (CSNK2A) has been shown to play a role in the replication of β-coronaviruses (β-CoV), including SARS-CoV-2.^4–6^. Recent studies have demonstrated the efficacy of CSNK2A inhibitors in reducing the replication of β-CoV in vitro.^6^ However, there has yet to be a demonstration of anti-COVID-19 efficacy in vivo.

3-cyano-7-cyclopropylamino-pyrazolo[1,5-a]pyrimidine (PZP) is a promising chemotype of CSNK2A inhibitor that has demonstrated potent activity in vitro.^7^ PZPs possess good selectivity for CSNK2A over other kinases and display unique chemical features that contribute to their potent inhibitory activity (Figure 1). Specifically, X-ray crystallography studies (Figure 1A) revealed that the 4’-methyl substituent of the chemical probe SGC-CK2-1 (**1a**) enforces an s-cis (*E*)-conformation of the propionamide, allowing it to coordinate with a bound water molecule, which also interacts with the cyano group of the PZP core (Figure 1B). Notably, the alkyl group of the propionamide contributes to the exquisite selectivity of the PZP-based CSNK2A inhibitors over other kinases.^8^ However, despite their promising in vitro activity, PZPs have shown limited bioavailability in rodents by oral dosing due to their low to moderate aqueous solubility, high first-pass metabolism, and rapid clearance.^9, 10^

**Figure 1.**
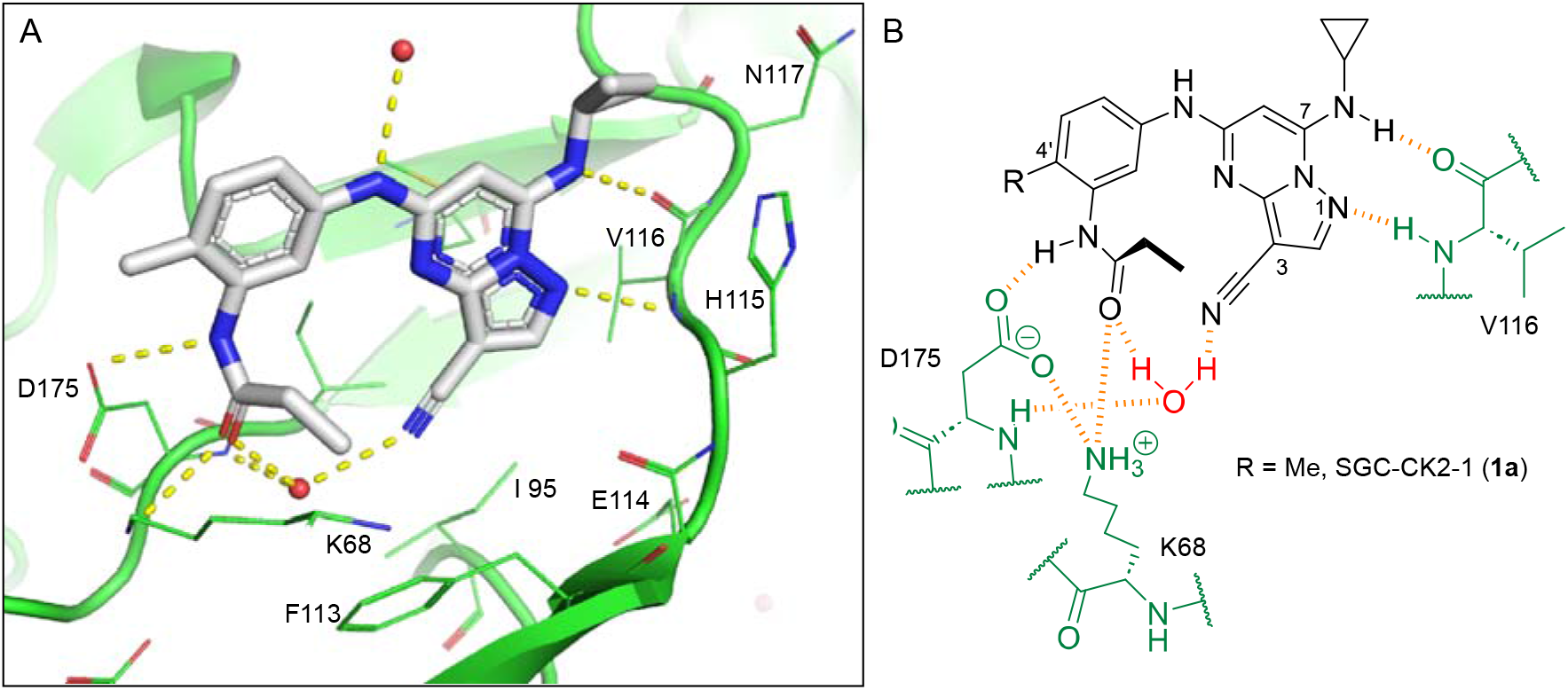
A. X-ray cocrystal structure of SGC-CK2-1 (**1a**) bound to CSNK2A1 (PDB: 6Z83). Image created in PyMOL 2.4.0 with the CSNK2A1 protein shown as a green ribbon and **1a** shown as sticks colored by atom type. Bound water molecules are shown as red spheres. Intermolecular hydrogen bonds are shown as yellow dashed lines. B. Illustration of the key interactions observed in the X-ray structure. The 7-cyclopropylamine and N-1 of the PZP form H-bond interactions with V116 in the hinge region. The 4’-substiuent (Me for **1a**) causes the amide to adopt an s-cis conformation that allows the carbonyl group to interact with K68 and a bound water molecule. The ethyl group of the amide is placed in a pocket that increases selectivity over other kinases. The cyano group of the PZP also interacts with the bound water molecule, which is further coordinated through interaction with the backbone NH of D175.

In this paper, we report our initial studies to identify a PZP-based CSNK2A inhibitor with potent antiviral activity and high sustained blood levels for use in mouse-adapted model of COVID-19 developed for preclinical evaluation of potential drug therapies.^11, 12^ We discovered that the rapid clearance of PZPs in mice was due to pathways of both Phase I and Phase II metabolism. Co-dosing with a glutathione S-transferase (GST) enzyme inhibitor and an irreversible cytochrome P450 inhibitor greatly reduced intrinsic clearance of a PZP analog in mice and may be a viable approach to increase in vivo exposure for preclinical pharmacology studies.

## Results

### Antiviral PZPs with Reduced Phase I Metabolism

All compounds were screened for CSNK2A1 and CSNK2A2 activity using a NanoBRET assay in HEK293 cells and for inhibition of mouse β-CoV replication using an MHV-nLuc assay in DBT cells.^6^ Kinetic solubility was determined using a miniaturized shake flask method. The stability of compounds to Phase I metabolism was determined by incubation with mouse liver microsome for 30 min. SGC-CK2-1 (**1a**) demonstrated low aqueous solubility and rapid metabolism in primary mouse liver microsomes (MLM), with only 40% of the parent compound remaining after 30 min (Table 1). Since hepatic metabolism of **1a** would likely lead to rapid in vivo clearance, we decided to focus initially on increasing its stability in MLM. SMARTCyp,^13, 14^ a web-based predictor of metabolic hot spots, analyzes sites of metabolism using a ligand-based approach from a library of molecular fragments that can be applied across multiple species and isoforms of CYPs.^15^ SMARTCyp suggested that that the cyclopropylamine and the 4’-methyl group might be sites of P450 oxidation (Figure S1). As modification of the cyclopropylamine, which forms part of the hinge-binding motif (Figure 1B), resulted in a large decrease in CSNK2A potency^6, 8^ we opted to study analogs with modification of the 4’-methyl group in order to lower logD, improve aqueous solubility, and possibly reduce P450 metabolism.^16^ Analogs **1b–g** with substituted ethylenediamines replacing the 4’-methyl group demonstrated good CSNK2A potency and retained antiviral activity, as previously reported.^6^ Analogs **1b–g** had ∼50–60-fold improved aqueous solubility. The ethylenediamines **1b–d** also had reduced metabolism in MLM compared to **1a** (Table 1). In contrast the pyrrolidine (**1e**) and piperidine (**1f**) containing diamines were highly metabolized despite their improved aqueous solubility, possibly due to their increased lipophilicity. The piperazine (**1g**) had greatly improved metabolic stability, but at a cost of lower CSNK2A potency and antiviral IC_50_ >1 µM. From this initial series of analogs of **1a** we found that addition of ethylenediamines at the 4’ position resulted in PZPs with good antiviral activity and reduced P450 metabolism in MLM. Although the piperazine **1g** had the best metabolic stability, it showed reduced potency in both CSNK2A and antiviral assays.

**Table 1:** Modification of the 4’-position group of the propionamide series.

| Compound | R | CSNK2A1<br>pIC<sub>50</sub><sup>a</sup> | CSNK2A2<br>pIC<sub>50</sub><sup>a</sup> | MHV<br>pIC<sub>50</sub><sup>b</sup> | cLogP<sup>c</sup> | Kinetic<br>solubility<br>(μg/mL)<sup>d</sup> | MLM<br>metabolism @<br>30 min (%)<sup>e</sup> |
| --- | --- | --- | --- | --- | --- | --- | --- |
| **1a<sup>f</sup>** | Me | 7.4 | 7.4 | 6.9 | 0.38 | 1.4 | 60 |
| **1b<sup>f</sup>** |  | 8.6 | 8.7 | 6.8 | 0.03 | 66 | 38 |
| **1c<sup>f</sup>** |  | 8.2 | 8.3 | 7.1 | 0.33 | 78 | 34 |
| **1d<sup>f</sup>** |  | 7.4 | 7.6 | 6.6 | 0.86 | 81 | 38 |
| **1e<sup>f</sup>** |  | 7.4 | 6.8 | 6.2 | 1.58 | 77 | 94 |
| **1f<sup>f</sup>** |  | 7.0 | 6.6 | 5.9 | 2.15 | 69 | 99 |
| **1g<sup>f</sup>** |  | 6.7 | 6.7 | 5.8 | -0.03 | 88 | 12 |
| **1h** |  | 7.1 | 7.2 | 6.3 | 0.40 | 65 | 11 |
| **1i** |  | 7.6 | 7.7 | 6.9 | 0.40 | 71 | 10 |

<sup>a</sup>In-cell target engagement by NanoBRET assay, n = 1. <sup>b</sup>Inhibition of MHV-nLuc replication, n = 3 with SE ± 0.1. <sup>c</sup>Values from ChemDraw Professional v16.0. <sup>d</sup>Miniaturized shake flask using CLND detection, n = 1; <sup>e</sup>Metabolism of compound in primary mouse liver microsomes (MLM) calculated as 100 – (% remaining), n = 1. <sup>f</sup>Compound previously reported in reference 6.

To build on these results we designed new analogs with 3-aminomethyl piperidine substituents at the 4’-position to combine the features of both the cyclic and linear diamines. The synthesis employed the methods previously developed for analogs **1b–g** (Scheme 1). Addition of the (*R*) or (*S*)-enantiomer of the *N*-Boc-protected 3-aminomethyl piperidine to the acyl-2-fluoro-5-nitroaniline (**x**, R^1^ = Et) followed by reduction of the nitro group yielded the aniline intermediate (**y**, R^1^ = Et). Palladium-catalyzed Buchwald-Hartwig cross coupling with the 5-chloro-pyrazolo[1,5-a]pyrimidine core followed by *N*-Boc deprotection yielded the 3*R*-isomer (**1h**) and the 3*S*-isomer (**1i**). Both isomers, **1h** and **1i**, demonstrated good aqueous solubility and low metabolism in mouse liver microsomes compared to **1a** (Table 1). The 3*S*-isomer (**1i**) was more potent than the 3*R*-isomer (**1h**) in both the CSNK2A assay and the antiviral assay. Overall, **1i** had comparable potency to **1a** but with improved solubility and metabolic stability.

**Scheme 1.**
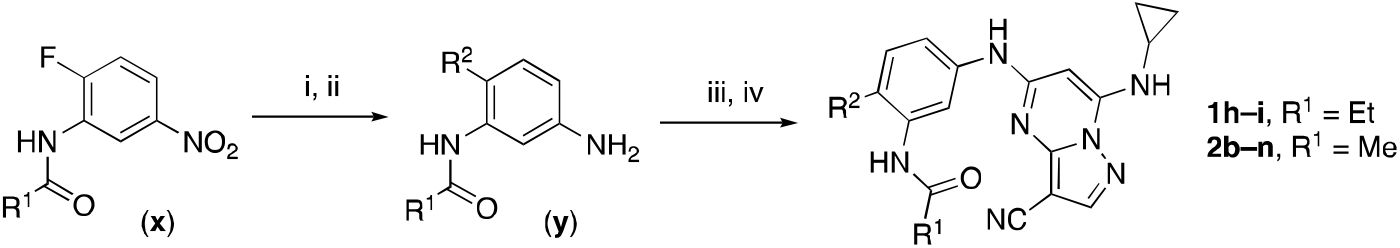
^a^Reagents and conditions: (i) R_2_-H, K_2_CO_3_, MeCN. (ii) H_2_, Pd-C, MeOH. (iii) 5-chloro-7-(cyclopropyl-amino)pyrazolo[1,5-*a*]pyrimidine-3-carbonitrile, BINAP, Pd(OAc)_2_, Cs_2_CO_3_, dioxane, μW, 130°C. (iv) TFA, CH_2_Cl_2_.

To further improve the potency, we synthesized 4’-substituted analogs in the acetamide series of PZPs (Table 2). The compounds were synthesized by the established route from the acyl-2-fluoro-5-nitroaniline (**x**, R^1^ = Me) and the appropriate secondary amine for the R^2^-substituent (Scheme 1). For those R^2^-substituents with chiral centers, the enantiomerically-pure amine was utilized as a building block. The 4’-methyl analog (**2a**) had shown sub-nanomolar activity on CSNK2A and good activity in the antiviral assay.^6^ **2a** had a lower cLogP and slightly improved solubility and metabolic stability compared to the corresponding propionamide (**1a**) (Table 2). Synthesis of the 3*R*-isomer (**2b**) and 3*S*-isomer (**2c**) of the 3-aminomethyl piperidine gave analogs with 10-fold improved potency in the CSNK2A and the antiviral assays while maintaining good metabolic stability. The 3*S*-isomer (**2c**) was again the more potent of the two enantiomers. Additional 4’-substituted analogs were synthesized in the acetamide series. The 3*R*-isomer (**2d**) and 3*S*-isomer (**2e**) of the 3-aminomethyl pyrrolidine also combined potency on CSNK2A and antiviral activity with good metabolic stability. Switching the orientation of the 3-aminomethyl pyrrolidine gave analog **2f** with similar potency but slightly poorer metabolic stability. The piperazine (**2g**) was less potent on CSNK2A and in the antiviral assay, but had good metabolic stability, as had been seen in the corresponding propionamide (**1g**). We had previously prepared and tested morpholine (**2h**),^6^ which had improved antiviral activity compared to **2g**. Although the morpholine (**2h**) showed low aqueous solubility its metabolic stability was much better than the parent analog **2a** (Table 2). Introduction of a hydroxy substituent in **2i** maintained activity and improved aqueous solubility but decreased metabolic stability. The oxo-analogs **2j–n** had lower activity in the antiviral assay and poor metabolic stability, despite some analogs (e.g. **2l**) having low cLogP and good solubility.

**Table 2:** Modification of the 4’-position group of the acetamide series.

| Compound | R | CSNK2A1<br>pIC<sub>50</sub><sup>a</sup> | CSNK2A2<br>pIC<sub>50</sub><sup>a</sup> | MHV<br>pIC<sub>50</sub><sup>b</sup> | cLogP<sup>c</sup> | Kinetic<br>solubility<br>(μg/mL)<sup>d</sup> | MLM<br>metabolism @<br>30 min (%)<sup>e</sup> |
| --- | --- | --- | --- | --- | --- | --- | --- |
| **2a<sup>f</sup>** | Me | 9.3 | 9.2 | 7.1 | -0.15 | 2.9 | 52 |
| **2b** |  | 7.8 | 8.0 | 7.0 | -0.13 | 52 | 14 |
| **2c** |  | 8.3 | 8.4 | 7.9 | -0.13 | 54 | 14 |
| **2d** |  | 8.6 | 8.8 | 7.5 | -0.69 | 64 | 8 |
| **2e** |  | 8.3 | 8.4 | 7.7 | -0.69 | 49 | 16 |
| **2f** |  | 8.4 | 8.5 | 7.9 | 0.50 | 68 | 24 |
| **2g** |  | 7.1 | 7.6 | 5.6 | -0.56 | 52 | 18 |
| **2h<sup>f</sup>** |  | 7.5 | 7.7 | 7.1 | -0.54 | 1.0 | 30 |
| **2i** |  | 7.5 | 7.7 | 7.0 | -0.50 | 37 | 55 |
| **2j** |  | 7.6 | 7.7 | 6.8 | 0.00 | 0.9 | 92 |
| **2k** |  | 6.9 | 7.0 | 6.5 | 0.03 | 21 | 97 |
| **2l** |  | 6.9 | 7.1 | 6.3 | 0.03 | 25 | 100 |
| **2m** |  | 8.0 | 8.1 | 6.4 | -0.23 | 2.4 | 76 |
| **2n** |  | 7.0 | 7.5 | 6.4 | -0.70 | 0.2 | 100 |

<sup>a</sup>In-cell target engagement by NanoBRET assay, n = 1. <sup>b</sup>Inhibition of MHV-nLuc replication, n = 3 with SE ± 0.1. <sup>c</sup>Values from ChemDraw Professional v16.0. <sup>d</sup>Miniaturized shake flask using CLND detection, n = 1; <sup>e</sup>Metabolism of compound in primary mouse liver microsomes (MLM) calculated as 100 – (% remaining), n = 1. <sup>f</sup>Compound previously reported in reference 6.

### Pharmacokinetic Profiling of PZPs

Having identified several analogs with improved metabolic stability in MLM and potent activity in the CSNK2A and antiviral assays, we selected three compounds (**1i**, **2c**, and **2e**) for pharmacokinetic studies. These analogs showed only 10– 16% metabolism in primary MLM after 30 min incubation. The compounds were dosed at 1.0 mg/kg i.v. and 10 mg/kg i.p. to mice and the level of the parent compound was monitored over 24 h (Table 3). Despite their stability in MLM, all three compounds demonstrated moderate to high clearance following i.v. dosing, with t_1/2_ of 1.2–1.8 h. Similar results were observed following i.p. dosing with t_1/2_ of 1.2–1.5 h. Although the bioavailability by i.p. dosing was good, the rapid clearance resulted in low total exposure and blood levels that fell below the IC_50_ for antiviral activity within 1 h.

**Table 3:** In vivo mouse pharmacokinetic parameters of 1i, 2c, and 2e***^a^***.

| Compound | i.v. (1 mg/kg) |  |  |  |  | i.p. (10 mg/kg) |  |  |  |
| --- | --- | --- | --- | --- | --- | --- | --- | --- | --- |
|  | C<sub>max</sub><br>(ng/mL) | AUC<sub>last</sub><br>(ng.h/mL) | Vd<sub>ss</sub><br>(L/kg) | t<sub>1/2</sub><br>(h) | CL<sub>int</sub><br>(mL/min/kg) | C<sub>max</sub><br>(ng/mL) | AUC<sub>last</sub><br>(ng.h/mL) | t<sub>1/2</sub><br>(h) | F<br>(%) |
| **1i** | 1100 | 670 | 2.60 | 1.3 | 36 | 4700 | 7010 | 1.2 | 100 |
| **2c** | 350 | 200 | 5.07 | 1.8 | 84 | 670 | 770 | 1.5 | 40 |
| **2e** | 810 | 250 | 2.67 | 1.2 | 69 | 1500 | 1300 | 1.4 | 53 |

<sup>a</sup>Mice dosed by i.v. or i.p. route of administration, n = 3. Compounds quantified by LC/MS.

### Phase II Metabolism of PZPs

To further explore the reason for the rapid in vivo clearance of the PZPs (**1i**, **2c**, and **2e**), we performed time course studies to measure their rate of intrinsic clearance using in vitro systems of hepatic metabolism (Table 4). In mouse primary liver microsomes the three compounds showed low intrinsic clearance consistent with the single time point measurements (Tables 1 and 2). Repeating the experiment with primary liver microsomes that included the S9 fraction resulted in a decrease in clearance indicating that metabolic stability was unaffected by inclusion of the cytosolic enzymes. However, when incubated with primary mouse hepatocytes, all three compounds were rapidly metabolized with high rates of intrinsic clearance (Table 4). The results suggested that a non-P450 metabolic pathway could be responsible for the rapid hepatic clearance of the PZPs in hepatocytes and in mice in vivo.^17^

**Table 4:** In vitro mouse metabolism of 1i, 2c, and 2e***^a^***.

| Compound | Liver Microsomes | S9 fraction | Hepatocytes |
| --- | --- | --- | --- |
|  | CL<sub>int</sub> (mL/min/kg) | CL<sub>int</sub> (mL/min/kg) | CL<sub>int</sub> (mL/min/kg) |
| **1i** | <24 | 0 | 143 |
| **2c** | <24 | 1 | 132 |
| **2e** | 27 | 4 | 69 |

<sup>a</sup>Compounds quantified by LC/MS at 5 time points over 5h, n = 1.

### Metabolite ID Study of 1i

To identify the metabolic pathway responsible for the rapid clearance in mouse hepatocytes, we performed a metabolite ID study using the 3*S*-aminomethyl piperidine (**1i**). Incubation of **1i** for 4 h in primary cultures of mouse hepatocytes resulted in the appearance of 12 metabolites by ultrahigh performance liquid chromatography (Figure 2A). Analysis of the metabolites by mass spectroscopy and quantification by UV absorbance enabled the assignment of the primary pathways of metabolic transformation (Table 4, Figures 2B and S2). Six of the metabolites (M1–4, M10–11) resulted from GSH conjugation of the PZPs (Figure 2B), a Phase II metabolic transformation that could only occur in whole hepatocytes that contain the active GST enzymes. GSH conjugation was observed in the major fraction of the total metabolites as quantified by UV absorbance (Table 5) and is likely to be an important pathway of clearance of the PZP **1i** (Figure 2B). In addition to GSH conjugation, demethylation of the secondary amine on the piperidine and dealkylation of the 7-amino group on the PZP were found among the metabolites (Table 4 and Figure 2B). Notably, dealkylation was often found in combination with GSH conjugation. P450 oxidation of electron-rich aromatic rings to an epoxide intermediate is well known activation step in GST-catalyzed GSH conjugation,^18^ making the electron-rich amino-substituted phenyl ring the most likely site of Phase II metabolism (Figure 2C). Although we initially considered the nitrile as a potential site of conjugation, since it can undergo a Pinner reaction with free GSH,^19^ there is no evidence that the reaction is catalyzed by GST enzymes.^20, 21^ The metabolite ID study implicated both cytochrome P450 and GST enzymes as likely to be responsible for the rapid clearance of the PZPs in mouse hepatocytes through a combination of demethylation, dealkylation, and GSH conjugation (Figure 2C).

**Figure 2.**
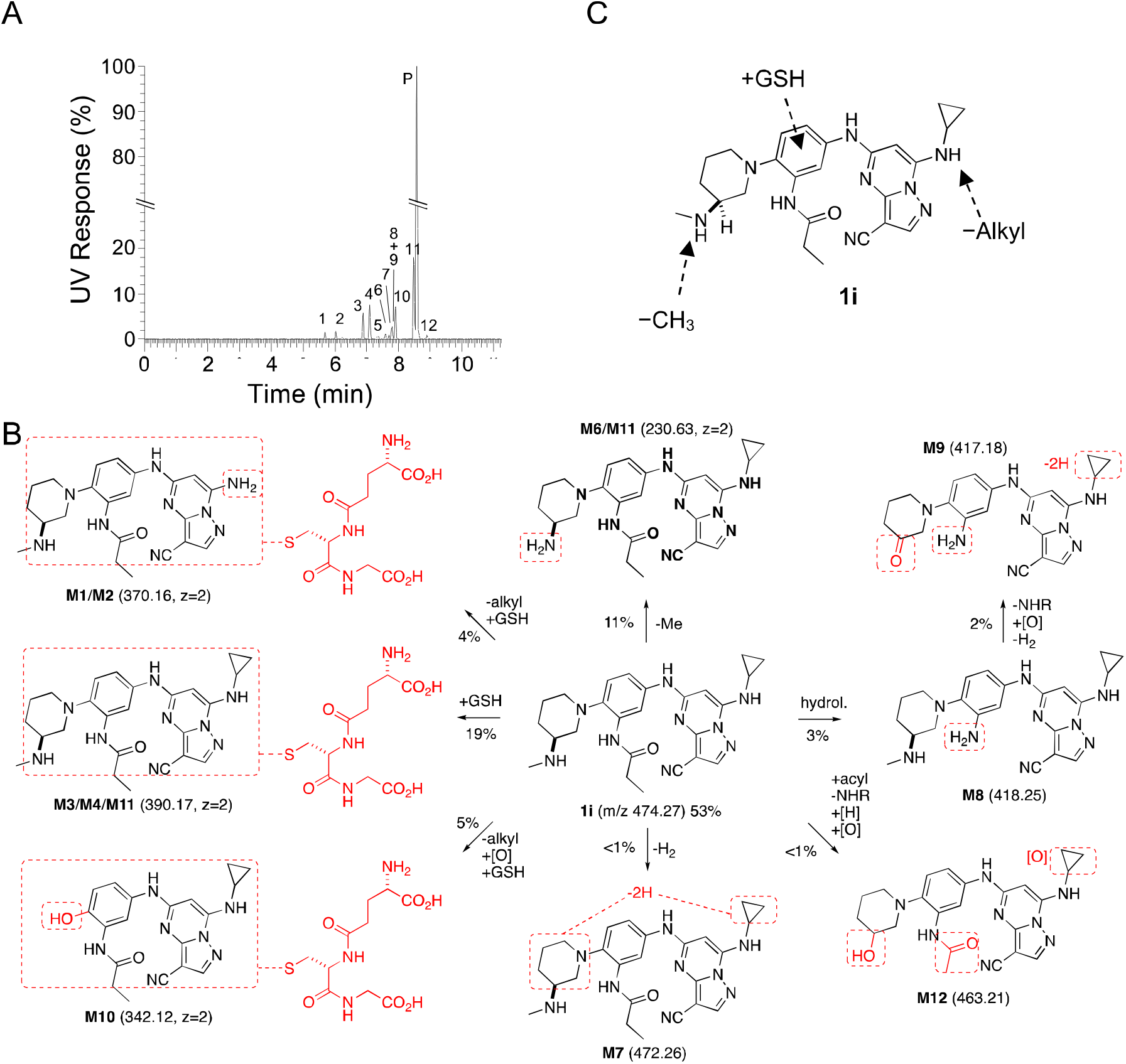
A. Identification of 12 metabolites of **1i** (Parent, P) after incubation in mouse primary hepatocytes for 4 h. The relative quantity of each metabolite was determined by UV. B. Structural assignment of the metabolites (**M1**–**M12**) from MS fragmentation (see Figure S2). Molecular ions shown in parentheses. Doubly charged ions are indicated as z=2. The relative percent contribution of each metabolite taken from Table 5. C. Summary of the major sites of metabolism of **1i** in mouse primary hepatocytes.

**Table 5:** Metabolites of 1i in Mouse Hepatocytes.

| Peak<sup>a</sup> | Time (min) | Mass (m/z) | Transformation |  |  |  |  |  |  |  |  | Rel. (%) |
| --- | --- | --- | --- | --- | --- | --- | --- | --- | --- | --- | --- | --- |
|  |  |  | -Alkyl | +GSH | -CH<sub>3</sub> | -H<sub>2</sub> | Hydrol. | -NH<sub>2</sub> | [O] | [H] | Acyl. |  |
| M1 | 5.67 | 370.16<sup>b</sup> | X | X |  |  |  |  |  |  |  | <1 |
| M2 | 6.03 | 370.16<sup>b</sup> | X | X |  |  |  |  |  |  |  | 3 |
| M3 | 6.89 | 390.17<sup>b</sup> |  | X |  |  |  |  |  |  |  | 9 |
| M4 | 7.10 | 390.17<sup>b</sup> |  | X |  |  |  |  |  |  |  | 9 |
| M5 | 7.34 | 390.17<sup>b</sup> |  | X |  |  |  |  |  |  |  | <1 |
| M6 | 7.59 | 230.63<sup>b</sup> |  |  | X |  |  |  |  |  |  | <1 |
| M7 | 7.70 | 472.26 |  |  |  | X |  |  |  |  |  | <1 |
| M8 | 7.79 | 418.25 |  |  |  |  | X |  |  |  |  | <1 |
| M9 | 7.80 | 417.18 |  |  |  | X | X | X | X |  |  | 2 |
| M10 | 7.91 | 342.12<sup>b</sup> | X | X |  |  |  |  | X |  |  | 5 |
| M11 | 8.47 | 230.63<sup>b</sup> |  |  | X |  |  |  |  |  |  | 10 |
| Parent | 8.58 | 474.27 |  |  |  |  |  |  |  |  |  | 53 |
| M12 | 8.90 | 463.21 |  |  |  |  | X | X | X | X | X | <1 |
| Total Rel. (%)<sup>c</sup> |  |  | 9 | 27 | 11 | 3 | 4 | 3 | 8 | 1 | 1 |  |

<sup>a</sup> See Figure 2 for structural assignments. <sup>b</sup> [M+2H]<sup>2+</sup>, z=2. <sup>c</sup> Sum of metabolites with the transformation, using <1 = 1.

### Hepatocyte Metabolism of Antiviral PZPs

Eight acetamide PZPs (**2a–f,h–i**) with IC_50_ ≤100 nM in the antiviral assay were selected for time course metabolic stability profiling in mouse and human and primary hepatocytes (Table 6). Unfortunately, all acetamide PZPs (**2a– f,h–i**) showed high intrinsic clearance in mouse hepatocytes, independent of their stability in liver microsomes (Table 2), suggesting that they were substrates for the Phase II GST enzymes in the hepatocytes. These in vitro metabolism data in mouse hepatocytes predicted that acetamide PZPs with potent antiviral activity would be likely to have rapid in vivo clearance in mice. In contrast, the compounds had much lower intrinsic clearance in human primary hepatocytes, with only **2h** and **2i** showing moderate and high rates of clearance, respectively.

**Table 6:**
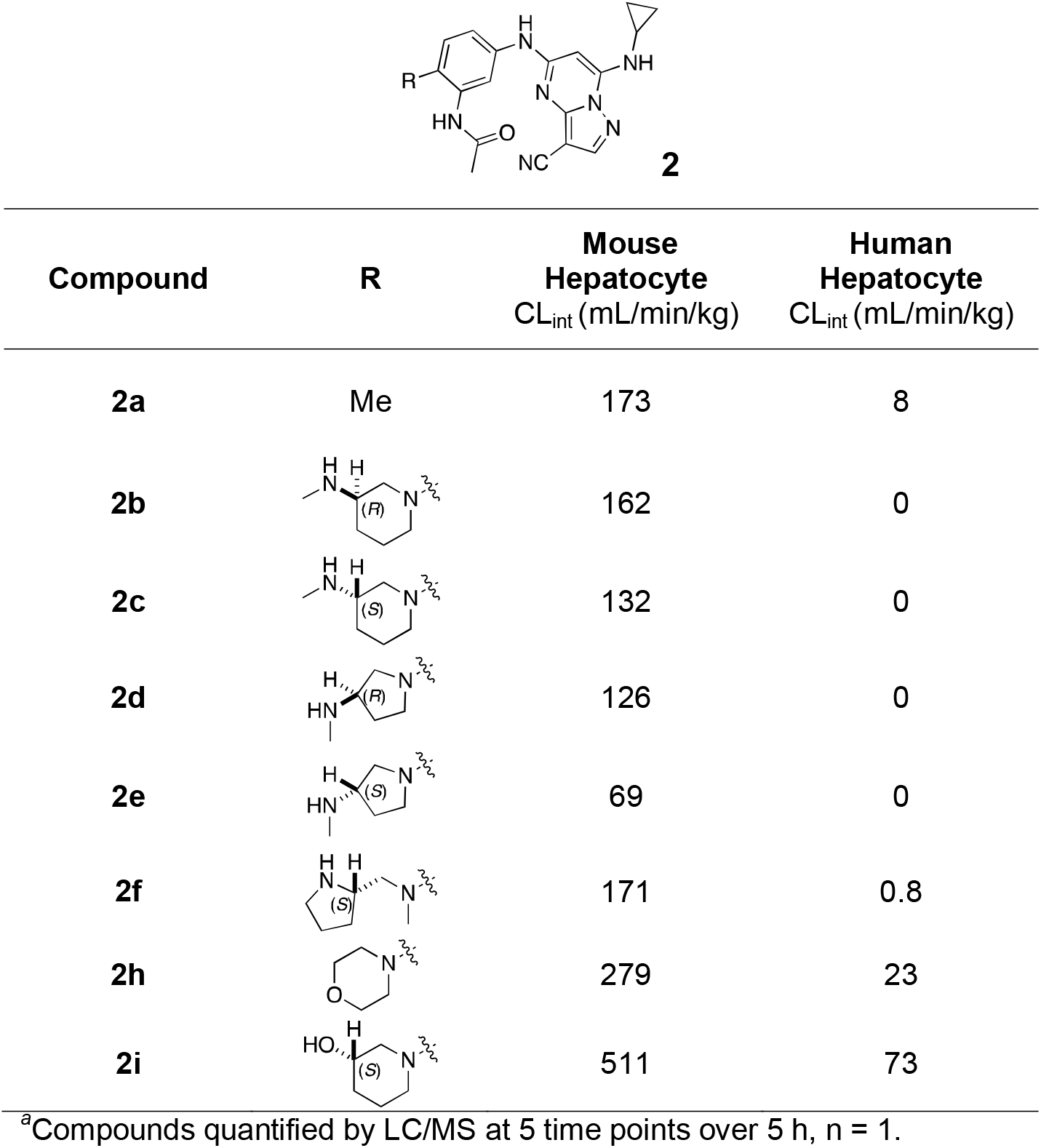
Mouse and human hepatocyte clearance***^a^***.

These results demonstrated that the metabolism of the PZPs in primary hepatocytes was a predominantly a species-specific issue, with intrinsic clearance always much higher in mouse than human hepatocytes. Although this was a promising result for the future development of PZPs as therapeutic drugs for use in humans, it still limited their utility as pharmacological tools for studies in mice.

### Inhibition of GSH Conjugation in Hepatocytes

Given our long-term objective to identify a potent CSNK2A inhibitor for use in an in vivo mouse COVID-19 efficacy model in vivo to validate the host kinase as a potential antiviral drug target, we opted to explore the effect of inhibition of the Phase II GST enzyme on intrinsic clearance. Ethacrynic acid (EA) is an FDA-approved loop diuretic that is also a potent covalent, but reversible, inhibitor of GST enzymes ^22^. EA has demonstrated potent inhibition of GST enzymes in perfused rat liver and human cancer cell lines,^22^ but although it has been used in human clinical studies to block GSH conjugation^23^ there are only a few reports of its use in rodents as a GST inhibitor.^24–26^ To determine if EA could improve the metabolic stability of a PZP in primary mouse hepatocytes we tested the effect of co-dosing on the stability of **2h**, which was one of the analogs with the highest intrinsic clearance (Table 6). The intrinsic clearance of **2h** in primary mouse hepatocytes was determined by measuring the decrease in the level of the parent compound over 2 h (Figure 3). Parallel sets of incubations were performed in the presence of increasing doses of EA from 0–400 µM. In the absence of EA intrinsic clearance was >250 mL/min/kg. However, in the presence of doses of EA from 10–50 µM the intrinsic clearance was decreased by >50% (Figure 3). At doses of EA above 100 µM, the intrinsic clearance of **2h** was further decreased to <50 mL/min/kg. The co-dosing mouse experiments in primary mouse hepatocytes demonstrated that EA was effective at increasing the metabolic stability of PZP **2h**, presumably by blocking its GSH conjugation.

**Figure 3.**
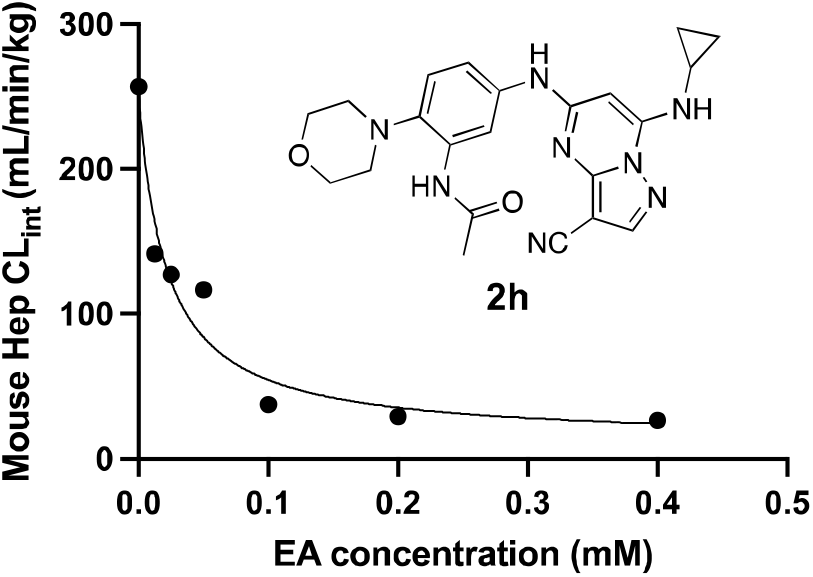
Mouse hepatocyte clearance of **2h** in the presence of GST inhibitor EA.

### Co-dosing of 2h with Inhibitors of Metabolic Enzymes In Vivo

To determine if inhibition of GSH conjugation could also be an effective strategy to increase the bioavailability of **2h** in vivo, we performed a series of co-dosing studies in mice of **2h** with either EA or 1-aminobenzotriaole (1-ABT), a widely used broad-spectrum inhibitor of mouse cytochrome P450s^27^ (Figure 4). When dosed alone to mice at 10 mg/kg i.p., **2h** was rapidly cleared from the circulation with a t_1/2_ ∼35 min and AUC ∼4000 h•nM. A 6-hour pre-treatment and co-dosing with 10 mg/kg i.p. EA resulted in a small increase in t_1/2_ and AUC. Increasing the dose to 30 mg/kg i.p. EA resulted in an additional increase in both parameters, demonstrating that inhibition of GST enzymes could reduce the intrinsic clearance of **2h**. The 30 mg/kg i.p. dose of EA increased the t_1/2_ 1.5-fold and AUC 1.8-fold. Unfortunately, we were unable to further increase the dose of EA to 100 mg/kg i.p. due to the observation of toxicity in mice on repeat dosing (Table S1). Since GSH conjugation of electron-rich aromatic groups requires a prior cytochrome P450 oxidation, we also evaluated the in vivo pharmacokinetics of **2h** in mice pretreated with 1-ABT for 2 h. 1-ABT was able to block the metabolism of **2h** in primary mouse hepatocytes (Figure S2). A decrease in the clearance of **2h** was observed in mice pretreated with 1-ABT at 100 mg/kg p.o. with an increase in t_1/2_ and AUC of 1.6-fold and 3.5-fold, respectively, compared to dosing of **2h** alone. The larger increase in AUC with 1-ABT compared to EA may be due to both inhibition of Phase I P450 metabolism and indirect inhibition of Phase II GSH conjugation.

**Figure 4.**
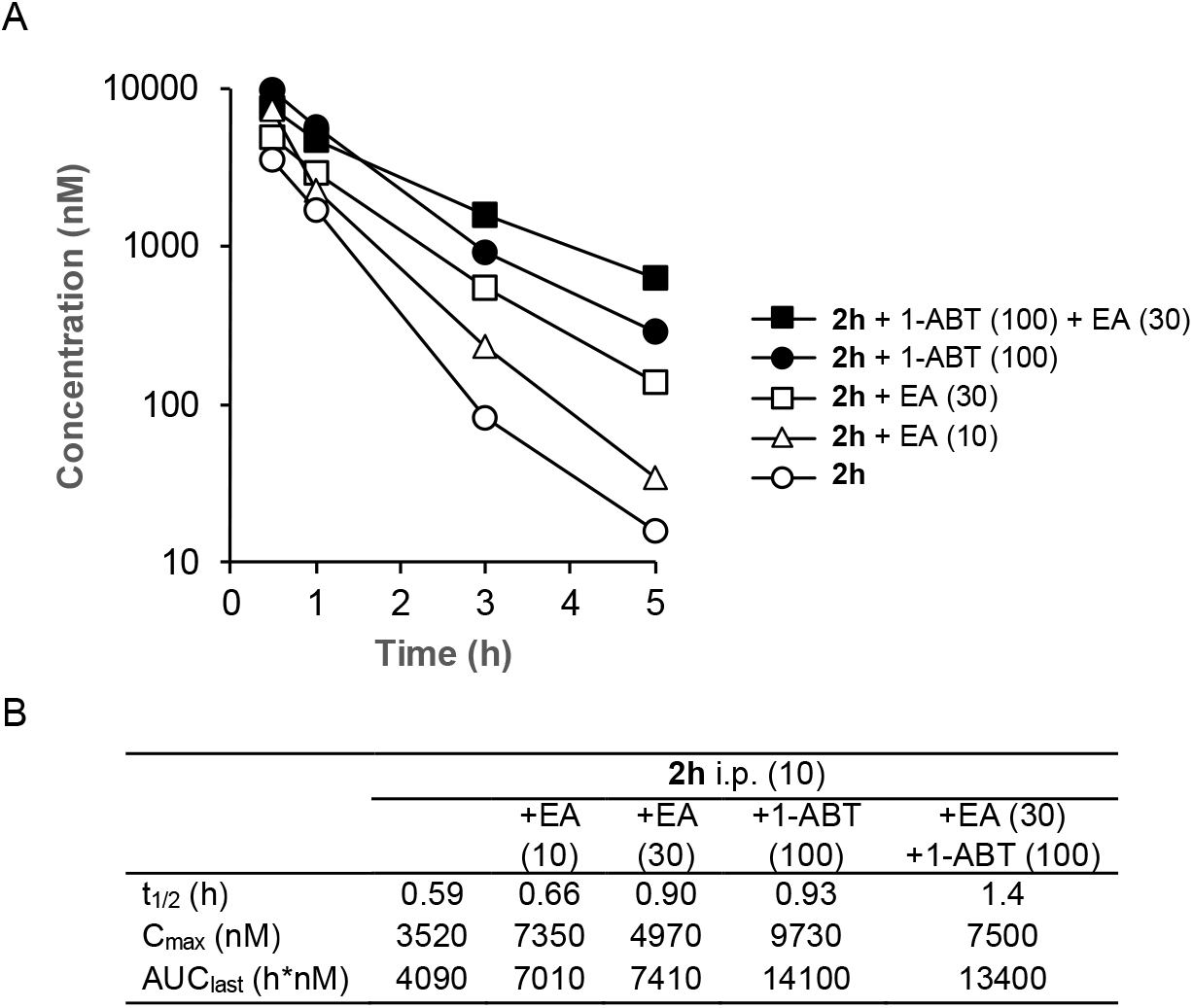
In vivo clearance of **2h** in mice in the presence of a GST inhibitor (EA) or a P450 inhibitor (1-ABT). A. Blood levels of **2h** over 5 h following a 10 mg/kg dose i.p. All data is the average from 3 mice. The dose of EA or 1-ABT is indicated in mg/kg. EA was administered i.p. as a 6 h pre-treatment and again co-dosed with **2h**. 1-ABT was administered p.o. as a 2 h pre-treatment. B. Pharmacokinetic parameters.

Importantly, the results demonstrate that it is possible to increase the circulating levels of the PZP CSNK2A inhibitor **2h** by co-dosing with either a GST inhibitor or a P450 inhibitor. Finally, to determine whether combined co-dosing with both EA and 1-ABT could further decrease the clearance of **2h**, we performed a study in the presence of both the GST and P450 inhibitors. A combination of 30 mg/kg i.p. EA (6-hour pre-treatment and co-dosed) with 100 mg/kg p.o. 1-ABT (2-hour pre-treatment) resulted in the highest levels of **2h** at the 3h and 5h time-points (Figure 4), with the t_1/2_ increased by 2.4-fold compared to dosing of **2h** alone.

## Discussion and Conclusions

The 3-cyano-7-cyclopropylamino-pyrazolo[1,5-a]pyrimidine (PZP) is the most potent and selective chemotype of ATP-competitive CSNK2A inhibitors,^7^ with fewer kinase off-targets than silmitasertib and related compounds.^28^ SGC-CK2-1 (**1a**)^8^ has been established as high-quality chemical probe for studying CSNK2 signaling in phenotypic assay and human-derived primary cells.^29–31^ However, the use of PZP-based CSNK2A inhibitors as chemical probes in rodent pharmacology models has been limited by their generally poor pharmacokinetic properties. Although Dowling and coworkers reported that **2a** had modest oral bioavailability in rats (F = 25%, C_max_ = ∼300 nM, t_1/2_ = 2.6 h), **2a** showed in vivo activity only at short time points after dosing.^9^ After extensive work on the series, Dowling concluded that an analog with an ethylenediamine substituent at the 4’-position could be used as a pharmacological tool by i.v. or i.p. dosing, despite its rapid clearance in rats.^10^ For the studies reported herein, we decided to focus on development of an in vivo pharmacological probe that would maintain high sustained exposure in mice following i.p. dosing, since this route of administration was preferred in many academic laboratories for preclinical target validation studies.^32^

Initially, we examined the potential of the 4’-substitued analogs of SGC-CK2-1 (**1a**) to improve solubility and reduce its rapid metabolism in mouse liver microsomes. Testing several of the previously prepared analogs of **1a**^6, 8^ confirmed prior observations^10^ of reduced metabolism of linear and cyclic diamine analogs with cLogP <0.5 (Table 1). After synthesizing many new 4’-substituted PZPs in the acetamide (**1**) and propionamide (**2**) series, the chiral 3-methylaminopipiperides **1h–I**, **2b–c** and 3-methylaminopyrollidines **2d–e** were found to have the best balance of reduced MLM metabolism and good cellular potency in the antiviral assay. However, despite their improved stability to Phase I metabolism **1i, 2c,** and **2e** showed rapid clearance in mice following i.v. or i.p. administration (Table 3).

After discovering that **1i**, although stable in MLM, was rapidly metabolized in mouse primary hepatocytes, we performed a metabolic ID study that identified Phase II GSH conjugation as a major metabolic transformation (Table 4 and Figure 2). This result suggested that Phase II metabolism by GST enzymes was a previously underappreciated metabolic liability of PZPs containing electron-rich di-and tri-aniline rings. Unfortunately, all potent antiviral PZPs showed rapid metabolism in mouse primary hepatocytes (Table 6). Although the metabolic instability may be species-specific, since it was not observed in human primary hepatocytes, the rapid clearance in mouse primary hepatocytes predicted that none of the potent antiviral PZP analogs would be suitable for use in the mouse-adapted COVID-19 model.

To explore ways to reduce the Phase II metabolism of a PZP analog, we investigated co-dosing with EA, a well characterized broad-spectrum covalent, but reversible, inhibitor of rodent and human GSTs.^22^ The studies were performed with the potent antiviral morpholine analog **2h**, which exhibited very high clearance in primary mouse hepatocytes. Co-dosing of EA with **2h** in primary mouse hepatocytes demonstrated that metabolism could be reduced in a dose dependent manner, with >90% reduction in intrinsic clearance at EA concentrations >100 µM (Figure 3).

Translating the use of EA as a GST inhibitor to an in vivo setting in mice was not straightforward. EA is an inhibitor of NKCC2, a Na-K-Cl transporter, and is approved for use in humans as a loop diuretic for treatment of high blood pressure and edema.^33^ At its commonly used 50 mg oral dose, EA does not cause significant drug-drug interactions that might result from GST inhibition. However, in rodents, no dose ranging studies to establish the lowest effective dose for GST inhibition have been reported. Although, doses of 10–20 mg/kg p.o. have been used to boost the activity of oncology drugs in mice.^24–26^ Our pharmacokinetic studies in mice revealed that the maximum tolerated dose of EA by i.p. dosing was only 30 mg/kg. Doses of 100 mg/kg and higher resulted in clinical symptoms including death (Table S1). Although the reason for the toxicity of EA by i.p. administration is unclear, the rapid onset suggests that it may be unrelated to inhibition liver metabolism enzymes. Nevertheless, we observed a reduction in the plasma clearance of **2h** at both the 10 and 30 mg/kg i.p. co-doses of EA. Furthermore, the clearance of **2h** was also reduced by a 100 mg/kg p.o. dose of the irreversible cytochrome P450 inhibitor 1-ABT, which would be expected to block direct Phase I and indirect Phase II metabolism. The most effective strategy to decrease the clearance of **2h** was the co-dosing protocol, using a combination of 30 mg/kg i.p. EA and 100 mg/kg p.o 1-ABT. With this approach the blood level of **2h** was boosted ∼40-fold at the 5 h time point.

In conclusion, our findings demonstrate that Phase II metabolism limits the in vivo exposure of PZP-based CSNK2A inhibitors in mice, which severely limits their utility as pharmacological probes despite their remarkable potency and selectivity in cells. Although the use of EA as a GST inhibitor provides an effective method to reduce the intrinsic clearance of the current molecules in mice, we are continuing to optimize the series to identify analogs with reduced potential for Phase II metabolism. Our goal is to find in vivo chemical probes for CSNK2A that can be dosed in mice without the additional complication of co-dosing with multiple inhibitors of drug metabolizing enzymes.

## Experimental Section

### NanoBRET Assay

Assays were run with a modified version of the previously published protocols.^6, 8^ HEK293 cells were cultured at 37 °C, 5% CO_2_ in Dulbecco’s modified Eagle medium (DMEM; Gibco) supplemented with 10% fetal bovine serum (VWR/Avantor). A transfection complex of DNA at 10 µg/mL was created, consisting of 9 µg/mL of carrier DNA (Promega) and 1 µg/mL of CSNK2A-NLuc fusion DNA in Opti-MEM without serum (Gibco). FuGENE HD (Promega) was added at 30 µl/mL to form a lipid:DNA complex. The solution was then mixed and incubated at room temperature for 20 min. The transition complex was mixed with a 20x volume of HEK293 cells at 20,000 cells per mL in DMEM/FBS and 100 µL per well was added to a 96-well plate that was incubated overnight at 37°C, 5% CO_2_. The following day, the media was removed via aspiration and replaced with 85 μL of Opti-MEM without phenol red. A total of 5 μL per well of 20x-NanoBRET Tracer K10 (Promega) at 10 μM for CSNK2A1 or 5 μM for CSNK2A2 in Tracer Dilution Buffer (Promega N291B) was added to all wells, except the “no tracer” control wells. Test compounds (10 mM in DMSO) were diluted 100x in Opti-MEM media to prepare stock solutions and evaluated at eleven concentrations. A total of 10 μL per well of the 10-fold test compound stock solutions (final assay concentration of 0.1% DMSO) were added. For “no compound” and “no tracer” control wells, DMSO in OptiMEM was added for a final concentration of 1.1% across all wells. 96-well plates containing cells with NanoBRET Tracer K10 and test compounds (100 µL total volume per well) were equilibrated (37°C / 5% CO_2_) for 2 h. The plates were cooled to room temperature for 15 min. NanoBRET NanoGlo substrate (Promega) at a ratio of 1:166 to Opti-MEM media in combination with extracellular NLuc Inhibitor (Promega) diluted 1:500 (10 μL of 30 mM stock per 5 mL Opti-MEM plus substrate) were combined to create a 3X stock solution. A total of 50 μL of the 3X substrate/extracellular NL inhibitor were added to each well. The plates were read within 30 min on a GloMax Discover luminometer (Promega) equipped with 450 nm BP filter (donor) and 600 nm LP filter (acceptor) using 0.3 s integration time. Raw milliBRET (mBRET) values were obtained by dividing the acceptor emission values (600 nm) by the donor emission values (450 nm) and multiplying by 1000. Averaged control values were used to represent complete inhibition (no tracer control: Opti-MEM + DMSO only) and no inhibition (tracer only control: no compound, Opti-MEM + DMSO + Tracer K10 only) and were plotted alongside the raw mBRET values. The data was first normalized and then fit using Sigmoidal, 4PL binding curve in Prism Software to determine IC_50_ values.

### MHV Assay

DBT cells were cultured at 37°C in Dulbecco’s modified Eagle medium (DMEM; Sigma) supplemented with 10% fetal bovine serum (Gibco) and penicillin and streptomycin (Sigma). DBT cells were plated in 96 well plates to be 80% confluent at the start of the assay. Test compounds were diluted to 15 µM in DMEM. Serial 4-fold dilutions were made in DMEM, providing a concentration range of 15 µM to 0.22 µM. Media was aspirated from the DBT cells and 100 μL of the diluted test compounds were added to the cells for 1 h at 37°C. After 1 h, MHV-nLuc^6^ was added at an MOI of 0.1 in 50 μL DMEM so that the final concentration of the first dilution of compound was 10 μM (T=0). After 10 h, the media was aspirated, and the cells were washed with PBS and lysed with passive lysis buffer (Promega) for 20 min at room temperature. Relative light units (RLUs) were measured using a luminometer (Promega; GloMax). Triplicate data was analyzed in Prism Graphpad to generate IC_50_ values.

### Kinetic Solubility

50 mL of phosphate buffered saline (PBS, Fisher, pH 7.4) was added to HPLC grade H_2_O (450 mL) for a total dilution factor of 1:10 and a final PBS concentration of 1X. 6 μL of the test compound as a 10 mM DMSO stock solutions was combined with the aqueous PBS solution (294 μL) for 50-fold dilution in a Millipore solubility filter plate with 0.45 μM polycarbonate filter membrane using a Hamilton Starlet liquid handler. The final DMSO concentration was 2.0% and maximum theoretical compound concentration was 200 μM. The filter plate was heat sealed for the duration of the 24 h incubation period. The sample was placed on a rotary shaker (200 RPM) for 24 h at ambient temperature (21.6–22.8°C) then vacuum filtered. All filtrates were injected into a chemiluminescence nitrogen detector for quantification. The equimolar nitrogen response of the detector was calibrated using standards which span the dynamic range of the instrument from 0.08 to 4500 μg/mL nitrogen. The filtrates were quantified with respect to this calibration curve. The calculated solubility values were corrected for background nitrogen present in the DMSO and the media used to prepare the samples.

### Mouse Liver Microsomal Stability

Compounds as 10 mM DMSO stock solutions were diluted to 2.5 mM with DMSO and again to 0.5 mM with MeCN to give a final solution containing 0.5 mM compound in 1:4 DMSO/MeCN. Liver microsomes from male CD-1 mice were sourced from Xenotech (Kansas City, KS), Lot No. 1710069. A reaction plate was prepared by adding 691.25 μL, pre-warmed (37°C) microsomal solution (0.63 mg/mL protein in 100 mM KPO_4_ with 1.3 mM EDTA) to an empty well of a 96-well plate and maintained at 37°C. The diluted 0.5 mM compound (8.75 μL) was added to the microsomal solution in the reaction plate and mixed thoroughly by repeated pipetting to give a final assay concentration of 5.0 μM compound. The resulting solutions were pre-incubated for 5 min at 37°C and then dispensed into T = 0 and incubation plates. For the T = 0 plates, an aliquot (160 μL) of each reaction solution was added to an empty well of a 96-well plate, as an exact replicate of the Reaction Plate. Cold (4°C)

MeOH (400 μL) was added to each well and mixed thoroughly by repeated pipetting. NADPH regeneration solution (40 μL) was added to each well and mixed thoroughly by repeated pipetting. For the T = 30-min Incubation Plate, NADPH (95 μL) was added to the remaining solution (microsomes + test compound) in each well in the previously prepared reaction plate to initiate the reaction. The plate was sealed and incubated at 37°C for 30 min. An aliquot (100 μL) was removed from each well at the desired time point and dispensed into a well of a 96-well plate. 200μL of cold (4°C) MeOH was added to quench the reaction. All plates were sealed, vortex and centrifuged at 3000 RPM, 4°C for 15 min, and the supernatants were transferred for analysis by LC-TOFMS. The supernatant (20 µL) was injected onto an AQUASIL C18 column and eluted using a fast-generic gradient program. TOFMS Data was acquired using Agilent 6538 Ultra High Accuracy TOF MS in extended dynamic range (m/z 100-1000) using generic MS conditions in positive mode. Following data acquisition, exact mass extraction and peak integration was performed using MassHunter Software (Agilent Technologies). The stability of the compound was calculated as the percent remaining of the unchanged parent at T=30 min relative to the peak area at T=0 min.

To determine CL_int_, aliquots of 50 µL were taken from the reaction solution at 0, 15, 30, 45 and 60 min. The reaction was stopped by the addition of 4 volumes of cold MeCN with IS (100 nM alprazolam, 200 nM imipramine, 200 nM labetalol and 2 μM ketoprofen). Samples were centrifuged at 3,220 g for 40 min and of 90 µL of the supernatant was mixed with 90 µL of ultra-pure H_2_O and then used for LC-MS/MS analysis. Peak areas were determined from extracted ion chromatograms and the slope value, k, was determined by linear regression of the natural logarithm of the remaining percentage of the parent drug vs. incubation time curve. The intrinsic clearance (CL_int_ in µL/min/mg) was calculated using the relationship CL_int_ = kV/N where V = incubation volume and N = amount of protein per well. To measure CL_int_ in the presence of microsomal and cytosolic enzymes, the experiment was repeated using liver S9 fractions microsomes from male CD-1 mice sourced from Xenotech (Kansas City, KS), Lot No. 1510255.

### Hepatocyte Stability

Human cryopreserved hepatocytes were supplied by BioIVT (lot QZW, 10 pooled donors). Mouse cryopreserved hepatocytes were supplied by BioIVT (lot ZPG, pooled male CD-1). Vials of cryopreserved hepatocytes were removed from storage and thawed in a 37°C water bath with gently shaking, and then the contents were poured into a 50 mL thawing medium conical tube. Vials were centrifuged at 100 g for 10 minutes at room temperature. Thawing medium was aspirated and hepatocytes were resuspended with serum-free incubation medium to yield ∼1.5 x 10^6^ cells/mL. Cell viability and density were counted using AO/PI fluorescence staining, and then cells were diluted with serum-free incubation medium to a working cell density of 0.5 x 10^6^ viable cells/mL. Aliquots of 198 μL hepatocytes were dispensed into each well of a 96-well non-coated plate. The plate was placed in an incubator for approximately 10 min. Aliquots of 2 μL of 100 μM test compound in duplicate and positive control were added into respective wells of the non-coated 96-well plate to start the reaction. The final concentration of test compound was 1 μM. The plate was placed in an incubator for the designed time points. 25 μL of contents were transferred and mixed with 6 volumes (150 μL) of cold MeCN with IS (100 nM alprazolam, 200 nM labetalol, 200 nM caffeine and 200 nM diclofenac) to terminate the reaction at time points of 0, 15, 30, 60, 90 and 120 minutes. Samples were centrifuged for 45 min at 3,220 g and aliquot of 100 µL of the supernatant was diluted with 100 µL of ultra-pure H_2_O, and the mixture was used for LC/MS/MS analysis. Peak areas were determined from extracted ion chromatograms and the slope value, k, was determined by linear regression of the natural logarithm of the remaining percentage of the parent drug vs. incubation time curve. The intrinsic clearance (CL_int_ in µL/min/10^6^ cells) was calculated using the relationship CL_int_ = kV/N where V = incubation volume (0.2 mL) and N = number of hepatocytes per well (0.1 x 10^6^ cells). Scaling factors to convert CL_int_ from µL/min/10^6^ cells to mL/min/kg were 2540 (human hepatocytes) and 11800 (mouse hepatocytes).

### Pharmacokinetics (PK)

Male CD-1 mice (6–8 weeks, 20–30 g) were dosed by intravenous (i.v.) or interperitoneal (i.p.) administration. For i.v. administration a single dose of the compound (1 mg/kg) was administered as a 5 mL/kg volume of a 0.2 mg/mL solution in NMP/Solutol/PEG-400/normal saline (v/v/v/v, 10:5:30:55) to three mice. For i.p. administration a single dose of the compound (10 mg/kg) was administered as a 10 mL/kg volume of a 1.0 mg/mL solution in 0.5% HPMC/0.2% Tween 80/99.3% H_2_O or DMSO/PEG-400/saline (v/v/v/v, 10:30:60) to three mice. The mice had free access to water and food. Blood (0.03 mL) was collected from dorsal metatarsal vein at 0.5, 1, 3, and 5 h timepoints (snapshot PK) or at 0.08, 0.25, 0.5, 1, 2, 4, 8, and 24 h (full PK). Blood from each sample was transferred into plastic microcentrifuge tubes containing EDTA-K2 and mixed well, then placed in a cold box prior to centrifugation. Blood samples were centrifuged at 4000 g for 5 min at 4°C to obtain plasma and then stored at −75°C prior to analysis. Concentrations of test compound in the plasma samples were determined using a Prominence LC-30AD, AB Sciex Triple Quan 5500 LC/MS/MS instrument fitted with a HALO 160A C18 column (2.7 µm, 2.1 x 50 mm) using a mobile phase of 5–95% MeCN in H_2_O with 0.1% formic acid. PK parameters were calculated from the mean plasma concentration versus time by a non-compartmental model using WinNonlin 8.3 (Phoenix) to determine C_max_, AUC_last_, t_1/2_, CL_int_, and *F*.

For co-dosing studies, **2h** (10 mg/kg) was administered i.p. as a 10 mL/kg volume of a 1.0 mg/mL solution in DMSO/PEG-400/saline (v/v/v/v, 10:30:60). EA (10 or 30 mg/kg) was administered i.p. as a 1 mL/kg volume of a 30 mg/mL solution in NMP/PEG-400/Water (v/v/v, 10:60:30) as a 6 h pretreatment and again at time of dosing of **2h**. 1-ABT (100 mg/kg) was administered p.o. as a 10 mL/kg volume of a 10 mg/mL solution in saline as a 2 h pretreatment. Blood (0.03 mL) was collected from dorsal metatarsal vein at 0.5, 1, 3, and 5 h timepoints and the level of **2h** in plasma determined by LC/MS/MS as described above. With this dosing protocol no adverse clinical observations were recorded during the study.

### General Synthetic Methods

All chemical reagents were commercially available except those whose synthesis is described below. All reaction mixtures and column eluents were monitored via analytical thin-layer chromatography (TLC) performed on pre-coated fluorescent silica gel plates, 200 μm with an F254 indicator; visualization was accomplished by UV light (254/365 nm). NMR spectra were obtained on a Bruker/AVANCE NEO 400MHz instrument. LCMS measurements were determined on Shimadzu LC-AB+LCMS2020, Shimadzu LC-AD+LCMS2020, Shimadzu LC-AD xR+LCMS2020, or Agilent 1200+ Infinitylab LC/MSD instruments. Purity was determined by HPLC measurement using a Shimadzu LC-20+LCMS-2020 instrument fitted with an Agilent PoroShell 120 EC-C18 column (45°C, 2.7 µm, 3.0*50 mm); 8 min chromatography run 0.037% TFA in water/MeCN (19:1) (solvent A), 0.018% TFA/MeCN (solvent B), gradient 0–60% (solvent B) over 6.0 min, held at 60% for 1.0 min, and returned to 0% (solvent B) for 1.0 min at a flow rate of 1.0 ml/min; 4 min chromatography run 0.037% TFA in water/MeCN (19:1) (solvent A), 0.018% TFA/MeCN (solvent B), gradient 10– 80% (solvent B) over 3.0 min, held at 80% for 0.5 min, and returned to 0% (solvent B) for 0.5 min at a flow rate of 1.0 ml/min; which indicated that all final compounds were >95% pure.

### General Procedure A

Synthesis of intermediate **y**. To a solution of *N*-(2-fluoro-5-nitrophenyl)propionamide (800 mg, 3.77 mmol) or *N*-(2-fluoro-5-nitrophenyl)acetamide (747 mg, 3.77 mmol) and an amine (3.77 mmol) in MeCN (8 mL) was added K_2_CO_3_ (1.56 g, 11.3 mmol) at 25°C. The mixture was stirred at 100°C for 10 h. The reaction mixture was concentrated in vacuo. Water (200 mL) was added, and the mixture was extracted with EtOAc (250 mL x 2). The combined organic phase was washed with brine (250 mL), dried over anhydrous Na_2_SO_4_, filtered, and concentrated in vacuo. To the resulting nitrobenzene (4.67 mmol) in MeOH (3 mL) was added 10% Pd/C (900 mg) under N_2_ atmosphere. The suspension was degassed and purged with H_2_ three times, and then stirred under H_2_ (15 psi) at 25°C for 10 h. The reaction mixture was filtered through a celite pad. The pad was washed with MeOH (8 mL x 2) and the combined filtrate concentrated in vacuo. The residue was purified by flash silica gel chromatography (eluent of 0–8%, MeOH/CH_2_Cl_2_) to yield intermediate **y**.

*(R)-N-(5-((3-cyano-7-(cyclopropylamino)pyrazolo[1,5-a]pyrimidin-5-yl)amino)-2-(3-(methylamino)piperidin-1-yl)phenyl)propionamide **(1h).*** Intermediate **y** was synthesized from tert-butyl (*R*)-methyl(piperidin-3-yl)carbamate and *N*-(2-fluoro-5-nitrophenyl)propionamide by Procedure A. To a solution of 5-chloro-7-(cyclopropylamino)pyrazolo[1,5-a]pyrimidine-3-carbonitrile^6^ (100 mg, 0.43 mmol) and intermediate **y** (170mg, 0.43 mmol) in dioxane (3 mL) was added Cs_2_CO_3_ (418 mg, 1.28 mmol), BINAP (40 mg, 0.06 mmol) and Pd(OAc)_2_ (14.0 mg, 0.06 mmol) at 25°C. The mixture was degassed and purged with N_2_ and then heated in a microwave reactor at 130°C for 0.5 h. The reaction mixture was concentrated in vacuo and the residue was purified by flash silica gel chromatography (eluent of 0–4%, MeOH/CH_2_Cl_2_). To the product in CH_2_Cl_2_ (5 mL) was added TFA (7 mL), and the mixture was stirred at 25°C for 2 h. The reaction mixture was concentrated in vacuo and the residue was purified by prep-HPLC (column: Xtimate C18 150*40 mm*10 μm; mobile phase: [water (HCO_2_H)-MeCN]; B%: 15–45%, 10 min). Compound **1h** (66.3 mg, 11.3%) was obtained as a white solid. ^1^H NMR (400 MHz, DMSO-*d_6_*) δ 9.65 (s, 1H), 9.33 (br s, 1H), 8.44 (s, 1H), 8.33 (s, 1H), 8.19 (s, 1H), 8.04 (d, *J* = 2.4 Hz, 1H), 7.82 (d, *J* = 7.2 Hz, 1H), 7.09 (d, *J* = 8.4 Hz, 1H), 6.03 (s, 1H), 3.13 (br s, 1H), 3.05 (d, *J* = 10.4 Hz, 1H), 2.89 -2.80 (m, 2H), 2.60-2.58 (m, 2H), 2.52-2.51 (m, 1H), 2.48-2.44 (m, 4H), 1.92-1.91 (m, 1H), 1.75 (s, 2H), 1.62-1.60 (m, 1H), 1.12 (t, *J* = 7.6 Hz, 3H), 0.80-0.79 (m, 2H), 0.72-0.70 (m, 2H). ^13^C NMR (101 MHz, DMSO-*d_6_*) δ 172.04, 165.85, 156.77, 150.78, 148.07, 144.88, 137.77, 136.07, 132.78, 119.97, 114.74, 76.13, 54.06, 52.95, 38.69, 30.66, 29.41, 23.17, 9.79, 6.39. HPLC R_t_ = 3.656 min in 8 min chromatography, purity 99.6%. LCMS R_t_ = 1.925 min in 4 min chromatography, purity 98.0%, MS ESI calcd. for 473.27 [M+H]^+^ 474.27, found 474.4.

*(S)-N-(5-((3-cyano-7-(cyclopropylamino)pyrazolo[1,5-a]pyrimidin-5-yl)amino)-2-(3-(methylamino)piperidin-1-yl)phenyl)propionamide (**1i**).* Intermediate **y** was synthesized from tert-butyl (*S*)-methyl(piperidin-3-yl)carbamate and *N*-(2-fluoro-5-nitrophenyl)propionamide by Procedure A. To a solution of 5-chloro-7-(cyclopropylamino)pyrazolo[1,5-a]pyrimidine-3-carbonitrile^6^ (112 mg, 0.48 mmol) and intermediate **y** (200 mg, 0.53 mmol) in dioxane (3 mL) was added Cs_2_CO_3_ (502 mg, 1.53 mmol), BINAP (48 mg, 0.07 mmol) and Pd(OAc)_2_ (16.8 mg, 0.07 mmol) at 25°C. The mixture was degassed and purged with N_2_ and heated in a microwave reactor at 130°C for 0.5 h. The reaction mixture was concentrated in vacuo and the residue was purified by flash silica gel chromatography (eluent of 0–4%, MeOH/CH_2_Cl_2_). To the product in CH_2_Cl_2_ (5 mL) was added TFA (7 mL), and the mixture was stirred at 25°C for 2 h. The reaction mixture was concentrated in vacuo and the residue was purified by prep-HPLC (column: Xtimate C18 150*40 mm*10 μm; mobile phase: [water (HCO_2_H)-MeCN]; B%: 0–38%, 36min). Compound **1i** (115 mg, 37.0%) was obtained as a yellow solid. ^1^H NMR (400 MHz, DMSO-*d_6_*) δ 9.65 (s, 1H), 9.09 (s, 1H), 8.43-8.36 (m, 1H), 8.34 (s, 1H), 8.20 (s, 1H), 8.07-8.01 (m, 1H), 7.89-7.78 (m, 1H), 7.12 (d, *J* = 8.8 Hz, 1H), 6.03 (s, 1H), 3.24 (s, 1H), 3.12-3.05 (m, 1H), 2.93-2.84 (m, 1H), 2.83-2.75 (m, 1H), 2.68 -2.54 (m, 5H), 2.48-2.41 (m, 2H), 1.95-1.60 (m, 4H), 1.13 (t, *J* = 7.6 Hz, 3H), 0.84-0.66 (m, 4H). ^13^C NMR (101 MHz, DMSO-*d_6_*) δ 172.16, 165.58, 156.97, 150.97, 148.26, 145.06, 137.96, 136.25, 132.97, 120.21, 115.33, 114.91, 76.32, 54.47, 53.83, 53.02, 31.24, 29.65, 23.36, 21.77, 9.95, 6.57. HPLC R_t_ = 3.612 min in 8 min chromatography, purity 99.6%. LCMS R_t_ = 1.567 min in 4 min chromatography, purity 100.0%, MS ESI calcd. for 473.3 [M+H]^+^ 474.3, found 474.5.

*(R)-N-(5-((3-cyano-7-(cyclopropylamino)pyrazolo[1,5-a]pyrimidin-5-yl)amino)-2-(3-(methylamino)piperidin-1-yl)phenyl)acetamide (**2b**).* Intermediate **y** was synthesized from tert-butyl (R)-methyl(piperidin-3-yl)carbamate and N-(2-fluoro-5-nitrophenyl)acetamide by Procedure A. To a solution of 5-chloro-7-(cyclopropylamino)pyrazolo[1,5-a]pyrimidine-3-carbonitrile^6^ (112 mg, 0.48 mmol) and intermediate **y** (200 mg, 0.55 mmol) in dioxane (3 mL) was added Cs_2_CO_3_ (541 mg, 1.65 mmol), BINAP (51 mg, 0.08 mmol) and Pd(OAc)_2_ (17.9 mg, 0.08 mmol) at 25°C. The mixture was degassed and purged with N_2_ and heated in a microwave reactor at 130°C for 0.5 h. The reaction mixture was concentrated in vacuo and the residue was purified by flash silica gel chromatography (eluent of 0–32%, EtOAc/petroleum ether) to give the product. To the product in CH_2_Cl_2_ (5 mL) was added TFA (7 mL), and the mixture was stirred at 25°C for 2 h. The reaction mixture was concentrated in vacuo and the residue was purified by prep-HPLC (column: Phenomenex C18 75*30 mm*3 μm; mobile phase: [water (HCO_2_H)-MeCN]; B%: 2– 32%, 24 min). Compound **2b** (32.8 mg, 27.4%) was obtained as a white solid. ^1^H NMR (400 MHz, DMSO-d_6_) δ 9.63 (s, 1H), 9.41 (s, 1H), 8.41 (s, 1H), 8.32 (s, 1H), 8.16 (s, 1H), 7.99 (d, *J* = 1.6 Hz, 1H), 7.83 (d, *J* = 6.4 Hz, 1H), 7.10 (d, *J* = 8.8 Hz, 1H), 6.03 (s, 1H), 3.22 (s, 1H), 3.06 (d, *J* = 11.2 Hz, 1H), 2.93 (d, *J* = 8.4 Hz, 1H), 2.84 (d, *J* = 11.2 Hz, 1H), 2.58 (s, 2H), 2.53 (s, 3H), 2.18 (s, 3H), 1.95 (s, 1H), 1.79 (s, 2H), 1.67-1.56 (m, 1H), 0.80 (d, *J* = 6.8 Hz, 2H), 0.70 (d, *J* = 2.8 Hz, 2H). ^13^C NMR (101 MHz, DMSO-*d*_6_) δ 168.62, 163.65, 156.97, 150.98, 148.26, 145.08, 137.91, 136.26, 133.03, 120.20, 115.42, 114.96, 76.32, 54.08, 53.19, 40.15, 39.99, 39.78, 24.09, 23.36, 6.59. HPLC R_t_ = 3.321 min in 8 min chromatography, purity 99.9%. LCMS R_t_ = 1.551 min in 4 min chromatography, purity 99.8%, MS ESI calcd. for 459.25 [M+H]^+^ 460.25, found 460.4.

*(S)-N-(5-((3-cyano-7-(cyclopropylamino)pyrazolo[1,5-a]pyrimidin-5-yl)amino)-2-(3-(methylamino)piperidin-1-yl)phenyl)acetamide (**2c**).* Intermediate **y** was synthesized from tert-butyl (*S*)-methyl(piperidin-3-yl)carbamate and *N*-(2-fluoro-5-nitrophenyl)acetamide by Procedure A. To a solution of 5-chloro-7-(cyclopropylamino)pyrazolo[1,5-a]pyrimidine-3-carbonitrile^6^ (237 mg, 1.02 mmol) and intermediate **y** (460 mg, 1.27 mmol) in dioxane (6 mL) was added Cs_2_CO_3_ (1.0 g, 1.65 mmol), BINAP (94.8 mg, 0.14 mmol) and Pd(OAc)_2_ (33.2 mg, 0.14 mmol) at 25°C. The mixture was degassed and purged with N_2_ and heated in a microwave reactor at 130°C for 0.5 h. The reaction mixture was concentrated in vacuo and the residue was purified by flash silica gel chromatography (eluent of 0–3%, MeOH/CH_2_Cl_2_). To the product in CH_2_Cl_2_ (5 mL) was added TFA (7 mL), and the mixture was stirred at 25°C for 2 h. The reaction mixture was concentrated in vacuo and the residue was purified by prep-HPLC (column: Xtimate C18 100*30 mm*10 μm; mobile phase: [water (HCO_2_H)-MeCN]; B%: 30%-50%, 10min). Compound **2c** (58 mg, 27.4%) was obtained as a white solid. ^1^H NMR (400 MHz, DMSO-*d_6_*) δ 9.64 (s, 1H), 9.38-9.21 (m, 1H), 8.39-8.30 (m, 1H), 8.17 (s, 1H), 7.97 (s, 1H), 7.89-7.77 (m, 1H), 7.10 (d, *J* = 8.8 Hz, 1H), 6.03 (s, 1H), 3.11-3.03 (m, 2H), 3.02-2.90 (m, 2H), 2.89-2.82 (m, 1H), 2.62-2.53 (m, 5H), 2.20 (s, 3H), 2.06-1.94 (m, 1H), 1.89-1.72 (m, 2H), 1.67-1.58 (m, 1H), 0.84-0.76 (m, 2H), 0.73-0.66 (m, 2H). ^13^C NMR (101 MHz, DMSO*-d_6_*) δ 168.66, 162.94, 156.95, 150.96, 148.25, 145.07, 137.91, 136.26, 132.85, 120.17, 115.48, 114.94, 113.94, 76.32, 54.13, 53.14, 39.99, 30.59, 24.20, 23.36, 6.58. HPLC R_t_ = 3.324 min in 8 min chromatography, purity 99.2%. LCMS R_t_ = 1.549 min in 4 min chromatography, purity 99.6%, MS ESI calcd. for 459.25 [M+H]^+^ 460.25, found 460.4.

*(R)-N-(5-((3-cyano-7-(cyclopropylamino)pyrazolo[1,5-a]pyrimidin-5-yl)amino)-2-(3-(methylamino)pyrrolidin-1-yl)phenyl)acetamide (**2d**).* Intermediate **y** was synthesized from tert-butyl (*R*)-methyl(pyrrolidin-3-yl)carbamate and *N*-(2-fluoro-5-nitrophenyl)acetamide by Procedure A. To a solution of 5-chloro-7-(cyclopropylamino)pyrazolo[1,5-a]pyrimidine-3-carbonitrile^6^ (134 mg, 0.57 mmol) and intermediate **y** (250 mg, 0.72 mmol) in dioxane (3 mL) was added Cs_2_CO_3_ (0.5 g, 0.82 mmol), BINAP (47.4 mg, 0.07 mmol) and Pd(OAc)_2_ (16.6 mg, 0.07 mmol) at 25°C. The mixture was degassed and purged with N_2_ and heated in a microwave reactor at 130°C for 0.5 h. The reaction mixture was concentrated in vacuo and the residue was purified by prep-HPLC (column: Waters Torus 2-PIC 150*19 mm*5 μm; mobile phase: [Heptane-EtOH (0.1% NH_3_/H_2_O)]; B%: 10–50%, 5min) to give the product as a yellow solid. To the product in CH_2_Cl_2_ (5 mL) was added TFA (7 mL), and the mixture was stirred at 25°C for 2 h. The reaction mixture was concentrated in vacuo and the residue was purified by prep-HPLC (column: Xtimate C18 100*30 mm*10 μm; mobile phase: [water (HCO_2_H)-MeCN]; B%: 25–45%, 10 min). Compound **2d** (20 mg, 30.2%) was obtained as a white solid. ^1^H NMR (400 MHz, MeOD-*d*_4_) δ 9.56 (s, 1H), 9.30 (s, 1H), 8.46-8.28 (m, 2H), 8.16 (s, 1H), 7.93-7.60 (m, 2H), 7.02 (d, *J* = 8.4 Hz, 1H), 5.99 (s, 1H), 3.55-3.50 (m, 1H), 3.34-3.09 (m, 4H), 2.95-2.84 (m, 1H), 2.61-2.51 (m, 4H), 2.26-2.06 (m, 4H), 1.9-1.86 (m, 1H), 0.86-0.76 (m, 2H), 0.75-0.66 (m, 2H). ^13^C NMR (101 MHz, DMSO-*d*_6_) δ 168.49, 157.01, 151.02, 148.19, 145.00, 134.54, 130.86, 118.11, 116.30, 114.95, 76.17, 57.99, 54.44, 49.20, 39.99, 23.89, 23.32, 6.55. HPLC R_t_ = 3.063 min in 8 min chromatography, purity 99.8%. LCMS R_t_ = 1.422 min in 4 min chromatography, purity 99.9%, MS ESI calcd. for 445.23 [M+H]^+^ 446.23, found 446.4.

*(S)-N-(5-((3-cyano-7-(cyclopropylamino)pyrazolo[1,5-a]pyrimidin-5-yl)amino)-2-(3-(methylamino)pyrrolidin-1-yl)phenyl)acetamide (**2e**).* Intermediate **y** was synthesized from tert-butyl (*S*)-methyl(pyrrolidin-3-yl)carbamate and *N*-(2-fluoro-5-nitrophenyl)acetamide by Procedure A. To a solution of 5-chloro-7-(cyclopropylamino)pyrazolo[1,5-a]pyrimidine-3-carbonitrile^6^ (520 mg, 2.23 mmol) and intermediate **y** (930 mg, 2.67 mmol) in dioxane (15 mL) was added Cs_2_CO_3_ (2.2 g, 6.7 mmol), BINAP (207 mg, 0.31 mmol) and Pd(OAc)_2_ (72.7 mg, 0.31 mmol) at 25°C. The mixture was degassed and purged with N_2_ and heated in a microwave reactor at 130°C for 0.5 h. The reaction mixture was concentrated in vacuo and the residue was purified flash silica gel chromatography (eluent of 0–51%, EtOAc/petroleum ether) to give the product as a gray solid. To the product in CH_2_Cl_2_ (5 mL) was added TFA (7 mL), and the mixture was stirred at 25°C for 2 h. The reaction mixture was concentrated in vacuo and the residue was purified by prep-HPLC (column: Welch Xtimate C18 150*30 mm*5 μm; mobile phase: [water (HCO_2_H)-MeCN]; B%: 0–90%, 14 min). Compound **2e** (30.6 mg, 16.2%) was obtained as a light-yellow solid. ^1^H NMR (400 MHz, DMSO-*d*_6_) δ 9.59 (s, 1H), 9.42 (s, 1H), 8.41 (s, 1H), 8.33 (s, 1H), 8.18 (s, 1H), 7.86 (s, 1H), 7.76 (s, 1H), 7.03 (d, *J* = 8.8 Hz, 1H), 6.00 (s, 1H), 3.56 (s, 1H), 3.30 (d, *J* = 4.4 Hz, 1H), 3.21 - 3.16 (m, 1H), 3.16 - 3.11 (m, 1H), 2.82 (q, *J* = 8.0 Hz, 1H), 2.57 (s, 1H), 2.52-2.51 (m, 3H), 2.26-2.17 (m, 1H), 2.14 (s, 3H), 1.97 (s, 1H), 0.84-0.76 (m, 2H), 0.72-0.66 (m, 2H). ^13^C NMR (101 MHz, DMSO-*d*_6_) δ 168.51, 165.64, 157.00, 151.01, 148.19, 145.00, 134.62, 131.13, 118.19, 116.20, 115.48, 114.95, 76.16, 76.04, 57.88, 54.30, 49.20, 31.64, 28.59, 23.90, 23.32, 6.55. HPLC R_t_ = 3.080 min in 8 min chromatography, purity 99.4%. LCMS R_t_ = 1.420 min in 4 min chromatography, purity 99.7%, MS ESI calcd. For 445.23 [M+H]^+^ 446.23, found 446.4.

*(S)-N-(5-((3-cyano-7-(cyclopropylamino)pyrazolo[1,5-a]pyrimidin-5-yl)amino)-2-(methyl(pyrrolidin-2-ylmethyl)amino)phenyl)acetamide (**2f**).* Intermediate **y** was synthesized from tert-butyl (*S*)-2-((methylamino)methyl)pyrrolidine-1-carboxylate and *N*-(2-fluoro-5-nitrophenyl)acetamide by Procedure A. To a solution of 5-chloro-7-(cyclopropylamino)pyrazolo[1,5-a]pyrimidine-3-carbonitrile^6^ (52.0 mg, 0.220 mmol) and intermediate **y** (100 mg, 0.280 mmol) in dioxane (2 mL) was added Cs_2_CO_3_ (209 mg, 0.64 mmol), BINAP (20 mg, 0.03 mmol) and Pd(OAc)_2_ (7.0 mg, 0.03 mmol) at 25°C. The mixture was degassed and purged with N_2_ and heated in a microwave reactor at 130°C for 0.5 h. The reaction mixture was concentrated in vacuo and the residue was purified flash silica gel chromatography (eluent of 0–2%, MeOH/CH_2_Cl_2_) to give the product as a brown solid. To the product in CH_2_Cl_2_ (5 mL) was added TFA (7 mL), and the mixture was stirred at 25°C for 2 h. The reaction mixture was concentrated in vacuo and the residue was purified by prep-HPLC (column: Xtimate C18 150*40 mm*10 μm; mobile phase: [water (NH_4_HCO_3_)-MeCN]; B%: 22– 62%, 25 min). Compound **2f** (25.0 mg, 30.3%) was obtained as a yellow solid. ^1^H NMR (400 MHz, DMSO-*d*_6_) δ 10.76-10.36 (m, 1H), 9.61 (s, 1H), 8.37-8.30 (m, 2H), 8.16 (s, 1H), 7.89-7.77 (m, 1H), 7.17 (d, *J* = 8.8 Hz, 1H), 6.05 (s, 1H), 3.48-3.39 (m, 1H), 3.36-3.33 (m, 2H), 3.01-2.91 (m, 1H), 2.85-2.74 (m, 1H), 2.66 (s, 3H), 2.61-2.54 (m, 1H), 2.14 (s, 3H), 1.92-1.58 (m, 3H), 1.30-1.17 (m, 1H), 0.83-0.75 (m, 2H), 0.74-0.67 (m, 2H). ^13^C NMR (101 MHz, DMSO-*d*_6_) δ 168.17, 156.98, 150.95, 148.20, 145.02, 137.56, 135.98, 132.98, 120.22, 114.90, 76.25, 65.21, 59.77, 51.82, 31.69, 24.21, 23.32, 22.38, 6.55. HPLC R_t_ = 3.426 min in 8 min chromatography, purity 99.4%. LCMS R_t_ = 2.058 min in 4 min chromatography, purity 95.5%, MS ESI calcd. For 445.23 [M+H]^+^ 446.23, found 460.3.

*N-(5-((3-cyano-7-(cyclopropylamino)pyrazolo[1,5-a]pyrimidin-5-yl)amino)-2-(piperazin-1-yl)phenyl)acetamide (**2g**).* Intermediate **y** was synthesized from tert-butyl piperazine-1-carboxylate and *N*-(2-fluoro-5-nitrophenyl)acetamide by Procedure A. To a solution of 5-chloro-7-(cyclopropylamino)pyrazolo[1,5-a]pyrimidine-3-carbonitrile^6^ (302 mg, 1.29 mmol) and intermediate **y** (433 mg, 1.29 mmol) in dioxane (9 mL) was added Cs_2_CO_3_ (1.25 g, 3.84 mmol), BINAP (120 mg, 0.18 mmol) and Pd(OAc)_2_ (42.0 mg, 0.18 mmol) at 25°C. The mixture was degassed and purged with N_2_ and heated in a microwave reactor at 130°C for 0.5 h. The reaction mixture was concentrated in vacuo and the residue was purified by flash silica gel chromatography (eluent of 30–70%, EtOAc/petroleum ether) to give the product as a brown solid. To the product in CH_2_Cl_2_ (5 mL) was added TFA (7 mL), and the mixture was stirred at 25°C for 2 h. The reaction mixture was concentrated in vacuo and the residue was purified by prep-HPLC (column: Xtimate C18 100*30 mm*10 μm; mobile phase: [water (HCO_2_H)-MeCN]; B%: 5–96%, 10 min]; B%: 4–34%, 10 min). Compound **2g** (178.4 mg, 27.4%) was obtained as a light yellow solid. ^1^H NMR (400 MHz, DMSO-*d*_6_) δ 9.68 (s, 1H), 8.92 (s, 1H), 8.33 (m, 1H), 8.48-8.31 (m, 1H), 8.12-7.99 (m, 1H), 7.89 (d, *J* = 6.8 Hz, 1H), 7.15 (d, *J* = 9.2 Hz, 1H), 6.03 (s, 1H), 3.35-3.22 (m, 4H), 3.02-2.88 (m, 4H), 2.63-2.55 (m, 1H), 2.16 (s, 3H), 0.87-0.76 (m, 2H), 0.73-0.66 (m, 2H). ^13^C NMR (101 MHz, DMSO-*d*_6_) δ 168.53, 156.91, 150.90, 148.24, 145.06, 136.92, 136.49, 133.12, 120.45, 115.14, 114.90, 112.84, 76.38, 49.29, 43.46, 39.99, 24.25, 23.34, 6.56. HPLC R_t_ = 1.599 min in 6 min chromatography, purity 95.6%. LCMS R_t_ = 1.003 min in 4 min chromatography, purity 95.9%, MS ESI calcd. for 431.22 [M+H]^+^ 432.22, found 432.2.

*(S)-N-(5-((3-cyano-7-(cyclopropylamino)pyrazolo[1,5-a]pyrimidin-5-yl)amino)-2-(3-hydroxypiperidin-1-yl)phenyl)acetamide (**2i**).* Intermediate **y** was synthesized from (*S*)-piperidin-3-ol hydrochloride and *N*-(2-fluoro-5-nitrophenyl)acetamide by Procedure A. To a solution of 5-chloro-7-(cyclopropylamino)pyrazolo[1,5-a]pyrimidine-3-carbonitrile^6^ (75 mg, 0.32 mmol) and intermediate **y** (100 mg, 0.40 mmol) in dioxane (3 mL) was added Cs_2_CO_3_ (388 mg, 1.19 mmol), BINAP (37.2 mg, 0.05 mmol) and Pd(OAc)_2_ (12.7 mg, 0.05 mmol) at 25°C. The mixture was degassed, purged with N_2_, and heated in a microwave reactor at 130°C for 0.5 h. The reaction mixture was concentrated in vacuo and the residue was purified by prep-HPLC (column: Xtimate C18 150*30 mm*5 μm; mobile phase: [water (NH_4_HCO_3_)-MeCN]; B%: 20–60%, 25 min) to give compound **2i** (25.0 mg, 13.8%) as a white solid. ^1^H NMR (400 MHz, DMSO-*d*_6_) δ 9.61 (s, 1H), 9.04 (s, 1H), 8.33 (s, 1H), 8.19-8.04 (m, 2H), 7.88-7.83 (m, 1H), 7.08 (d, *J* = 8.4 Hz, 1H), 6.03 (s, 1H), 5.09-4.87 (m, 1H), 3.80 (s, 1H), 2.86-2.76 (m, 2H), 2.75-2.67 (m, 1H), 2.65-2.54 (m, 2H), 2.13 (s, 3H), 1.95-1.81 (m, 1H), 1.77-1.55 (m, 2H), 1.54-1.42 (m, 1H), 0.83-0.75 (m, 2H), 0.74-0.67 (m, 2H). ^13^C NMR (101 MHz, DMSO-*d*_6_) δ 168.74, 157.10, 151.04, 148.24, 145.08, 137.49, 136.04, 134.09, 120.57, 115.02, 114.57, 111.52, 76.29, 48.67, 45.67, 28.83, 24.17, 23.38, 6.61. HPLC R_t_ = 3.771 min in 8 min chromatography, purity 99.0%. LCMS R_t_ = 2.050 min in 4 min chromatography, purity 98.3%, MS ESI calcd. for 446.22 [M+H]^+^ 447.22, found 447.3.

*N-(5-((3-cyano-7-(cyclopropylamino)pyrazolo[1,5-a]pyrimidin-5-yl)amino)-2-(methyl(oxetan-3-yl)amino)phenyl)acetamide (**2j**).* Intermediate **y** was synthesized from *N-*methyloxetan-3-amine and *N*-(2-fluoro-5-nitrophenyl)acetamide by Procedure A. To a solution of 5-chloro-7-(cyclopropylamino)pyrazolo[1,5-a]pyrimidine-3-carbonitrile^6^ (96.0 mg, 0.410 mmol) and intermediate **y** (120 mg, 0.510 mmol) in dioxane (3 mL) was added Cs_2_CO_3_ (398 mg, 1.21 mmol), BINAP (38.1 mg, 0.06 mmol) and Pd(OAc)_2_ (13.3 mg, 0.06 mmol) at 25°C. The mixture was degassed, purged with N_2_, and heated in a microwave reactor at 130°C for 0.5 h. The reaction mixture was concentrated in vacuo and the residue was purified by prep-HPLC (column: Xtimate C18 150 * 40 mm * 10 μm; mobile phase: [water (NH_4_HCO_3_)-MeCN]; B%: 18– 58%, 25 min). Compound **2j** (4.70 mg, 5.88 %) was obtained as a yellow solid. ^1^H NMR (400 MHz, DMSO-*d*_6_) δ 9.65 (s, 1H), 9.06 (s, 1H), 8.34 (s, 1H), 8.19 (s, 1H), 8.10 (s, 1H), 7.86 (d, *J* = 8.0 Hz, 1H), 6.97 (d, *J* = 8.8 Hz, 1H), 6.03 (s, 1H), 4.64-4.56 (m, 2H), 4.44-4.37 (m, 2H), 4.38-4.30 (m, 1H), 2.62-2.54 (m, 1H), 2.46 (s, 3H), 2.18 (s, 3H), 0.85-0.76 (m, 2H), 0.75-0.67 (m, 2H). HPLC R_t_ = 4.106 min in 8 min chromatography, purity 99.9%. LCMS R_t_ = 2.487 min in 4 min chromatography, purity 97.9 %, MS ESI calcd. for 432.2, [M+H]^+^ 433.2, found 433.2.

*(S)-N-(5-((3-cyano-7-(cyclopropylamino)pyrazolo[1,5-a]pyrimidin-5-yl)amino)-2-(methyl(tetrahydrofuran-3-yl)amino)phenyl)acetamide (**2k**).* Intermediate **y** was synthesized from (*S*)-N-methyltetrahydrofuran-3-amine hydrochloride and *N*-(2-fluoro-5-nitrophenyl)acetamide by Procedure A. To a solution of 5-chloro-7-(cyclopropylamino)pyrazolo[1,5-a]pyrimidine-3-carbonitrile^6^ (84.3 mg, 0.361 mmol) and intermediate **y** (90.0 mg, 0.361 mmol) in dioxane (2 mL) was added Cs_2_CO_3_ (470 mg, 1.44 mmol), BINAP (33.7 mg, 0.054 mmol), and Pd(OAc)_2_ (12.2 mg, 0.054 mmol) 25°C. The mixture was degassed and purged with N_2_ three times, and then heated in a microwave reactor at 120°C for 2 h. The mixture was filtered and concentrated to give a residue that was purified by prep-HPLC purification (column: Phenomenex luna C18 150*25 mm*10 μm; mobile phase: [water (HCO_2_H)-MeCN]; B%: 30–60%, 10 min) to give product **2k** (19.3 mg, 11.6% yield) as a white solid. ^1^H NMR (400 MHz, DMSO-*d*_6_) δ 9.68 (s, 1H), 9.06 (s, 1H), 8.35 (s, 1H), 8.24-8.15 (m, 2H), 7.99-7.88 (m, 1H), 7.30 (d, *J* = 8.8 Hz, 1H), 6.06 (s, 1H), 3.89-3.75 (m, 1H), 3.71-3.62 (m, 3H), 3.57-3.48 (m, 1H), 2.60-2.52 (m, 1H), 2.53 (s, 3H), 2.14 (s, 3H), 2.00-1.88 (m, 1H), 1.86-1.74 (m, 1H), 0.86-0.78 (m, 2H), 0.76-0.68 (m, 2H). ^13^C NMR (101 MHz, DMSO-d_6_) δ 171.96, 155.87, 149.91, 149.11, 145.60, 139.08, 138.85, 129.13, 119.45, 117.37, 114.13, 78.81, 77.86, 39.99, 39.00, 23.50, 6.58. LCMS: R_t_ = 0.393 min in 0.8 min chromatography, purity 100.0%, MS ESI calcd. for C_23_H_27_N_8_O_2_ [M+H]^+^ 447.22, found 447.1. HPLC: R_t_ = 1.850 min in 4 min chromatography, purity 97.4%.

*(R)-N-(5-((3-cyano-7-(cyclopropylamino)pyrazolo[1,5-a]pyrimidin-5-yl)amino)-2-(methyl(tetrahydrofuran-3-yl)amino)phenyl)acetamide (**2l**).* Intermediate **y** was synthesized from (*R*)-N-methyltetrahydrofuran-3-amine hydrochloride and *N*-(2-fluoro-5-nitrophenyl)acetamide by Procedure A. To a solution of 5-chloro-7-(cyclopropylamino)pyrazolo[1,5-a]pyrimidine-3-carbonitrile^6^ (98 mg, 0.421 mmol) and intermediate **y** (105 mg, 0.421 mmol) in dioxane (5 mL) was added Cs_2_CO_3_ (411 mg, 1.26 mmol), BINAP (39.3 mg, 0.063 mmol), and Pd(OAc)_2_ (14.2 mg, 0.063 mmol) 25°C. The mixture was degassed and purged with N_2_ three times, and then heated in a microwave reactor at 120°C for 2 h. The reaction mixture was filtered and concentrated under reduced pressure to give a residue that was purified by prep-TLC (SiO_2_, CH_2_Cl_2_/MeOH = 20/1), and the crude product was purified by prep-HPLC (column: Phenomenex Luna C18 150*25 mm*10 μm; mobile phase: [water (HCO_2_H)-MeCN]; B%: 29–59%, 8 min). Compound **2l** (18.2 mg, 9.71% yield) was obtained as a white solid. ^1^H NMR (400 MHz, DMSO-*d*_6_) δ 9.67 (s, 1H), 8.98 (s, 1H), 8.34 (s, 1H), 8.17 (d, *J* = 12.8 Hz, 2H), 7.92 (br d, *J* = 5.2 Hz, 1H), 7.29 (d, *J* = 8.8 Hz, 1H), 6.05 (s, 1H), 3.88 -3.79 (m, 1H), 3.71-3.62 (m, 3H), 3.56-3.47 (m, 1H), 2.62-2.53 (m, 1H), 2.52 (s, 3H), 2.14 (s, 3H), 1.99-1.86 (m, 1H), 1.84-1.76 (m, 1H), 0.83-0.77 (m, 2H), 0.73-0.68 (m, 2H). ^13^C NMR (101 MHz, DMSO-*d*_6_) δ 172.48, 156.90, 150.87, 148.20, 145.05, 138.50, 137.18, 134.98, 123.04, 114.87, 76.44, 70.96, 67.04, 63.59, 41.65, 30.73, 23.31, 6.53. LCMS R_t_ = 0.395 min in 0.8 min chromatography, purity 44.5%, MS ESI calcd. for 446.50 [M+H]^+^ 447.50, found 447.1. HPLC: Rt = 1.011 min in 4 min chromatography, purity 96.7%.

*(R)-N-(5-((3-cyano-7-(cyclopropylamino)pyrazolo[1,5-a]pyrimidin-5-yl)amino)-2-((tetrahydro-2H-pyran-3-yl)amino)phenyl)acetamide (**2m**).* Intermediate **y** was synthesized from (*R*)-tetrahydro-2H-pyran-3-amine hydrochloride and *N*-(2-fluoro-5-nitrophenyl)acetamide by Procedure A. To a solution of 5-chloro-7-(cyclopropylamino)pyrazolo[1,5-a]pyrimidine-3-carbonitrile^6^ (220 mg, 0.943 mmol) and intermediate **y** (235 mg, 0.943 mmol) in dioxane (6 mL) was added Cs_2_CO_3_ (915 mg, 2,83 mmol), BINAP (87.6 mg, 0.13 mmol), and Pd(OAc)_2_ (30.2 mg, 0.13 mmol) 25°C. The mixture was degassed and purged with N_2_ three times, and then heated in a microwave reactor at 120°C for 2 h. The mixture was filtered and concentrated to give a residue that was purified by prep-HPLC purification (column: Phenomenex luna C18 150*25 mm*10 μm; mobile phase: [water (HCO_2_H)-MeCN]; B%: 26–56%, 10 min, and column: Phenomenex C18 150*25 mm*10 μm; mobile phase: [water (NH_4_HCO_3_)-MeCN]; B%: 22%-52%, 14 min) to give compound **2m** (10 mg, 22.40 μmol, 2.38% yield) as an off-white solid. ^1^H NMR (400 MHz, DMSO-*d*_6_) δ 9.33 (s, 1H), 9.19 (s, 1H), 8.29 (s, 1H), 8.09 (s, 1H), 7.45 (s, 1H), 7.39 (s, 1H), 6.73 (d, *J* = 8.8 Hz, 1H), 5.90 (s, 1H), 4.54 (d, *J* = 8.0 Hz, 1H), 3.89-3.82 (m, 1H), 3.75-3.67 (m, 1H), 3.42-3.38 (m, 2H), 3.27-3.11 (m, 1H), 2.59-2.52 (m, 1H), 2.06 (s, 3H), 2.01-1.93 (m, 1H), 1.75-1.67 (m, 1H), 1.63-1.45 (m, 2H), 0.82-0.74 (m, 2H), 0.72-0.65 (m, 2H). HPLC: R_t_ = 1.670 min in 4 min chromatography, purity 97.6%. LCMS R_t_ = 0.365 min in 0.8 min chromatography, purity 99.1%, MS ESI calcd. for 446.22 [M+H]^+^ 447.22, found 447.2.

*N-(5-((3-cyano-7-(cyclopropylamino)pyrazolo[1,5-a]pyrimidin-5-yl)amino)-2-(methyl(oxetan-3-ylmethyl)amino)phenyl)acetamide (**2n**).* Intermediate **y** was synthesized from *N*-methyl-1-(oxetan-3-yl)methanamine hydrochloride and *N*-(2-fluoro-5-nitrophenyl)acetamide by Procedure A. To a solution of 5-chloro-7-(cyclopropylamino)pyrazolo[1,5-a]pyrimidine-3-carbonitrile^6^ (293 mg, 1.26 mmol) and intermediate **y** (313 mg, 1.26 mmol) in dioxane (10 mL) was added Cs_2_CO_3_ (1.64 g, 5.02 mmol), BINAP (117 mg, 0.188 mmol), and Pd(OAc)_2_ (42.2 mg, 0.188 mmol) at 25°C. The mixture was degassed, purged with N_2,_ and then heated in a microwave reactor at 120 °C for 2 hours. The reaction mixture was diluted with CH_2_Cl_2_/MeOH (10/1, 30 mL), then filtered, and the filtrate was concentrated to give a crude product that purified by flash silica gel chromatography (Eluent of 30–80% EtOAc/petroleum ether). The crude product was further purified by prep-HPLC (column: Phenomenex luna C18 150*25 mm*10 μm; mobile phase: [water (HCO_2_H)-MeCN]; B%: 22–52%, 10 min) to give Compound **2n** (150.6 mg, 32.9%) as a light-yellow solid. ^1^H NMR (400 MHz, DMSO-*d*_6_) δ 9.56 (s, 1H), 8.93 (s, 1H), 8.33 (s, 1H), 8.17 (s, 1H), 8.10 (s, 1H), 7.88 (d, *J* = 7.2 Hz, 1H), 7.24 (d, *J* = 8.8 Hz, 1H), 6.04 (s, 1H), 4.53 (t, *J* = 7.2 Hz, 2H), 4.22 (t, *J* = 6.0 Hz, 2H), 3.15-3.09 (m, 2H), 3.08-3.01 (m, 1H), 2.56 (s, 3H), 2.55-2.51 (m, 1H), 2.12 (s, 3H), 0.84-0.76 (m, 2H), 0.75-0.66 (m, 2H). ^13^C NMR (101 MHz, DMSO-*d*_6_) δ 168.25, 156.93, 150.89, 148.20, 145.01, 136.87, 136.41, 134.17, 122.04, 114.96, 114.85, 111.72, 76.34, 74.77, 59.67, 42.84, 32.92, 24.24, 23.30, 6.52. HPLC R_t_ = 2.093 min in 4 min chromatography, purity 96.6%. LCMS R_t_ = 1.136 min in 3 min chromatography, purity 100%, MS ESI calcd. for 446.22 [M+H]^+^ 447.22, found 447.2.

## Ancillary Information

*Supporting Information:* SMARTCyp prediction of P450 metabolism of SGC-CK2-1 (**1a**), inhibition of **2h** metabolism by 1-ABT, dose ranging study of EA tolerability in mice by i.p. dosing, and NMR spectra (PDF).

## Author Contributions

T.M.W, A.D.A., and N.J.M conceived of the study. X.Y., H.W.O., J.W.B, W.T., E.C., and A.D.A designed experiments and compounds. R.J.D. and J.L.S. performed biological studies. T.M.W wrote the manuscript. All authors read and approved the manuscript.

## Notes

The authors declare no competing financial interest.

## Acknowledgement

Constructs for NanoBRET measurements of CSNK2A1 and CSNK2A2, were provided by Promega Corporation (Madison, WI). We thank K. Saikatendu Singh (Takeda, San Diego, CA) for facilitating collaborative interactions and constructive criticism throughout the project. WuXi AppTec (Shanghai, China) provided chemical synthesis support. Analiza, Inc (Cleveland, OH) performed kinetic solubility and mouse liver microsome studies. Pharmaron (Beijing, China) provided in vitro and in vivo pharmacokinetic assay support.

## Abbreviations Used

CSNK2A: Casein Kinase 2a
PZP: 3-cyano-7-cyclopropylamino-pyrazolo[1,5-a]pyrimidine
GST: glutathione S-transferase
MHV: mouse hepatitis virus
MLM: mouse liver microsome
i.v.: intravenous
i.p.: intraperitoneal
C_max_: maximum concentration
AUC: area under the curve
CL_int_: intrinsic clearance
EA: ethacrynic acid
1-ABT: 1-aminobenzotriazole.

