## Supporting Information for "Optimization of 3-Cyano-7-cyclopropylamino-pyrazolo[1,5-a]pyrimidines Toward the Development of an In Vivo Chemical Probe for CSNK2A"

**Table of Contents**

S2 **Figure S1.** SMARTCyp analysis of SGC-CK2-1 (**1a**)

S3 **Figure S2.** MS fragmentation analysis and structure assignment of **1i** metabolites M1–12.

S9 **Figure S3.** 1-ABT inhibits the metabolism of **2h** in mouse hepatocytes

S10 **Table S1.** Dose ranging study of EA tolerability in mice

S11 NMR and HPLC Spectra for **1h–i**, **2b–g, 2i–n**

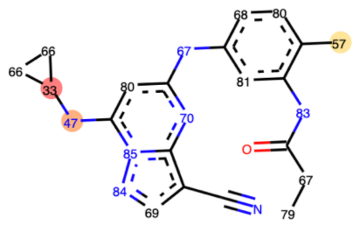


| 3A4 Ranking | Atom | 3A4 Score | Energy | 2DSASA | Span2end | Relative Span | Similarity |
|---|---|---|---|---|---|---|---|
| 1 | C.21 | 33.0 | 41.1 | 17.1 | 1 | 0.9 | 0.3 |
| 2 | N.20 | 46.8 | 54.1 | 13.5 | 2 | 0.8 | 0.3 |
| 3 | C.28 | 57.3 | 66.4 | 58.7 | 2 | 0.8 | 0.7 |
| 4 | C.22 | 65.9 | 75.9 | 50.1 | 0 | 1.0 | 0.3 |
| 5 | N.9 | 67.2 | 72.0 | 12.3 | 6 | 0.5 | 0.7 |
| 6 | C.3 | 67.3 | 75.9 | 30.6 | 1 | 0.9 | 0.7 |
| 7 | C.25 | 68.1 | 74.1 | 26.4 | 5 | 0.6 | 0.7 |
| 8 | C.16 | 69.4 | 78.1 | 33.4 | 1 | 0.9 | 0.3 |
| 9 | N.11 | 69.6 | 75.6 | 11.0 | 4 | 0.7 | 0.3 |
| 10 | C.4 | 79.0 | 89.6 | 64.4 | 0 | 1.0 | 0.7 |
| 11 | C.26 | 79.7 | 86.3 | 27.5 | 4 | 0.7 | 0.7 |
| 12 | C.24 | 80.0 | 86.3 | 18.6 | 4 | 0.7 | 0.3 |
| 13 | C.7 | 80.6 | 86.3 | 19.9 | 5 | 0.6 | 0.3 |
| 14 | N.5 | 83.0 | 89.6 | 10.8 | 3 | 0.8 | 0.3 |
| 15 | N.17 | 83.8 | 92.1 | 22.1 | 1 | 0.9 | 0.3 |
| 16 | N.18 | 85.2 | 92.1 | 3.3 | 2 | 0.8 | 0.3 |

**Figure S1.** SMARTCyp analysis of SGC-CK2-1 (**1a**). Output from the SMARTCyp server at https://smartcyp.sund.ku.dk/mol_to_som. The predicted hotspots of cytochrome CYP3A4 metabolism are ranked for each heavy atom in the molecule. The top ranked atoms are located on the cycloproylamine of the PZP and the 4'-methyl group of the aniline.

**M1**

RT: 5.68 min;

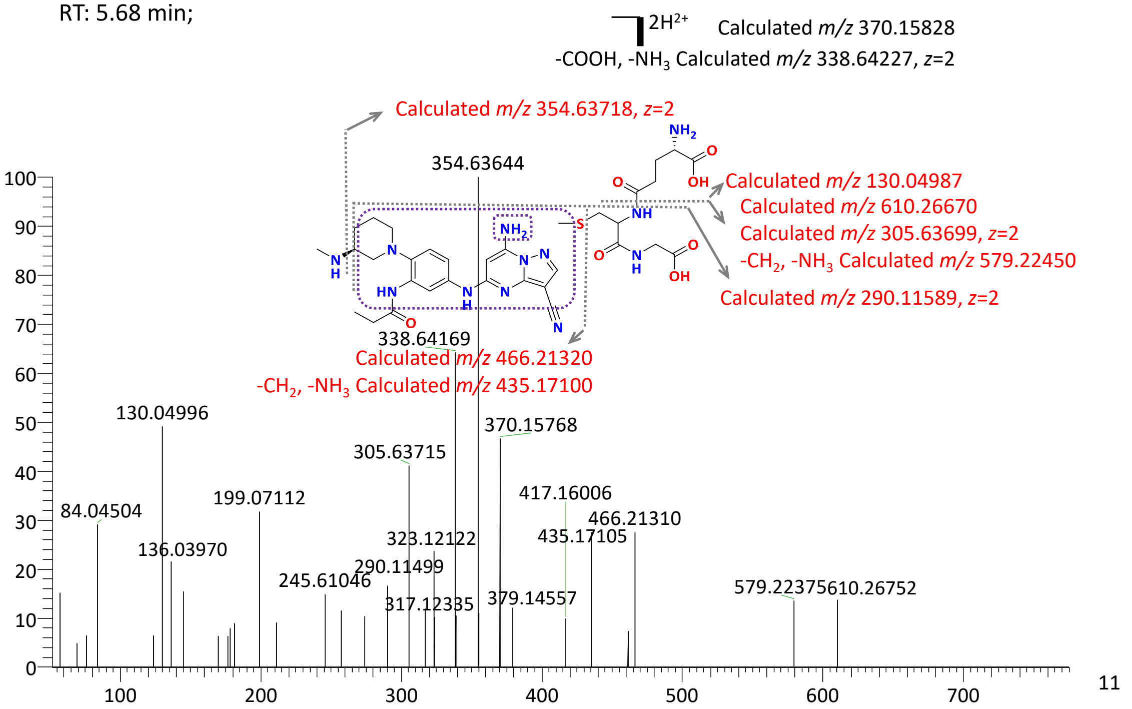


**M2**

RT: 6.03 min;

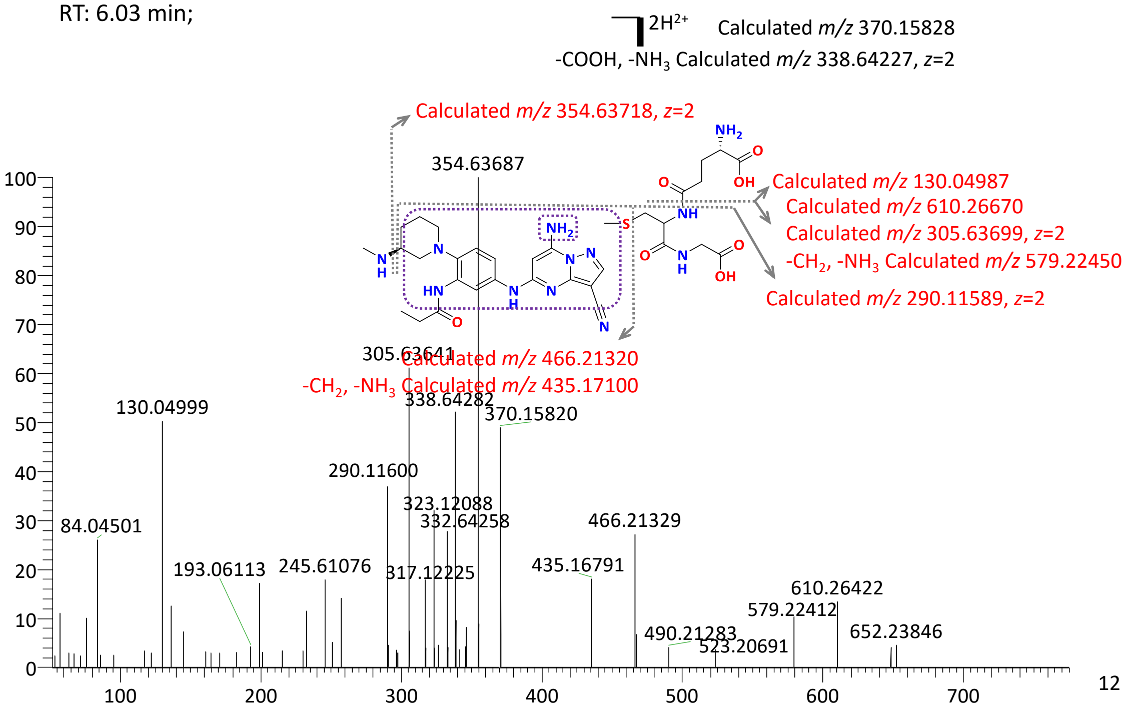

**M3**

RT: 6.89 min;

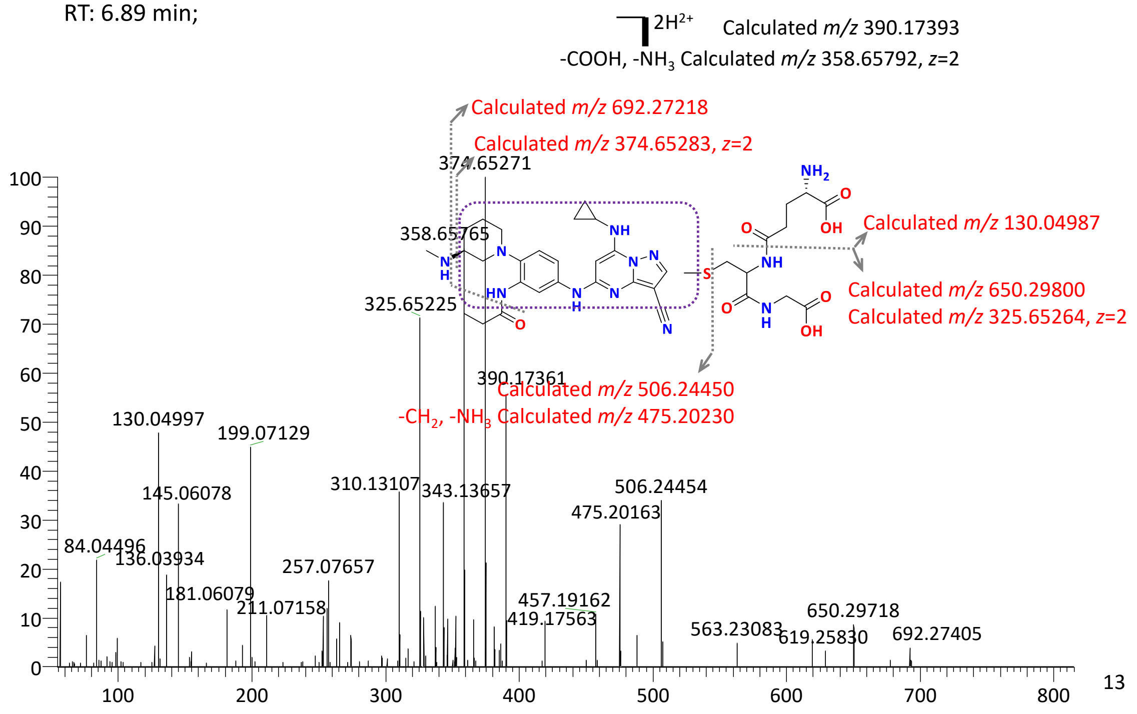


**M4**

RT: 7.10 min;

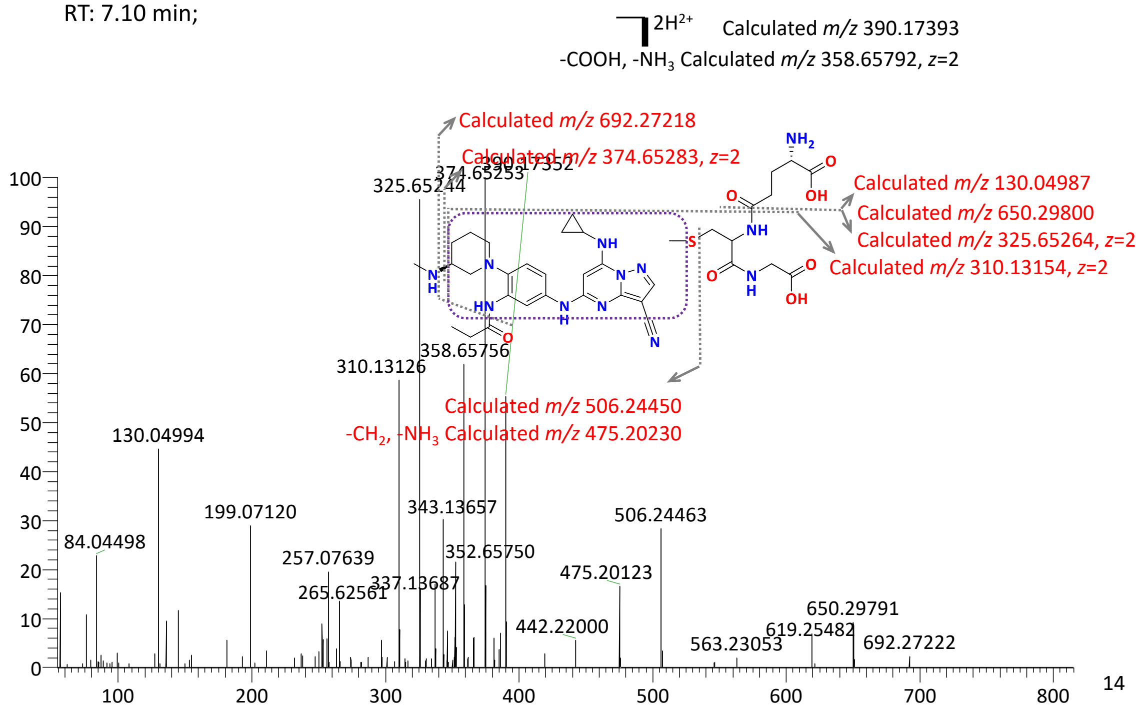

## M5

RT: 7.10 min;

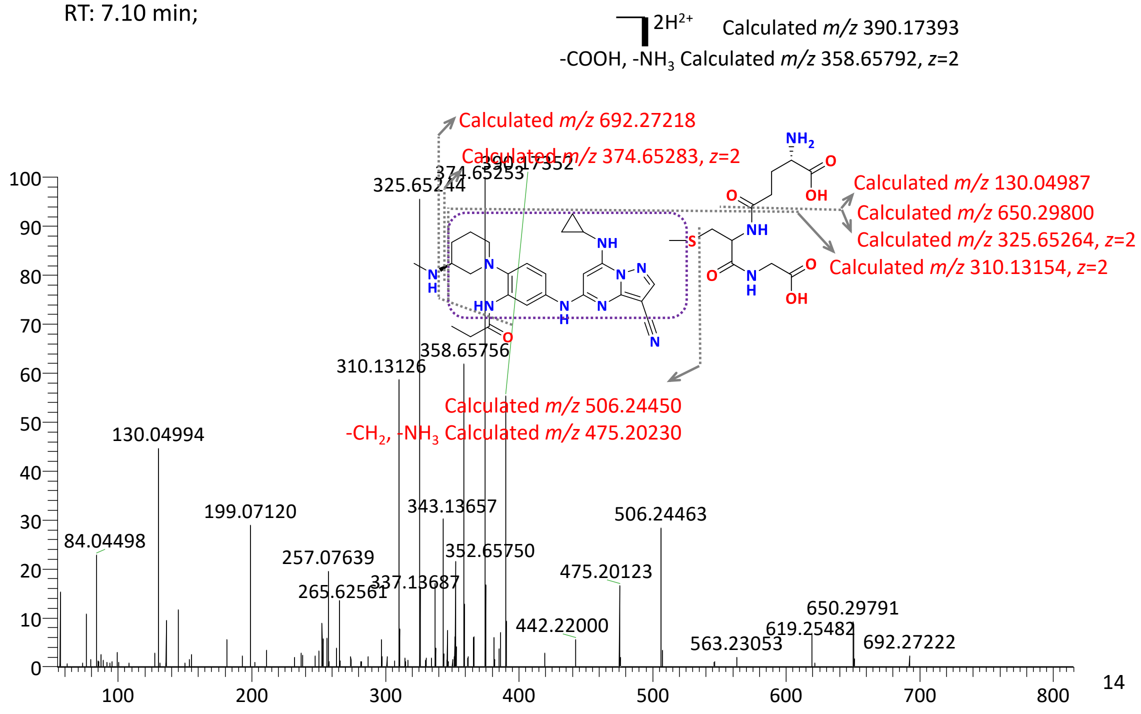


## M6

RT: 7.59 min;

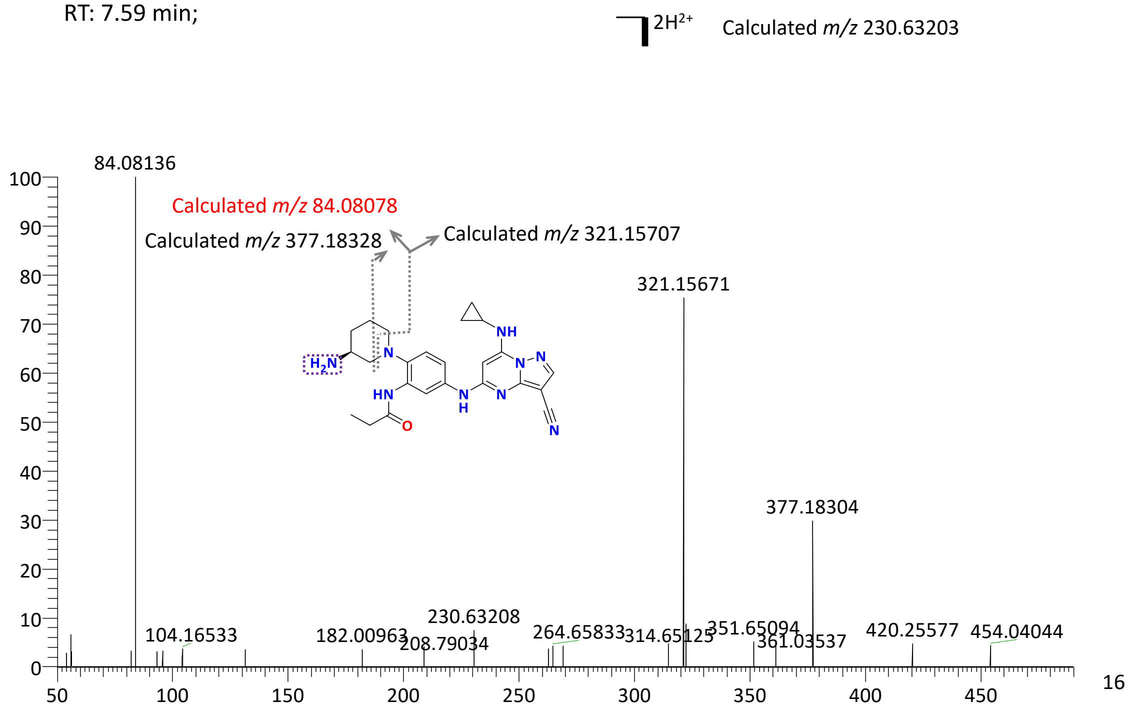

## M7

RT: 7.70 min;

$]H^+$ Calculated *m/z* 472.25678

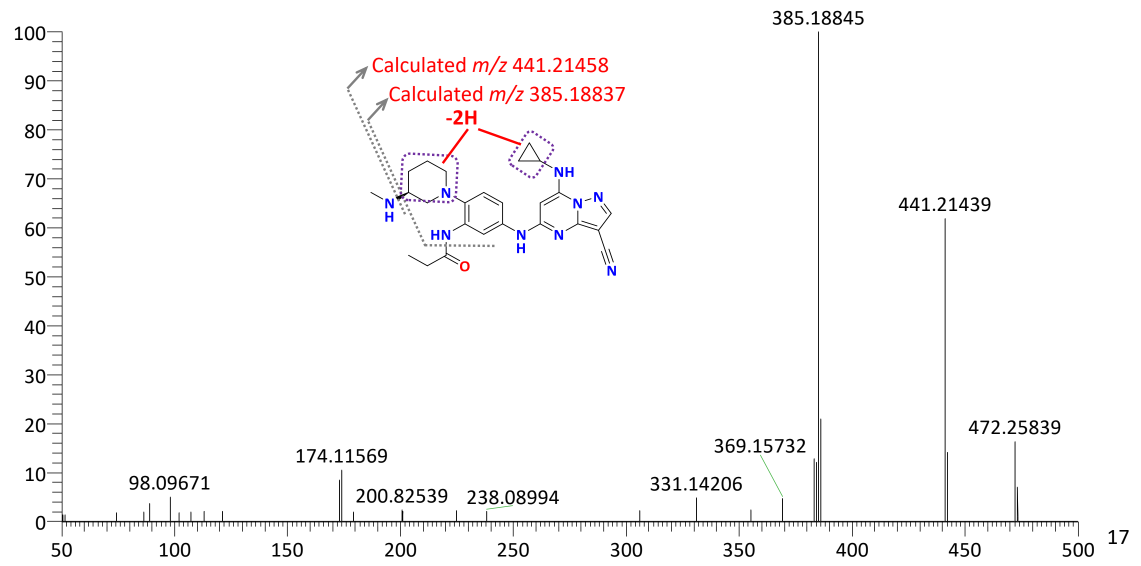


## M8

RT: 7.79 min;

$]H^+$ Calculated *m/z* 418.24622

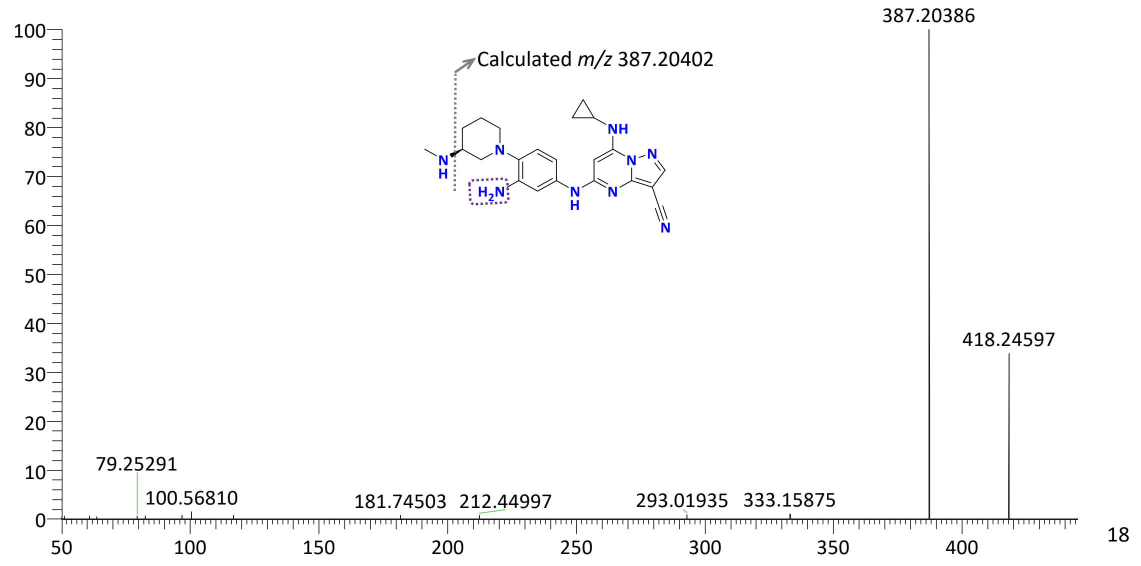

## M9

RT: 7.80 min;

H⁺ Calculated *m/z* 417.17820

$-H_2O$ Calculated *m/z* 399.16763

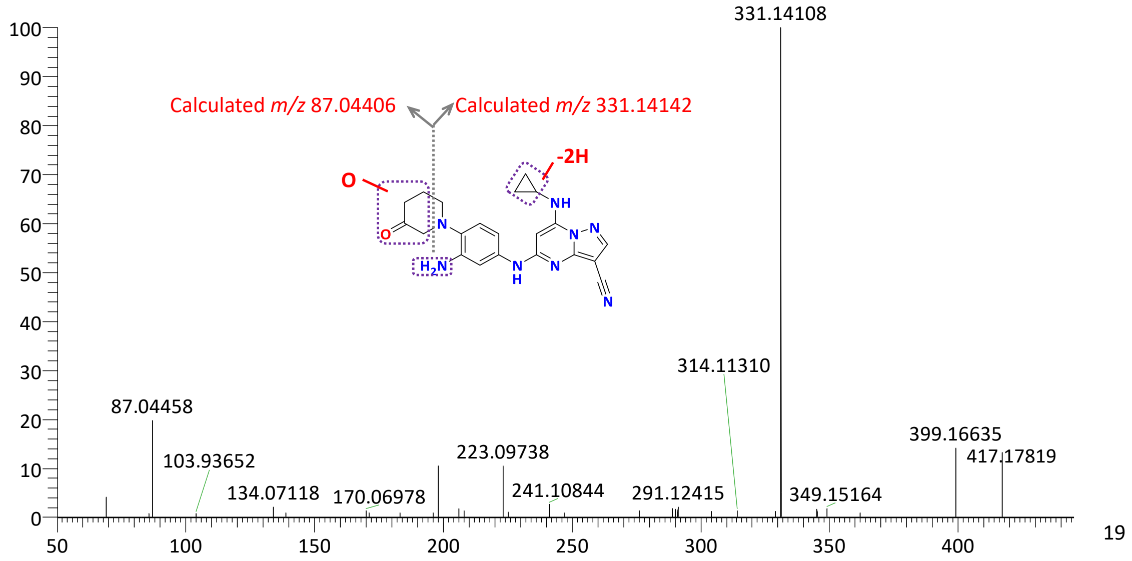


## M10

RT: 7.91 min;

$2H^{2+}$ Calculated *m/z* 342.12137

$-NH_3$ Calculated *m/z* 333.60809, *z*=2

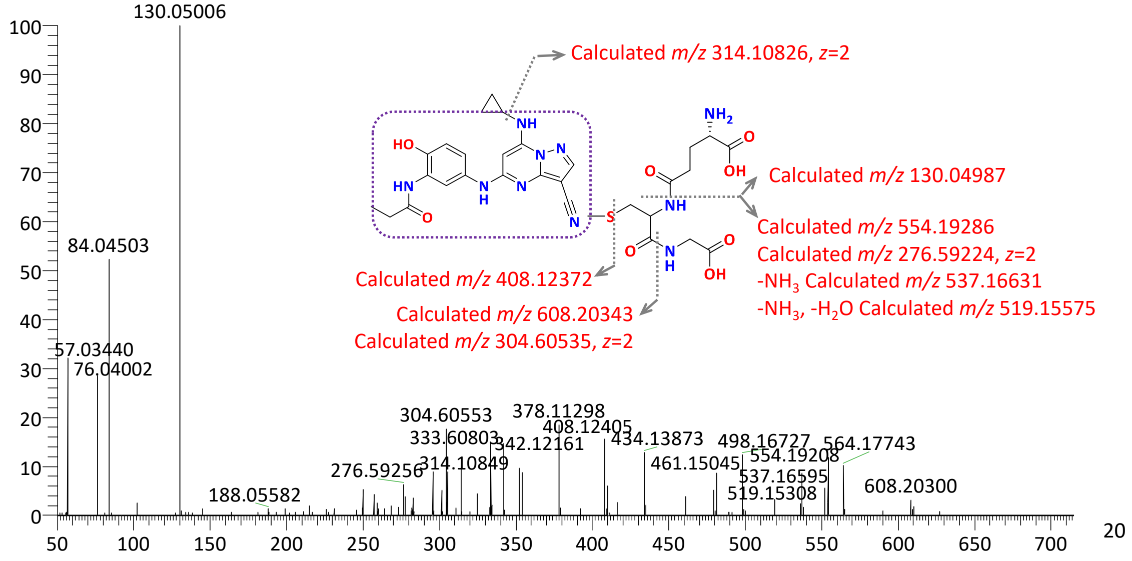

## M11

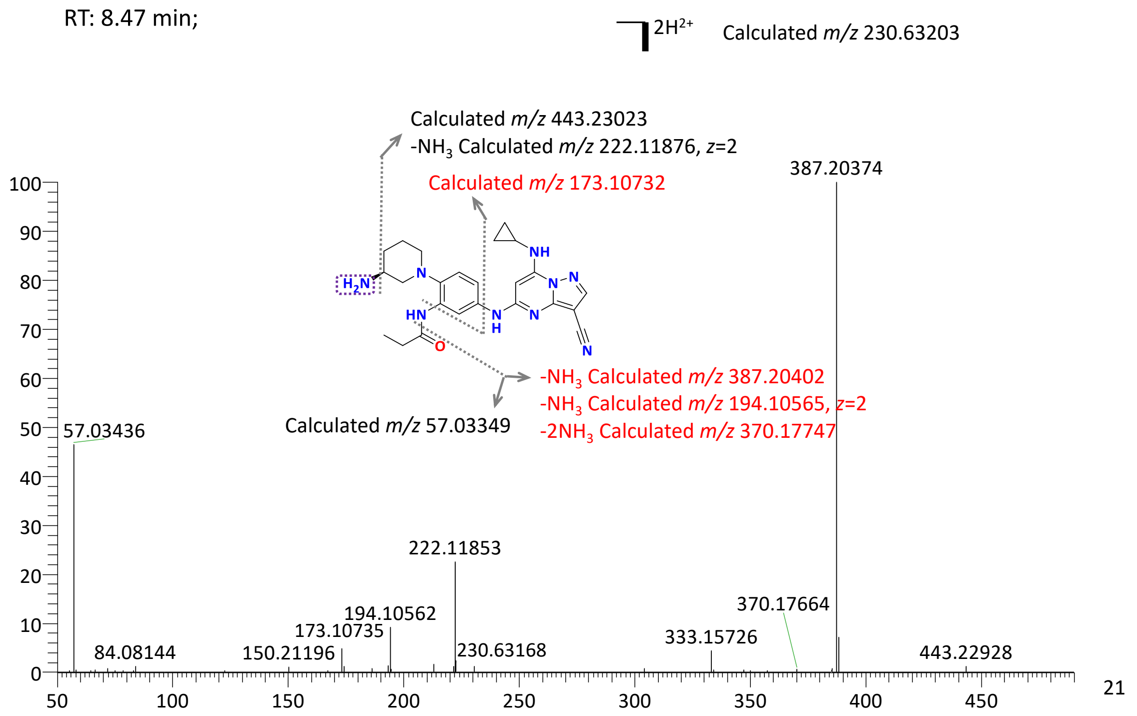


## M12

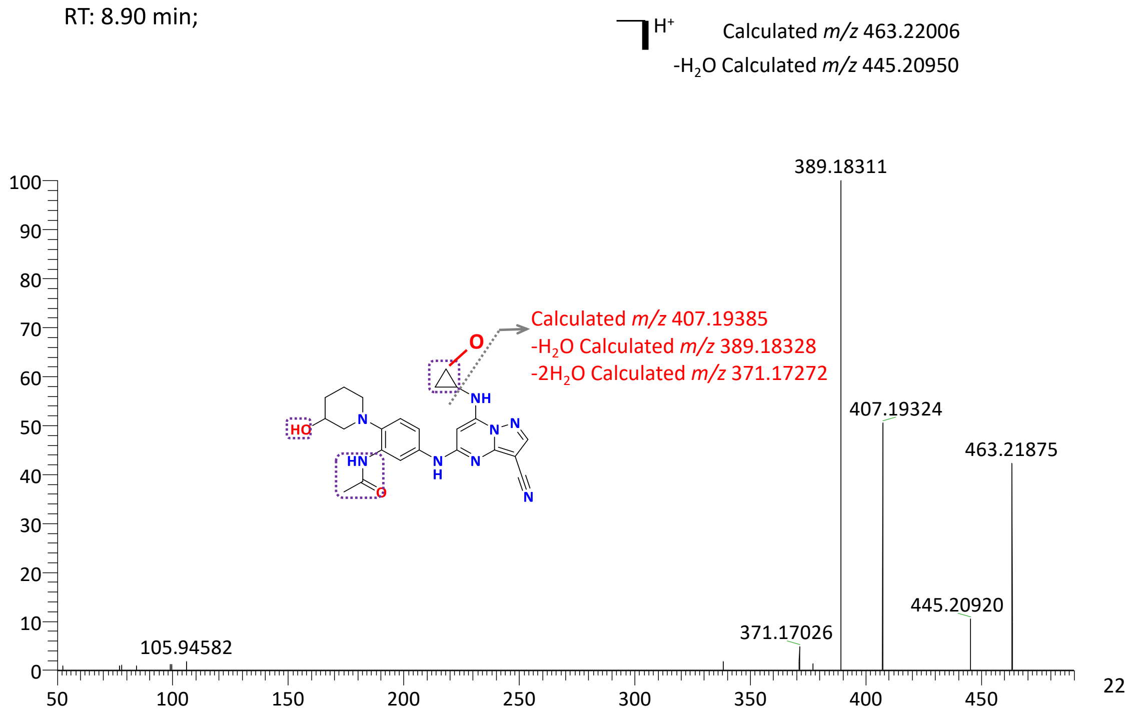


**Figure S2.** MS fragmentation analysis and structure assignment of **1i** metabolites **M1–M12**.

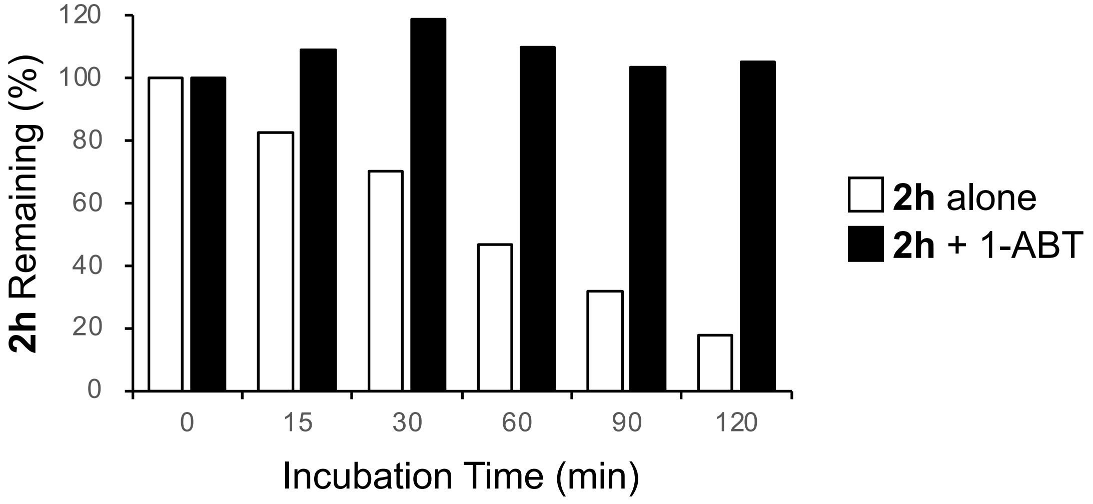


**Figure S3.** 1-ABT inhibits the metabolism of **2h** in mouse hepatocytes. **2h** (1 μM) was incubated with mouse hepatocytes in the presence or absence of 1-ABT (1 mM). The percentage of **2h** remaining was measured by LC/MS/MS at 0.25, 0.5, 1.0, 1.5, and 2.0h.

**Table S1. Dose ranging study of EA tolerability in mice.**

| Dose (mg/kg) | Route | Volume (mL/kg)[a] | Formul-ation[b] | Adverse effects after 1st dose[c] | Adverse effects after 2nd dose[c] |
|---|---|---|---|---|---|
| 300 | i.p. | 10 | A | Tachycardia, hypothermia, death @ 15 min | – |
| 100 | i.p. | 10 | A | Decreased activity, hypothermia, death @ 40 min | – |
| 30 | i.p. | 10 | A | Decreased activity | Decreased activity, death @ 60 min |
| 10 | i.p. | 10 | A | Decreased activity | Decreased activity, hypothermia, death @ 70 min |
| 3 | i.p. | 10 | A | Decreased activity | Decreased activity, hypothermia |
| 10 | i.p. | 1 | B | None | Decreased activity, hypothermia[d] |
| 10 | p.o. | 1 | B | None | Decreased activity, hypothermia[d] |

Mice were dosed at 6h intervals with EA and observations made of clinical symptoms. A single mouse was used for each experiment. [a]Injection volume. [b]Formulation A: NMP, PEG400, Water (v/v/v, 10:60:30), Formulation B: DMSO, PEG300, Tween80, 20% HP-β-CD in water (v/v/v/v, 10:40:5:45). [c]Clinical observations with time of death after dosing noted. [d]Symptoms resolved after 30 min. –, no second dose.

### NMR and HPLC spectra of 1h:

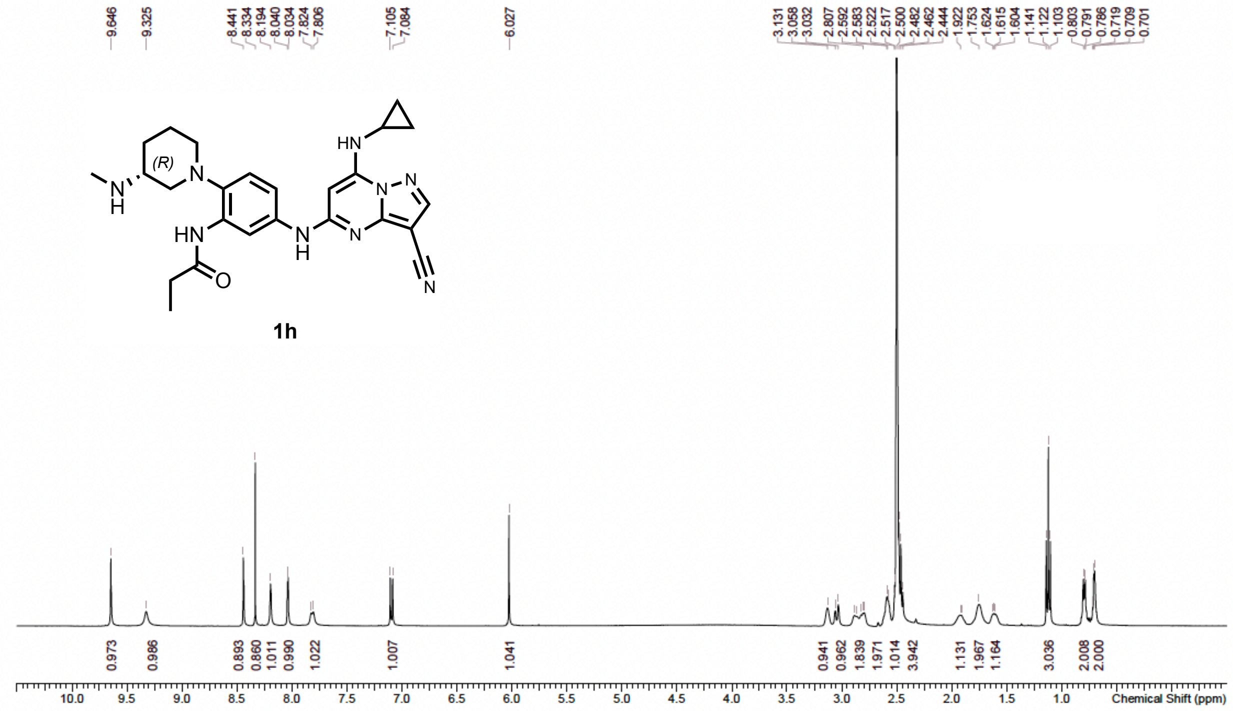


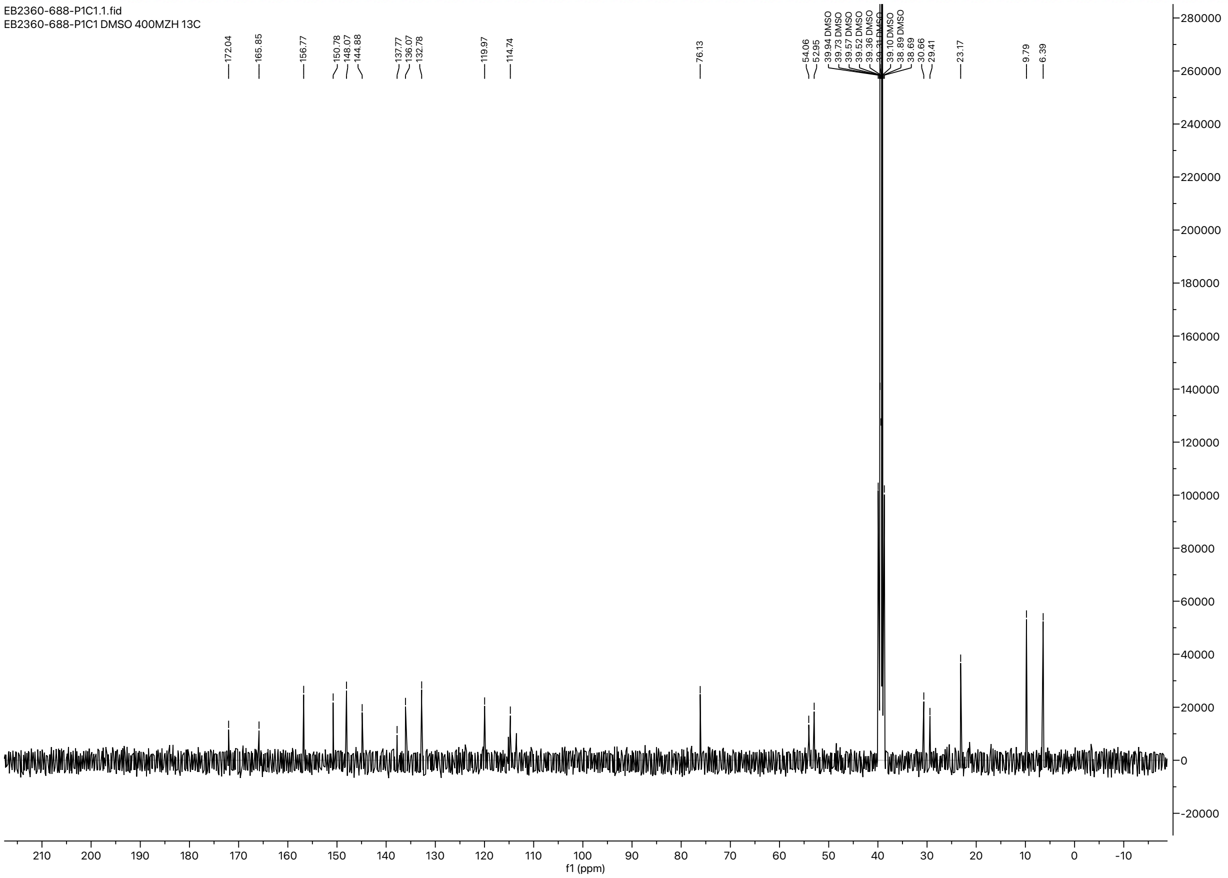

Chromatogram
WX-TKDU-F_009 D:\DATA\2021\2109\210915\EB2360-688-P1H1.lcd

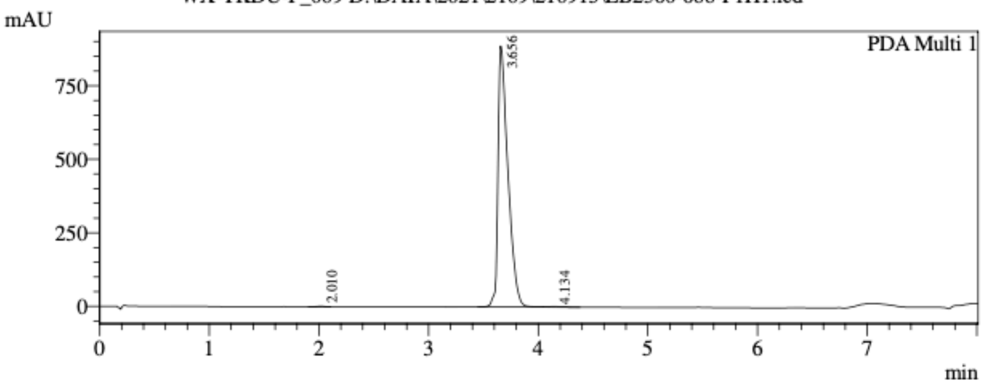


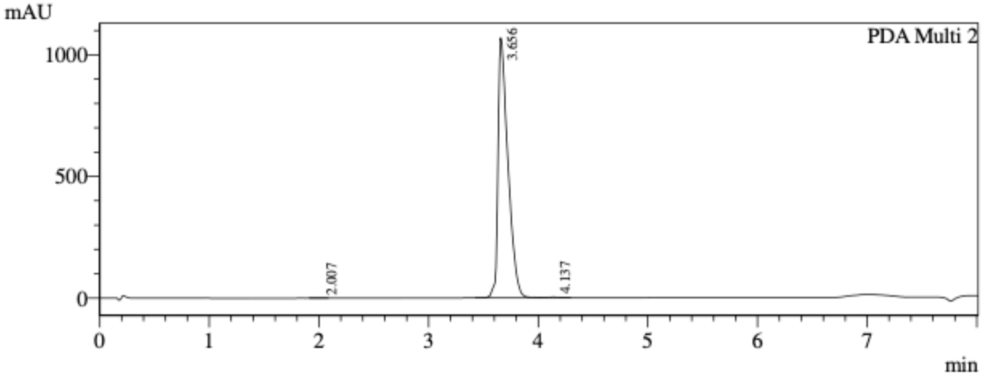


1 PDA Multi 1 / 220nm 4nm
2 PDA Multi 2 / 254nm 4nm

Integration Result

PeakTable

Ch1 220nm 4nm

| Peak# | Ret. Time | Height | Height % | Area | Area % |
|---|---|---|---|---|---|
| 1 | 2.010 | 1586 | 0.178 | 8125 | 0.145 |
| 2 | 3.656 | 886751 | 99.581 | 5578720 | 99.572 |
| 3 | 4.134 | 2145 | 0.241 | 15832 | 0.283 |
| Total | | 890482 | 100.000 | 5602677 | 100.000 |

PeakTable

Ch2 254nm 4nm

| Peak# | Ret. Time | Height | Height % | Area | Area % |
|---|---|---|---|---|---|
| 1 | 2.007 | 596 | 0.056 | 3088 | 0.046 |
| 2 | 3.656 | 1068344 | 99.725 | 6733801 | 99.686 |
| 3 | 4.137 | 2348 | 0.219 | 18112 | 0.268 |
| Total | | 1071288 | 100.000 | 6755001 | 100.000 |

PeakTable

Ch3

### NMR and HPLC spectra of 1i:

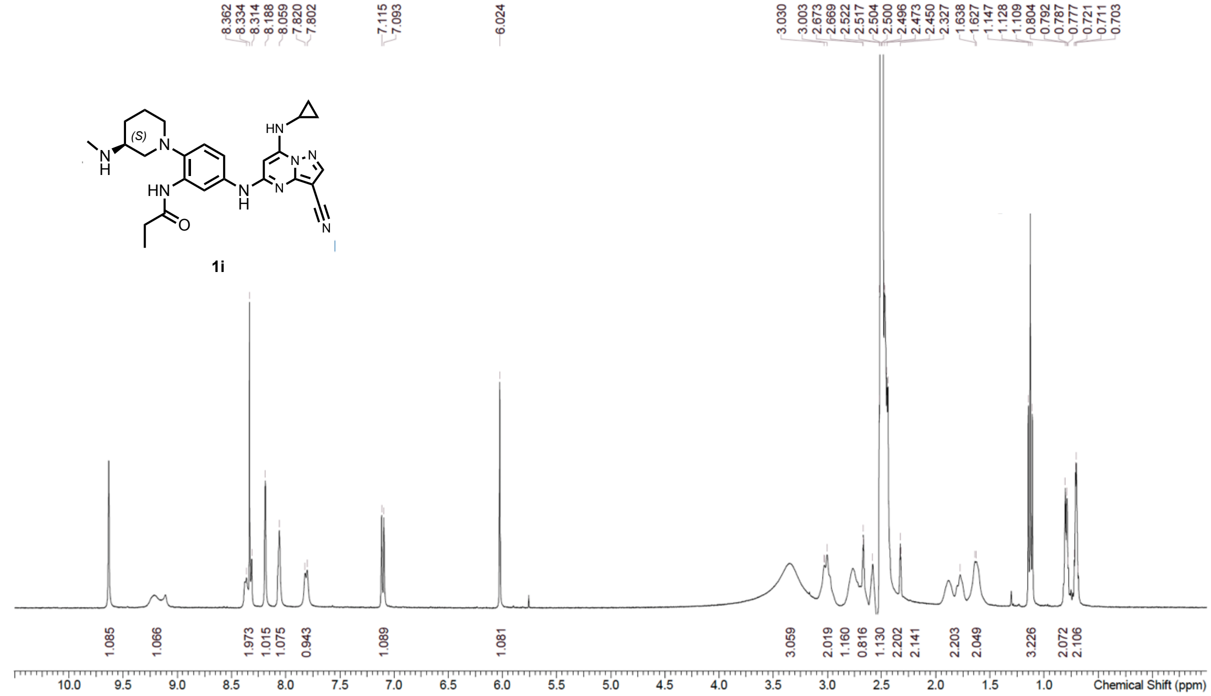


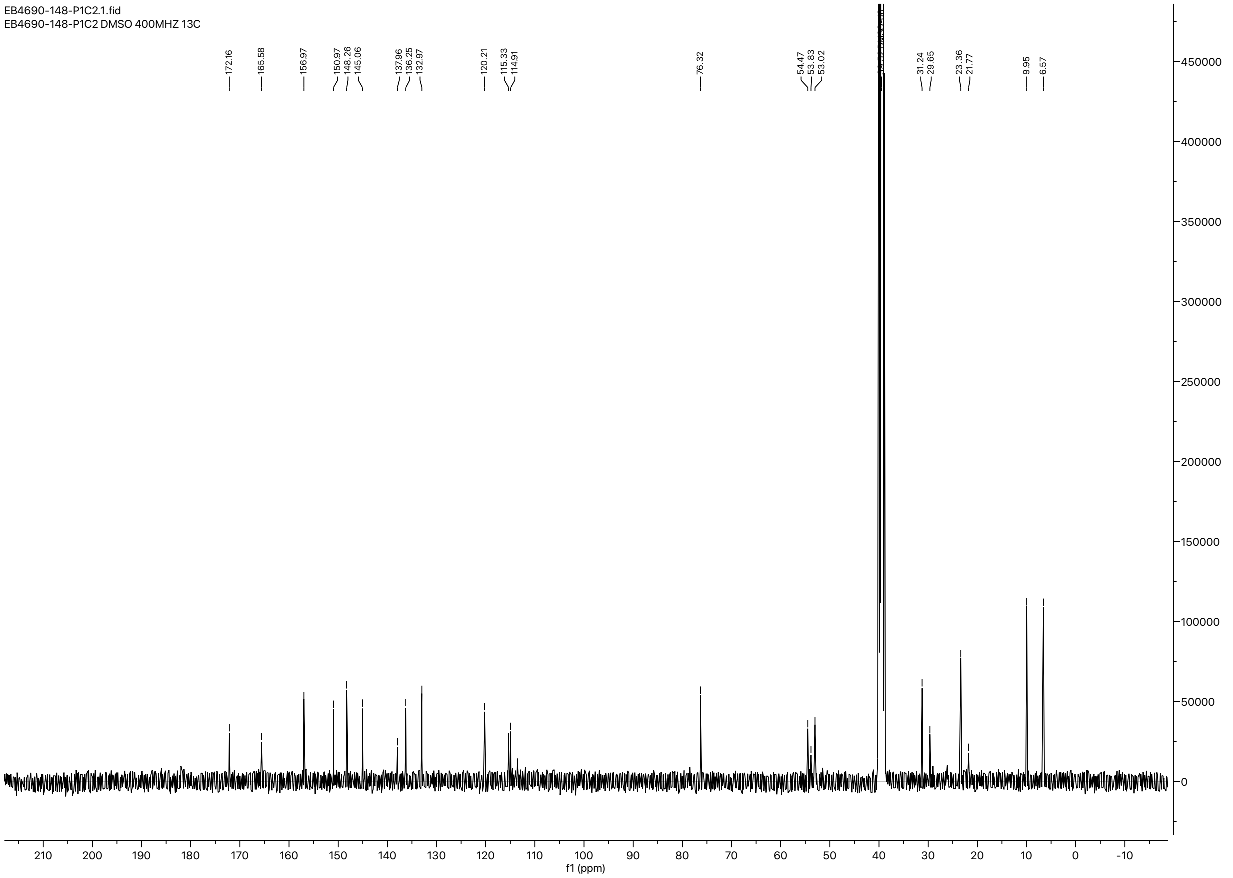

<Chromatogram>

Chromatogram
WX-TKDU-F_010 D:\DATA\2021\2109\210916\EB4690-148-P1H1.lcd

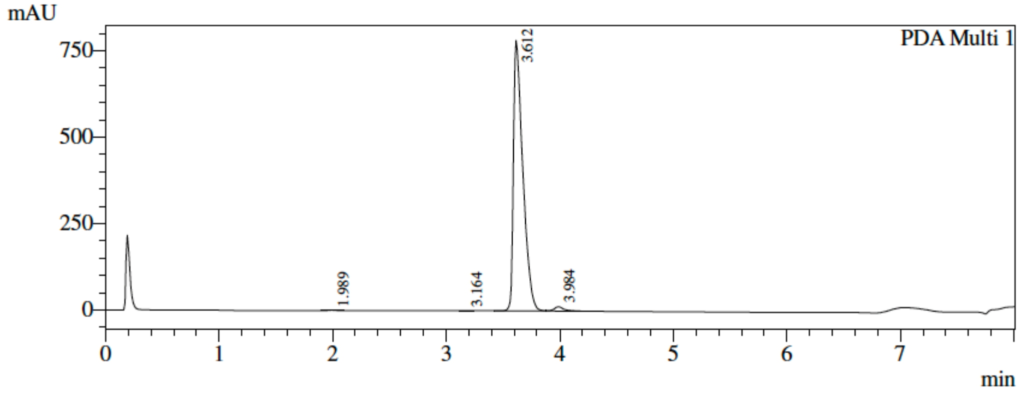


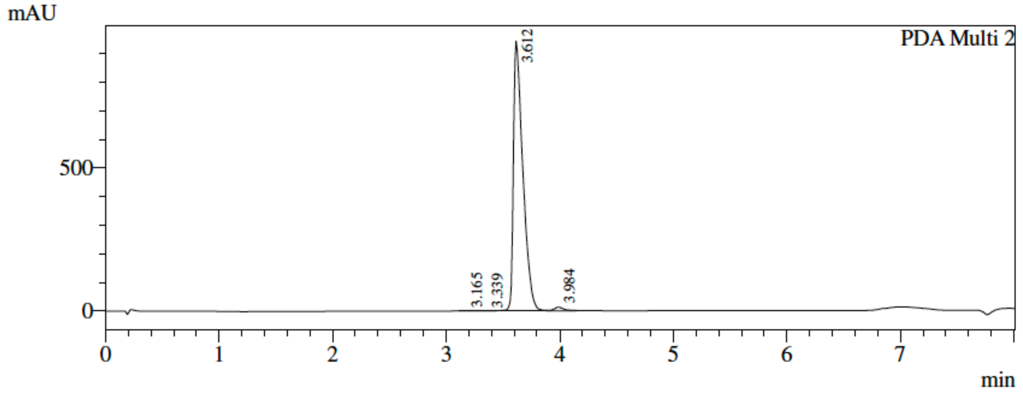


1 PDA Multi 1 / 220nm 4nm
2 PDA Multi 2 / 254nm 4nm

Integration Result

PeakTable

Ch1 220nm 4nm

| Peak# | Ret. Time | Height | Height % | Area | Area % |
|---|---|---|---|---|---|
| 1 | 1.989 | 1411 | 0.177 | 5667 | 0.123 |
| 2 | 3.164 | 842 | 0.106 | 2792 | 0.061 |
| 3 | 3.612 | 781856 | 98.124 | 4505087 | 98.071 |
| 4 | 3.984 | 12697 | 1.593 | 80171 | 1.745 |
| Total | | 796806 | 100.000 | 4593717 | 100.000 |

PeakTable

Ch2 254nm 4nm

| Peak# | Ret. Time | Height | Height % | Area | Area % |
|---|---|---|---|---|---|
| 1 | 3.165 | 615 | 0.064 | 2024 | 0.037 |
| 2 | 3.339 | 524 | 0.055 | 4383 | 0.079 |
| 3 | 3.612 | 942218 | 98.585 | 5445281 | 98.407 |
| 4 | 3.984 | 12384 | 1.296 | 81740 | 1.477 |
| Total | | 955740 | 100.000 | 5533428 | 100.000 |

PeakTable

Ch3

### NMR and HPLC spectra of 2a:

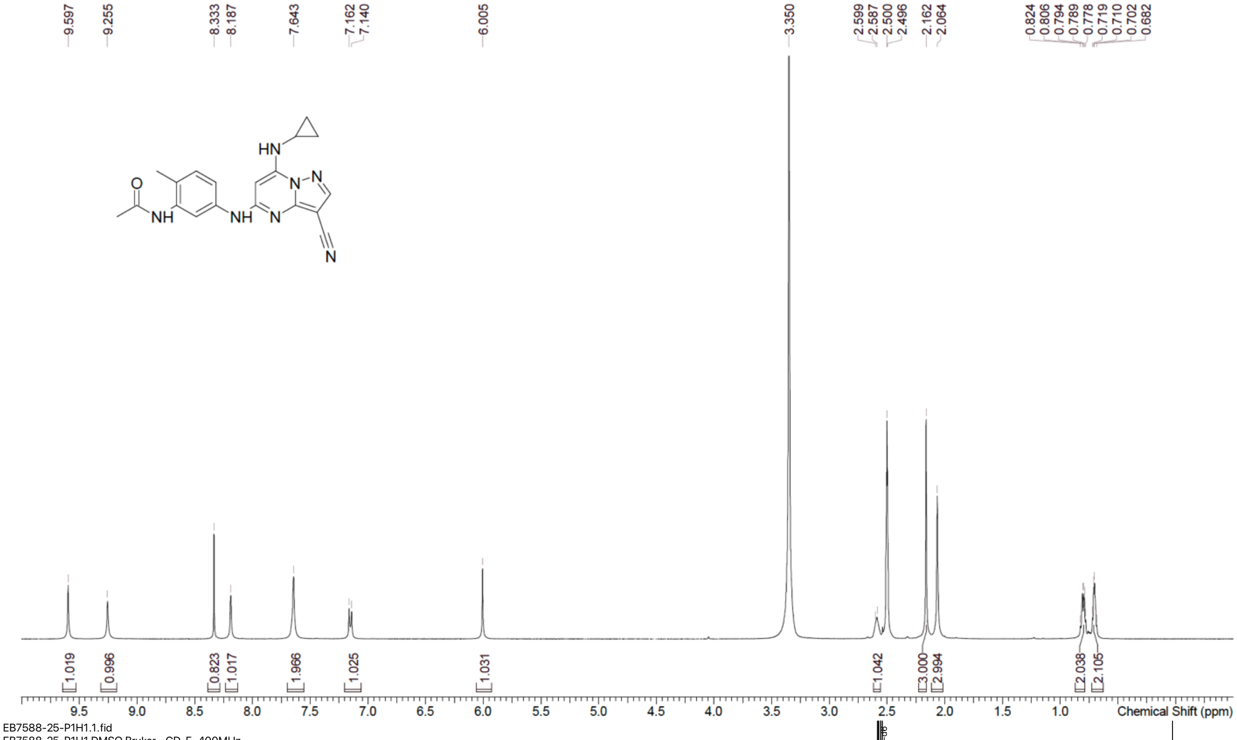


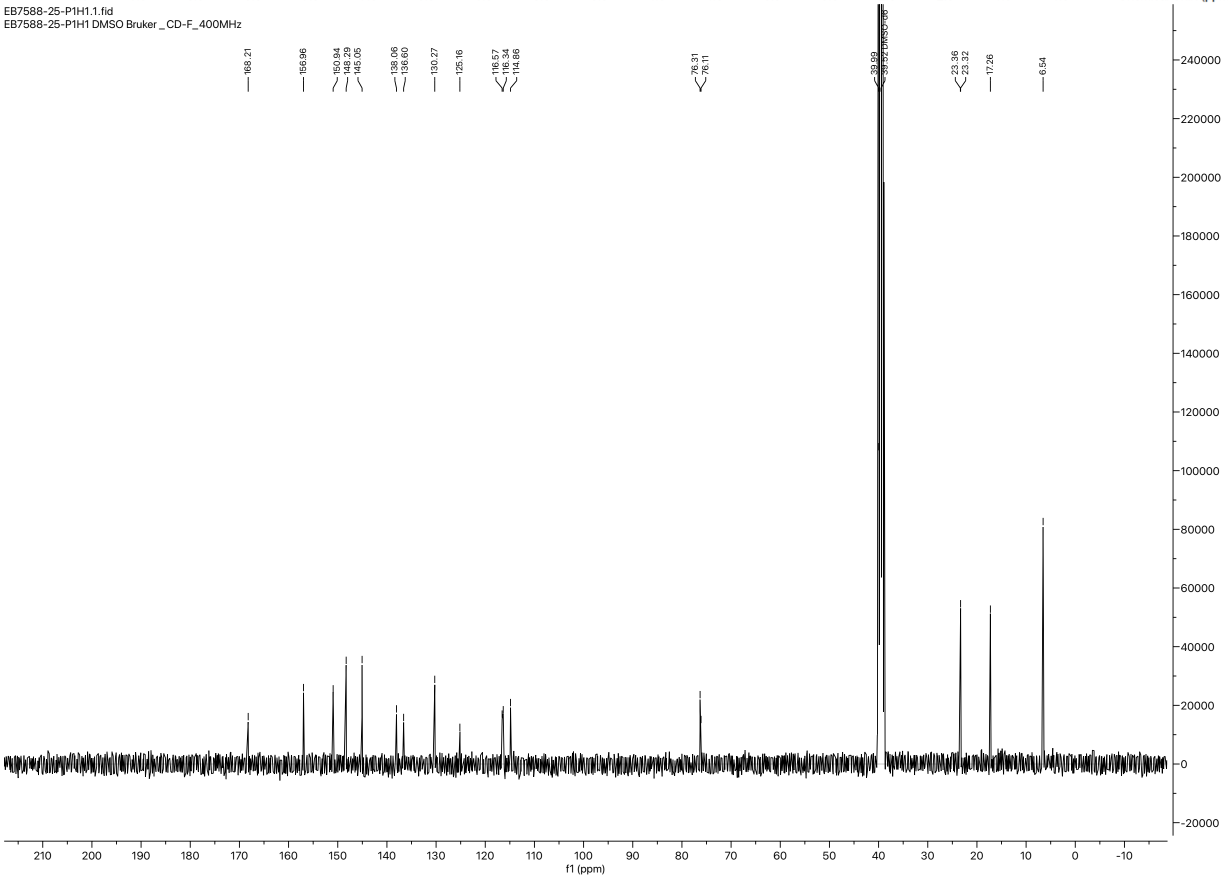

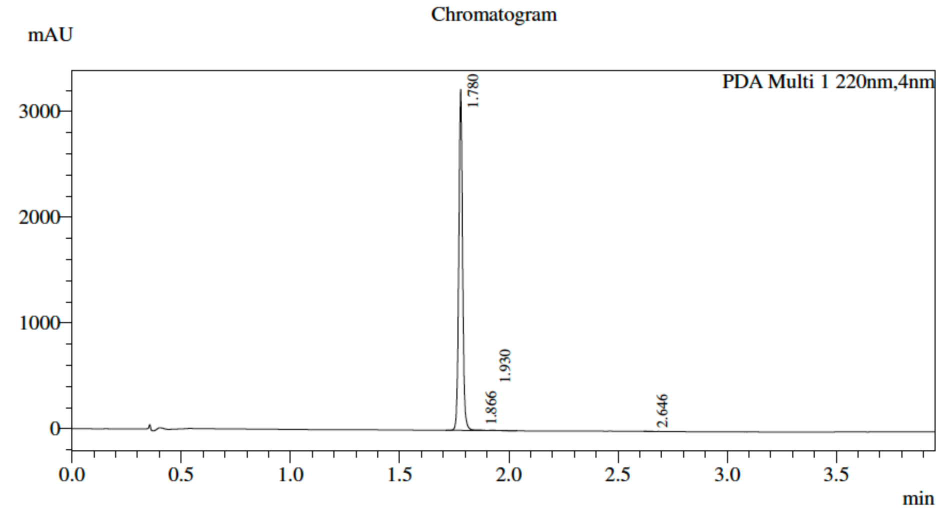


### Integration Result

PDA Ch1 220nm

| Peak# | Ret. Time | USP Width | Height | Height% | Area | Area% |
|---|---|---|---|---|---|---|
| 1 | 1.780 | 0.032 | 3226270 | 99.670 | 3879046 | 99.446 |
| 2 | 1.866 | 0.080 | 5152 | 0.159 | 9754 | 0.250 |
| 3 | 1.930 | 0.066 | 3910 | 0.121 | 9447 | 0.242 |
| 4 | 2.646 | 0.039 | 1610 | 0.050 | 2390 | 0.061 |
| Total | | | 3236942 | 100.000 | 3900637 | 100.000 |

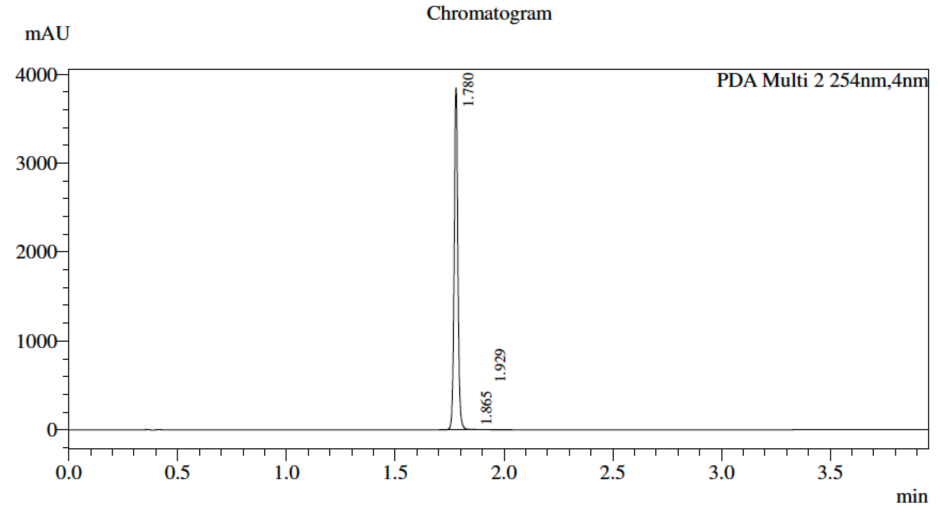


### Integration Result

PDA Ch2 254nm

| Peak# | Ret. Time | USP Width | Height | Height% | Area | Area% |
|---|---|---|---|---|---|---|
| 1 | 1.780 | 0.028 | 3846935 | 99.780 | 4569098 | 99.570 |
| 2 | 1.865 | 0.101 | 5193 | 0.135 | 10743 | 0.234 |
| 3 | 1.929 | 0.097 | 3295 | 0.085 | 8985 | 0.196 |
| Total | | | 3855422 | 100.000 | 4588826 | 100.000 |

### NMR and HPLC spectra of 2b:

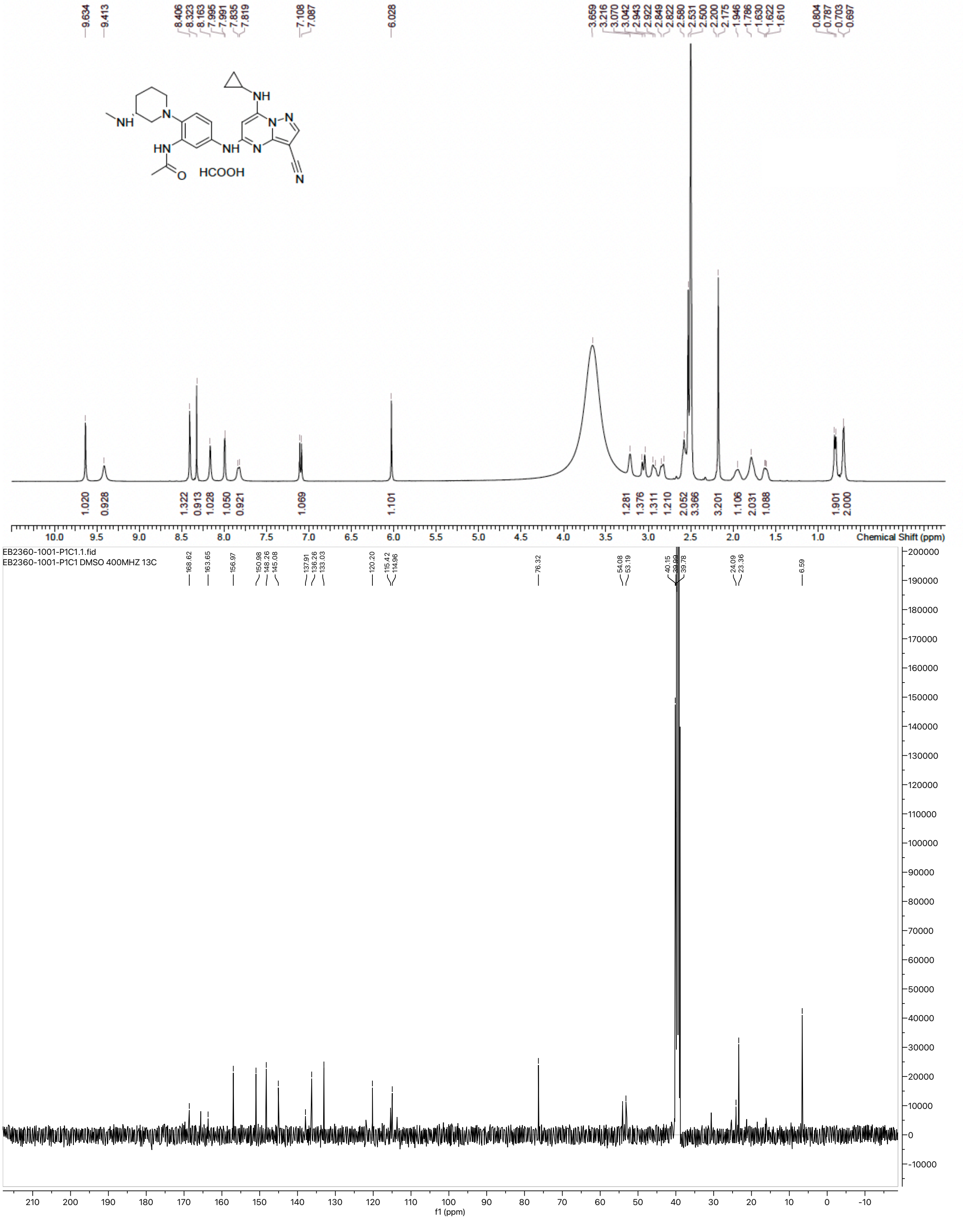

<Chromatogram>

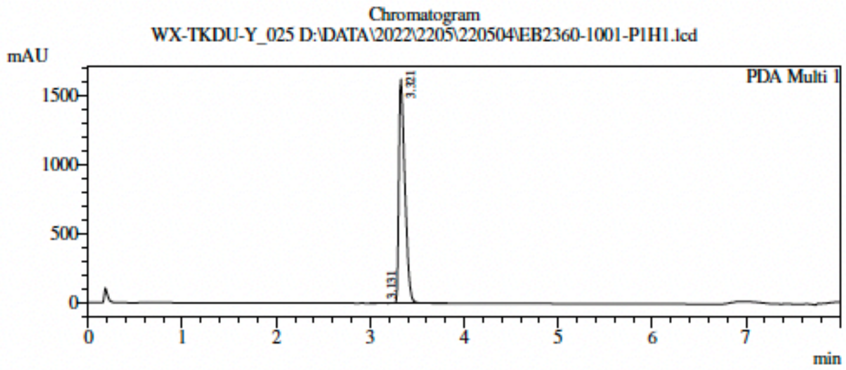


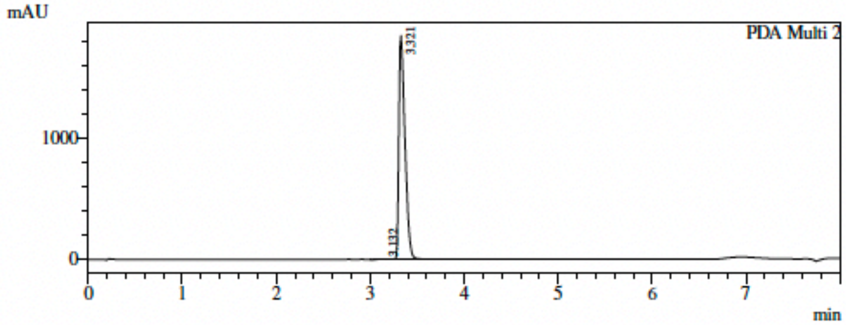


1 PDA Multi 1 / 220nm 4nm
2 PDA Multi 2 / 254nm 4nm

Integration Result

PeakTable

Ch1 220nm 4nm

| Peak# | Ret. Time | Height | Height % | Area | Area % |
|---|---|---|---|---|---|
| 1 | 3.131 | 1033 | 0.064 | 6654 | 0.091 |
| 2 | 3.321 | 1622252 | 99.936 | 7297195 | 99.909 |
| Total | | 1623285 | 100.000 | 7303848 | 100.000 |

PeakTable

Ch2 254nm 4nm

| Peak# | Ret. Time | Height | Height % | Area | Area % |
|---|---|---|---|---|---|
| 1 | 3.132 | 1380 | 0.074 | 8612 | 0.103 |
| 2 | 3.321 | 1854411 | 99.926 | 8393331 | 99.897 |
| Total | | 1855790 | 100.000 | 8401943 | 100.000 |

PeakTable

Ch3

### NMR and HPLC spectra of 2c:

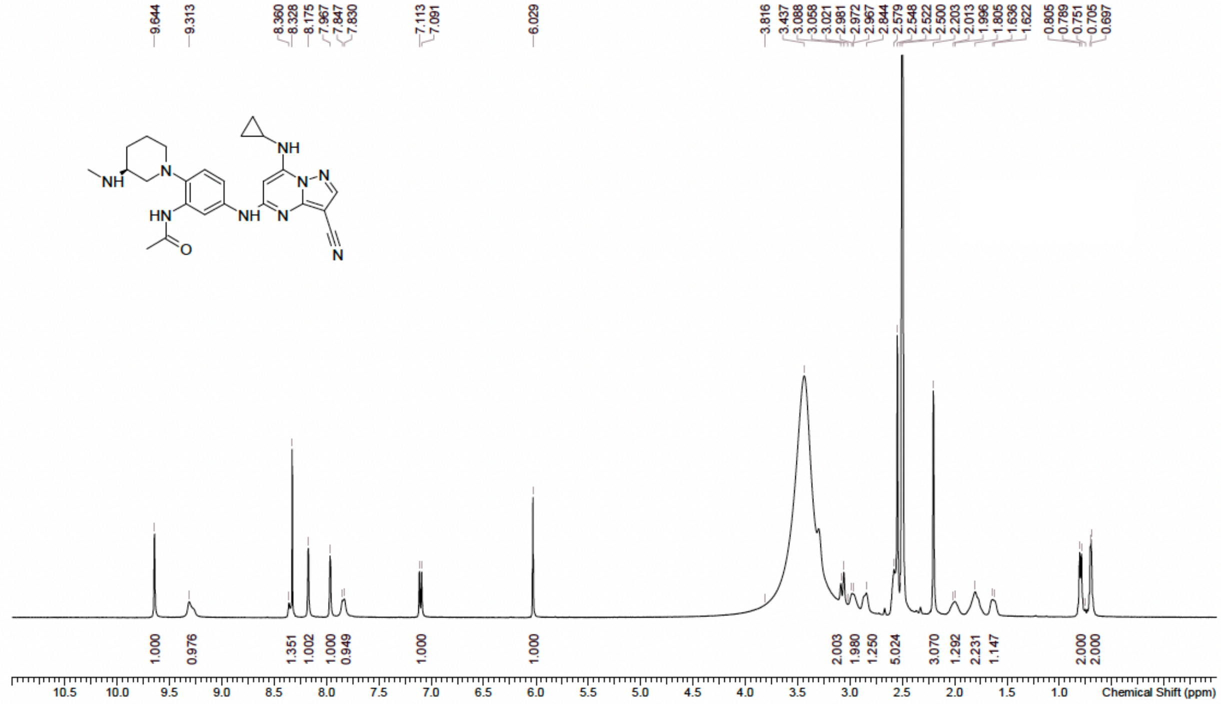


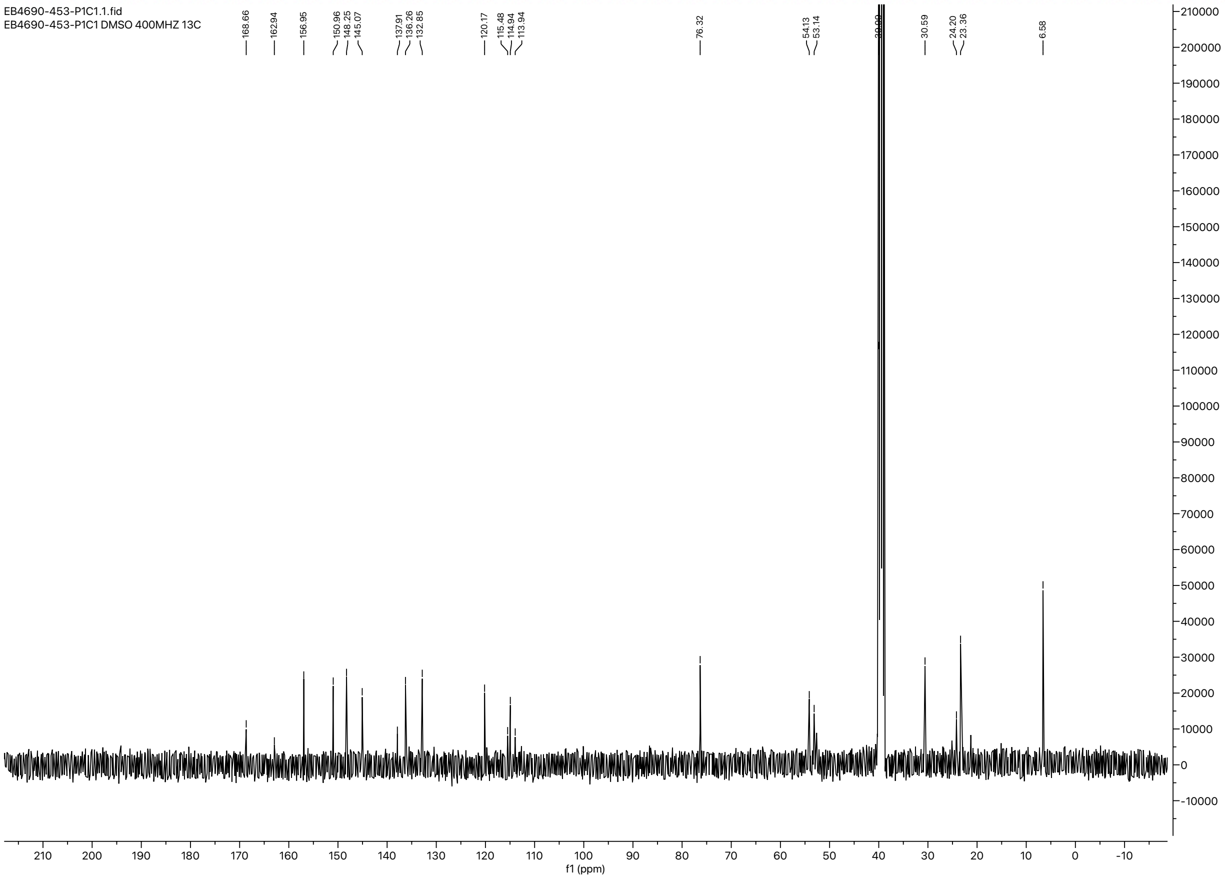

**<Chromatogram>**

Chromatogram
WX-TKDU-Y_026 D:\DATA\2022\2205\220505\EB4690-453-P1H1.lcd

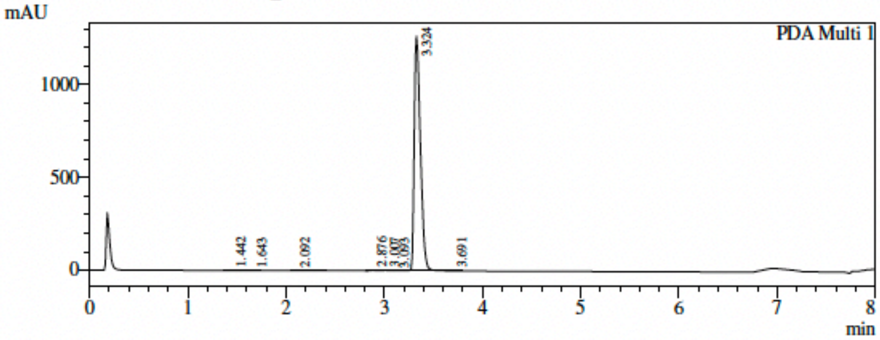


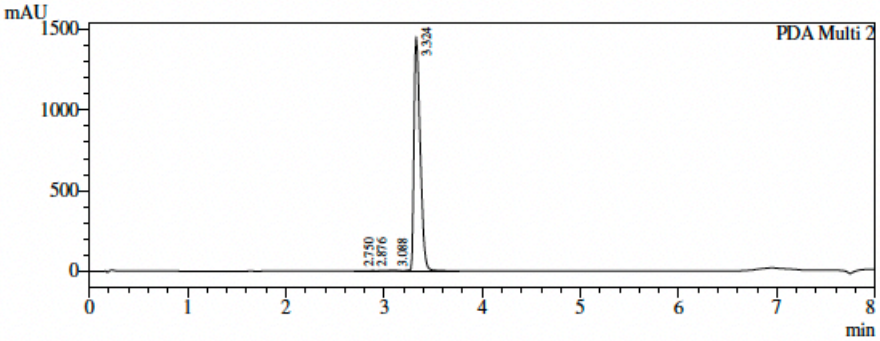


1 PDA Multi 1 / 220nm 4nm
2 PDA Multi 2 / 254nm 4nm

Integration Result

| Peak# | Ret. Time | Height | Height % | Area | Area % |
|---|---|---|---|---|---|
| 1 | 1.442 | 669 | 0.052 | 3821 | 0.068 |
| 2 | 1.643 | 904 | 0.071 | 3964 | 0.071 |
| 3 | 2.092 | 936 | 0.073 | 4694 | 0.084 |
| 4 | 2.876 | 2818 | 0.221 | 8102 | 0.145 |
| 5 | 3.007 | 2896 | 0.227 | 11392 | 0.204 |
| 6 | 3.093 | 2306 | 0.181 | 9965 | 0.178 |
| 7 | 3.324 | 1262348 | 99.123 | 5553022 | 99.206 |
| 8 | 3.691 | 642 | 0.050 | 2505 | 0.045 |
| Total | | 1273520 | 100.000 | 5597466 | 100.000 |

PeakTable

Ch2 254nm 4nm

| Peak# | Ret. Time | Height | Height % | Area | Area % |
|---|---|---|---|---|---|
| 1 | 2.750 | 704 | 0.048 | 1976 | 0.031 |
| 2 | 2.876 | 2048 | 0.141 | 5918 | 0.092 |
| 3 | 3.088 | 2344 | 0.161 | 17110 | 0.266 |
| 4 | 3.324 | 1452495 | 99.650 | 6407337 | 99.611 |
| Total | | 1457591 | 100.000 | 6432342 | 100.000 |

PeakTable

Ch3

### NMR and HPLC spectra of 2d:

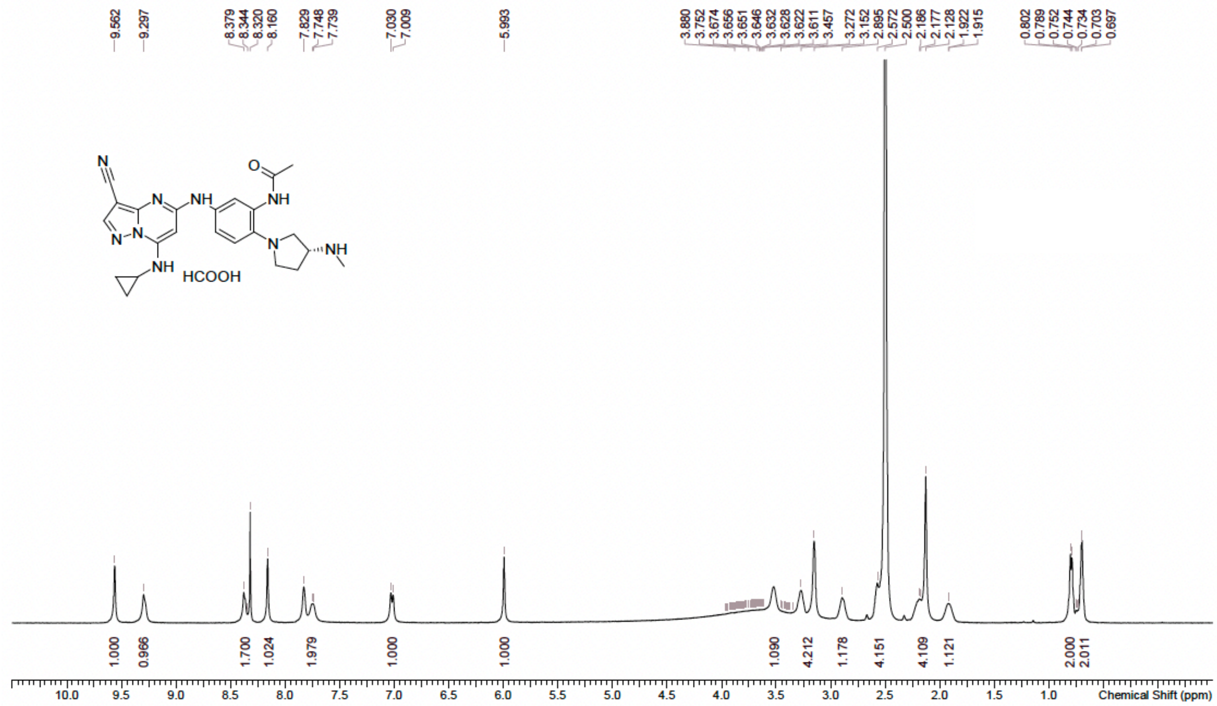


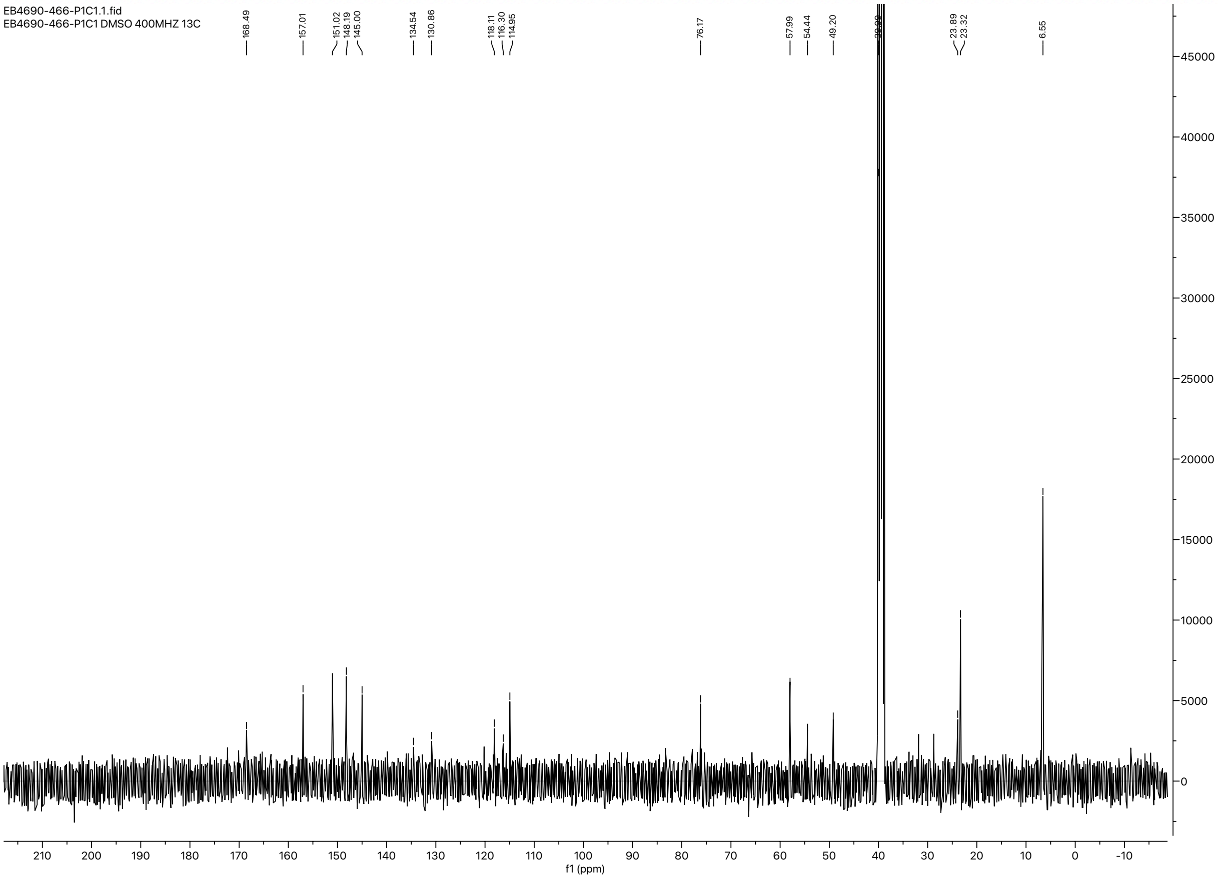

<Chromatogram>

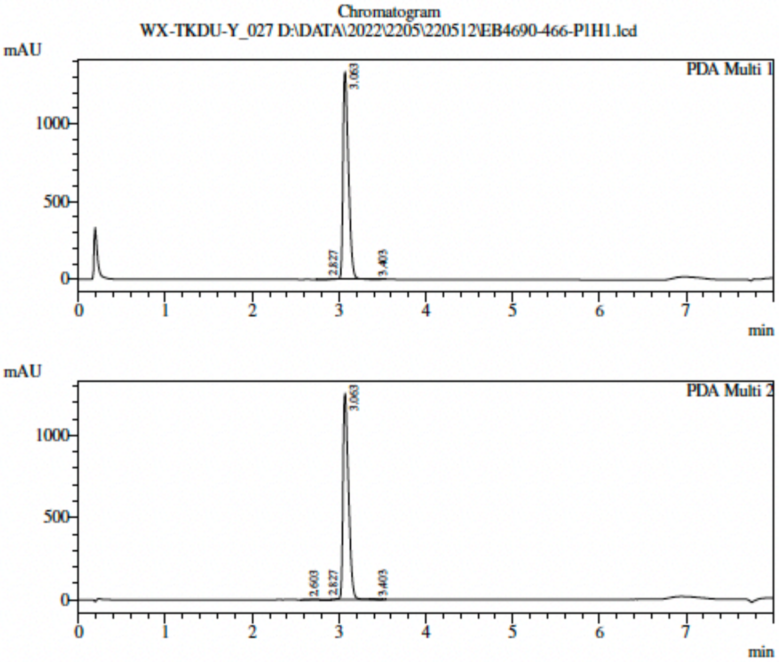


1 PDA Multi 1 / 220nm 4nm
2 PDA Multi 2 / 254nm 4nm

Integration Result

PeakTable

Ch1 220nm 4nm

| Peak# | Ret. Time | Height | Height % | Area | Area % |
|---|---|---|---|---|---|
| 1 | 2.827 | 537 | 0.040 | 3741 | 0.064 |
| 2 | 3.063 | 1335188 | 99.896 | 5817773 | 99.882 |
| 3 | 3.403 | 847 | 0.063 | 3116 | 0.054 |
| Total | | 1336572 | 100.000 | 5824631 | 100.000 |

PeakTable

Ch2 254nm 4nm

| Peak# | Ret. Time | Height | Height % | Area | Area % |
|---|---|---|---|---|---|
| 1 | 2.603 | 311 | 0.025 | 2263 | 0.041 |
| 2 | 2.827 | 596 | 0.047 | 4191 | 0.076 |
| 3 | 3.063 | 1254815 | 99.850 | 5523486 | 99.812 |
| 4 | 3.403 | 981 | 0.078 | 3952 | 0.071 |
| Total | | 1256703 | 100.000 | 5533892 | 100.000 |

PeakTable

Ch3

### NMR and HPLC spectra of 2e:

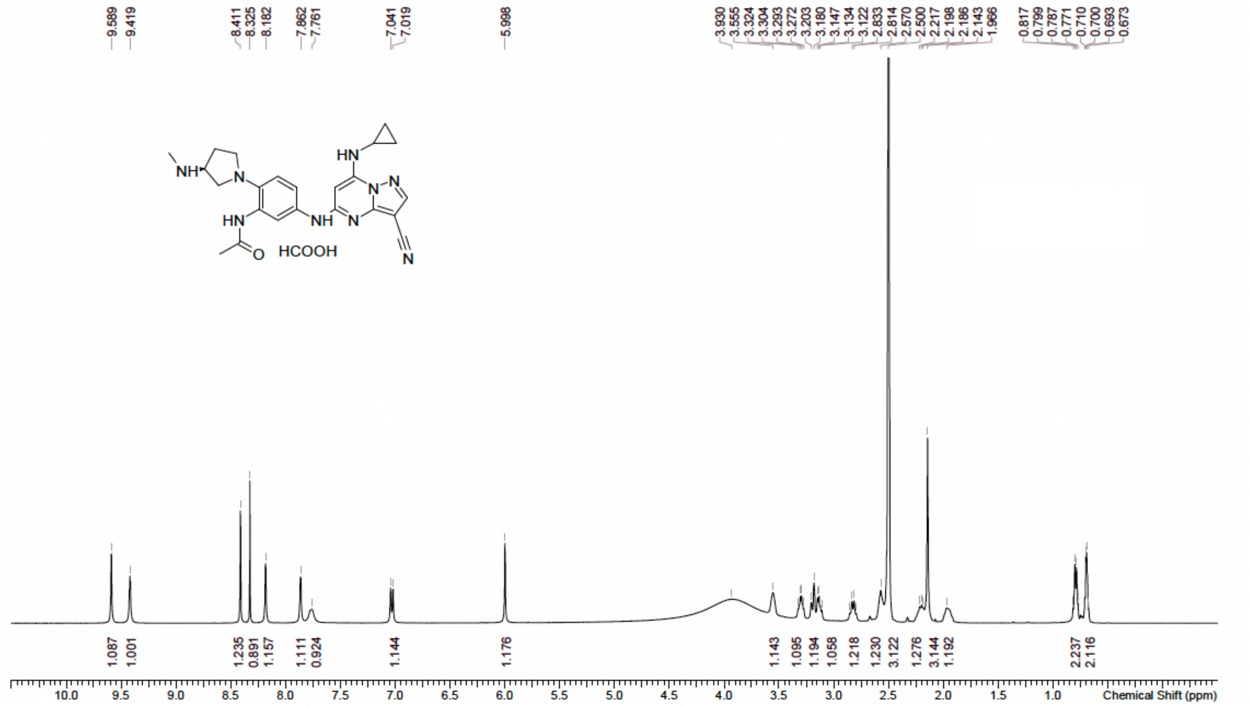


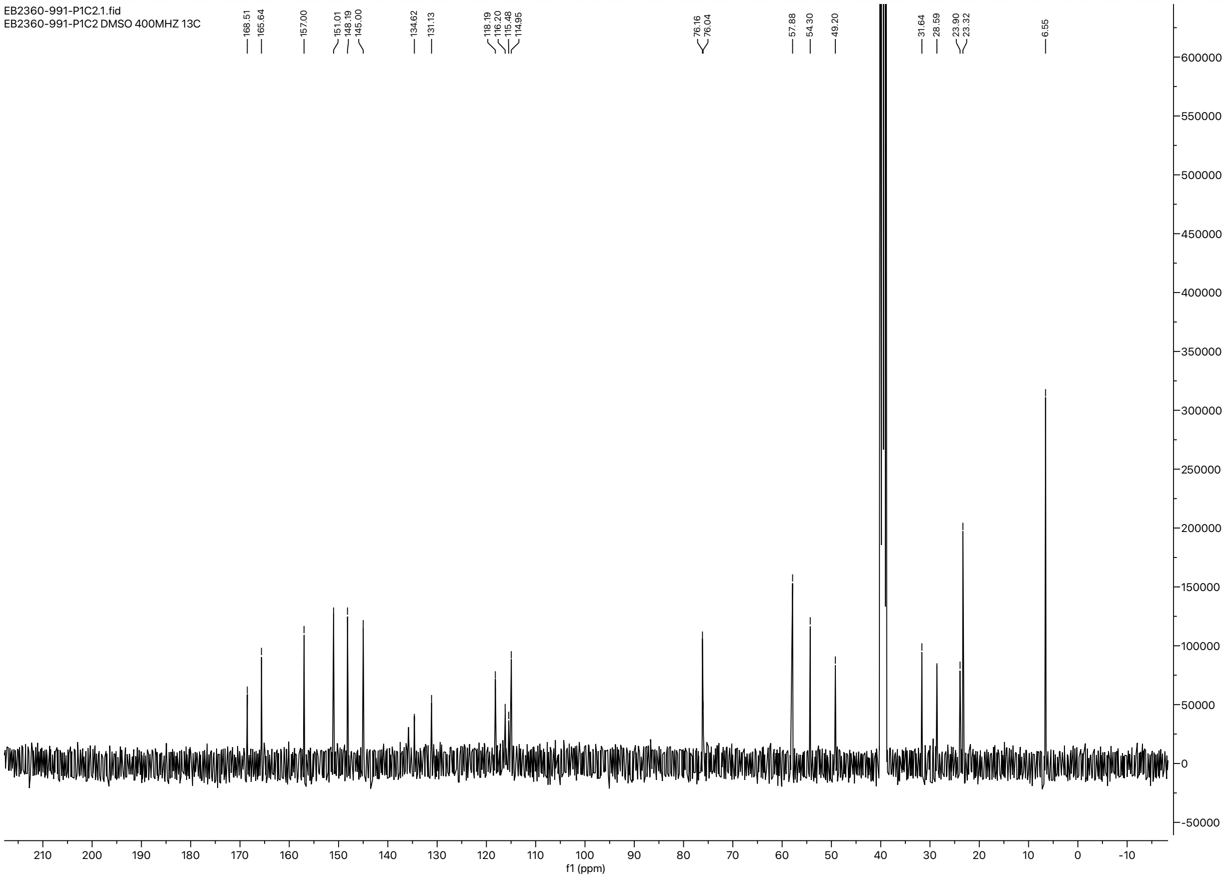

<Chromatogram>

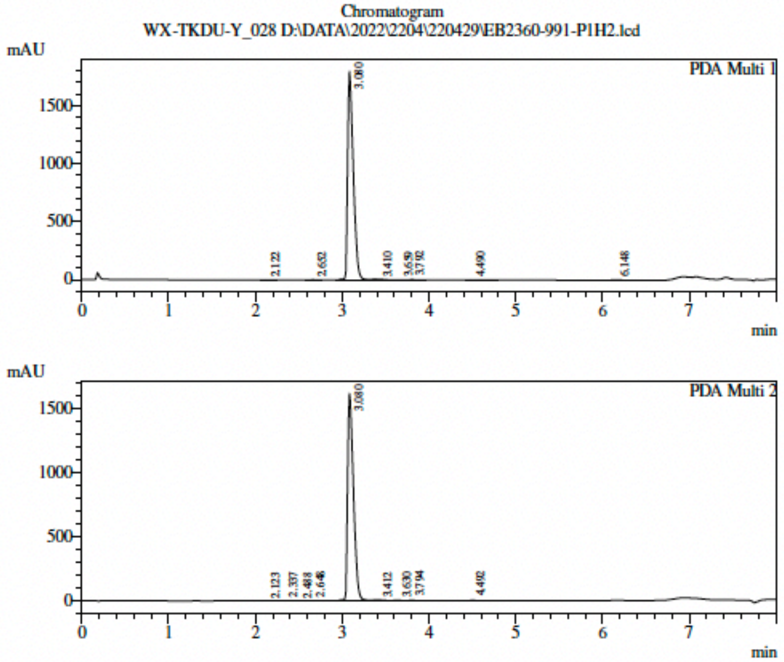


1 PDA Multi 1 / 220nm 4nm
2 PDA Multi 2 / 254nm 4nm

Integration Result

PeakTable

Ch1 220nm 4nm

| Peak# | Ret. Time | Height | Height % | Area | Area % |
|---|---|---|---|---|---|
| 1 | 2.122 | 879 | 0.049 | 3332 | 0.041 |
| 2 | 2.652 | 1488 | 0.082 | 6243 | 0.076 |
| 3 | 3.080 | 1795487 | 99.464 | 8126700 | 99.409 |
| 4 | 3.410 | 1132 | 0.063 | 4311 | 0.053 |
| 5 | 3.659 | 503 | 0.028 | 3100 | 0.038 |
| 6 | 3.792 | 1757 | 0.097 | 6196 | 0.076 |
| 7 | 4.490 | 1052 | 0.058 | 7485 | 0.092 |
| 8 | 6.148 | 2873 | 0.159 | 17664 | 0.216 |
| Total | | 1805171 | 100.000 | 8175032 | 100.000 |

PeakTable

Ch2 254nm 4nm

| Peak# | Ret. Time | Height | Height % | Area | Area % |
|---|---|---|---|---|---|
| 1 | 2.123 | 367 | 0.023 | 1856 | 0.024 |
| 2 | 2.337 | 647 | 0.040 | 2470 | 0.032 |
| 3 | 2.488 | 678 | 0.042 | 2791 | 0.037 |
| 4 | 2.648 | 1147 | 0.071 | 5862 | 0.077 |
| 5 | 3.080 | 1618889 | 99.554 | 7573209 | 99.586 |
| 6 | 3.412 | 1503 | 0.092 | 5688 | 0.075 |
| 7 | 3.630 | 616 | 0.038 | 3590 | 0.047 |
| 8 | 3.794 | 1405 | 0.086 | 4974 | 0.065 |
| 9 | 4.492 | 886 | 0.054 | 4223 | 0.056 |
| Total | | 1626137 | 100.000 | 7604665 | 100.000 |

PeakTable

Ch3

### NMR and HPLC spectra of 2f:

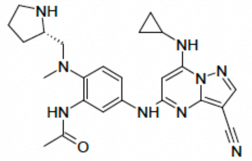

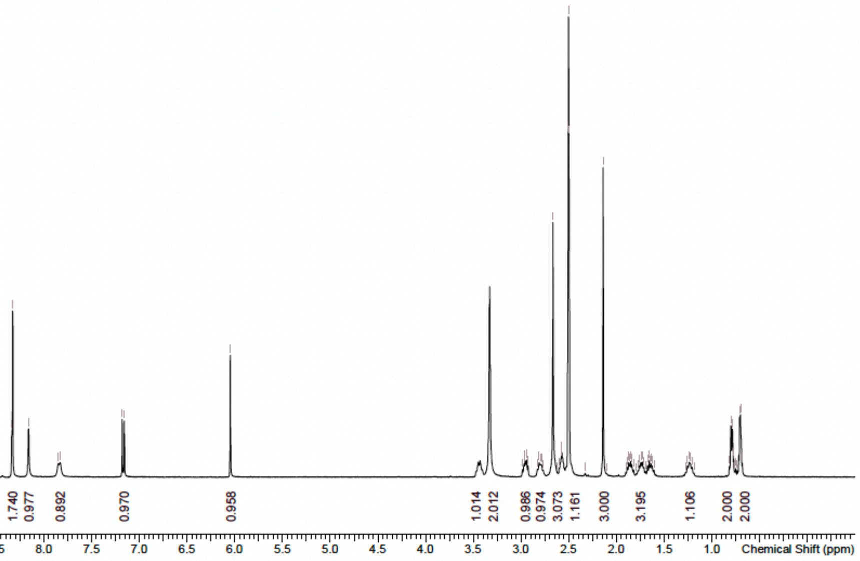


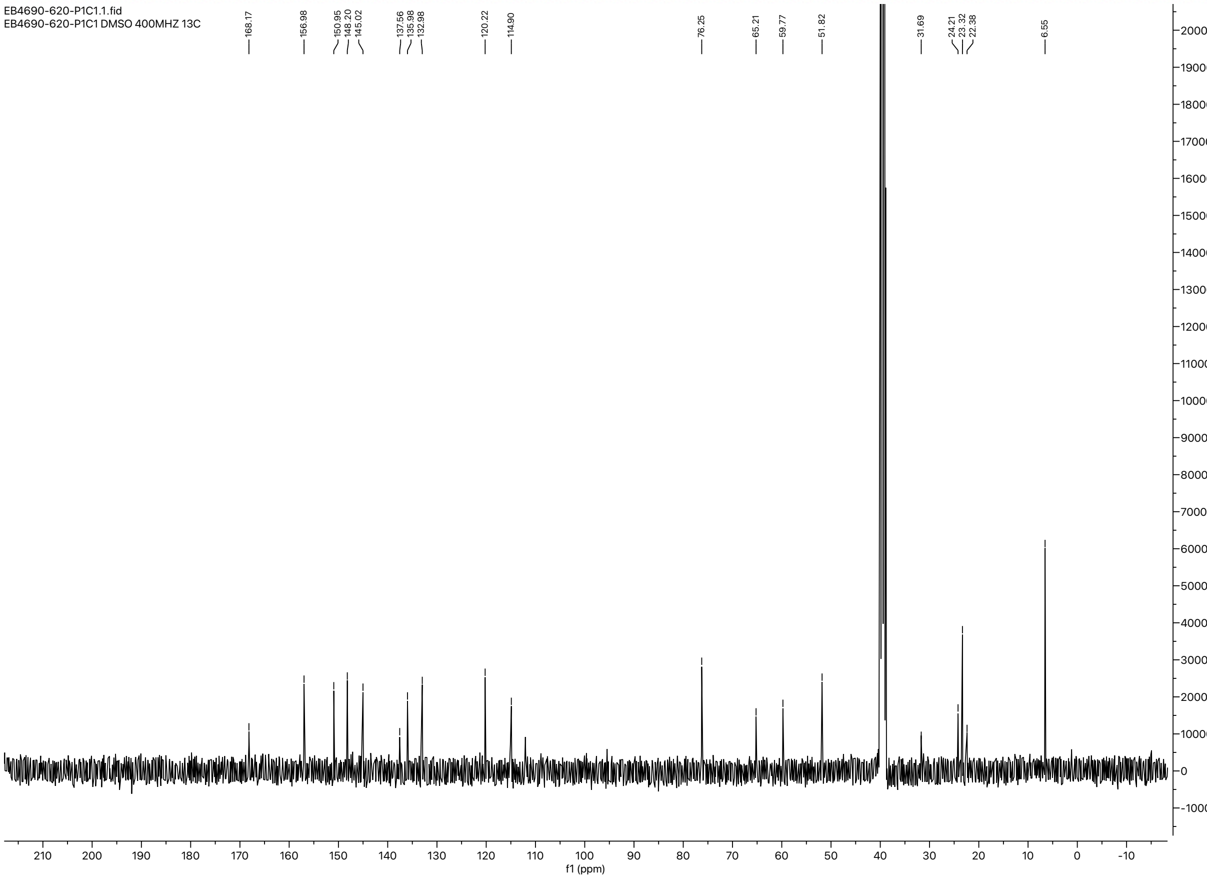

<Chromatogram>

Chromatogram
WX-TKDU-G_032 D:\DATA\2022\2209\220923\EB4690-634-P1H1.lcd

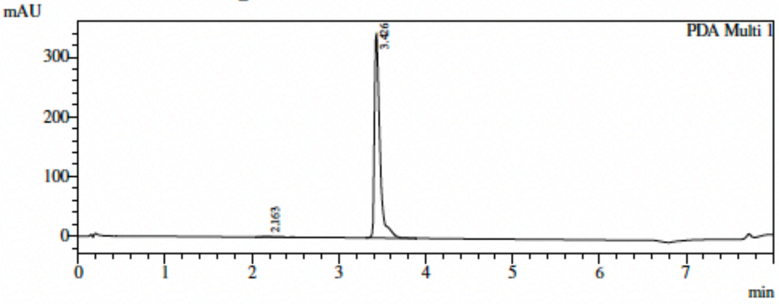


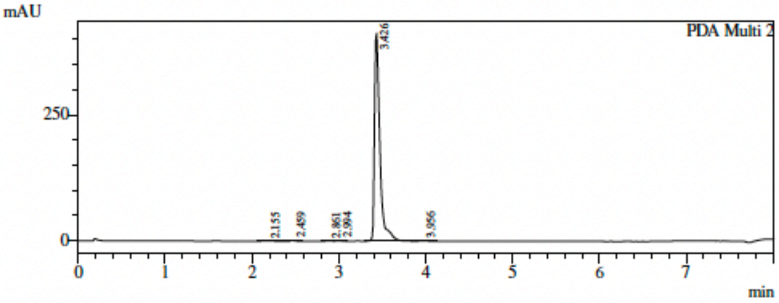


1 PDA Multi 1 / 220nm 4nm
2 PDA Multi 2 / 254nm 4nm

Integration Result

PeakTable

PDA Ch1 220nm 4nm

| Peak# | Ret. Time | Height | Height % | Area | Area % |
|---|---|---|---|---|---|
| 1 | 2.163 | 939 | 0.273 | 9684 | 0.645 |
| 2 | 3.426 | 343510 | 99.727 | 1491336 | 99.355 |
| Total | | 344450 | 100.000 | 1501019 | 100.000 |

PeakTable

PDA Ch2 254nm 4nm

| Peak# | Ret. Time | Height | Height % | Area | Area % |
|---|---|---|---|---|---|
| 1 | 2.155 | 550 | 0.132 | 4596 | 0.254 |
| 2 | 2.459 | 1237 | 0.297 | 3627 | 0.200 |
| 3 | 2.861 | 741 | 0.178 | 2560 | 0.141 |
| 4 | 2.994 | 573 | 0.137 | 2121 | 0.117 |
| 5 | 3.426 | 412783 | 99.069 | 1792504 | 99.025 |
| 6 | 3.956 | 777 | 0.186 | 4750 | 0.262 |
| Total | | 416661 | 100.000 | 1810158 | 100.000 |

PeakTable

PDA Ch3

### NMR and HPLC spectra of 2g:

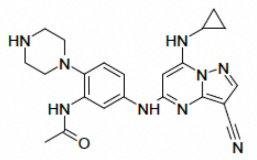

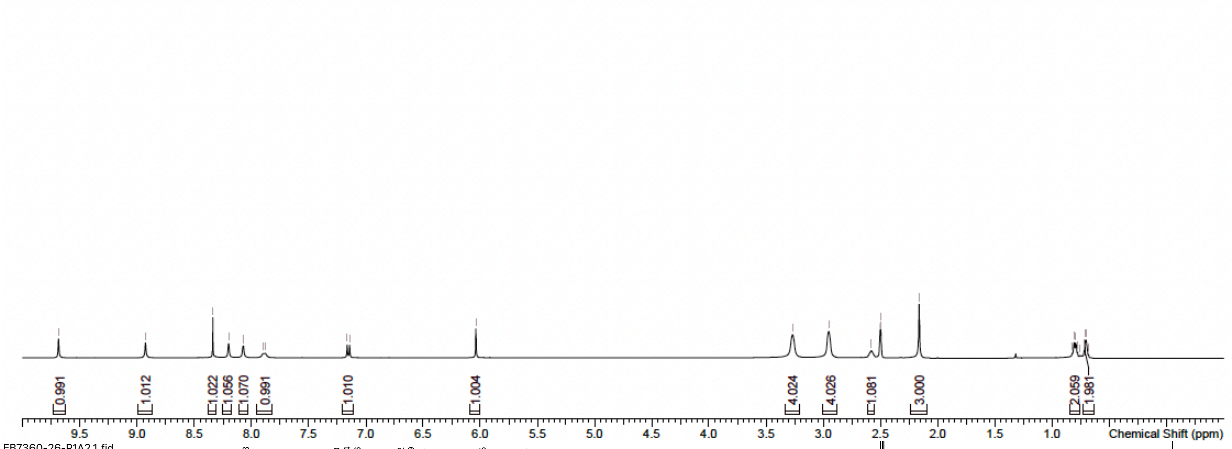


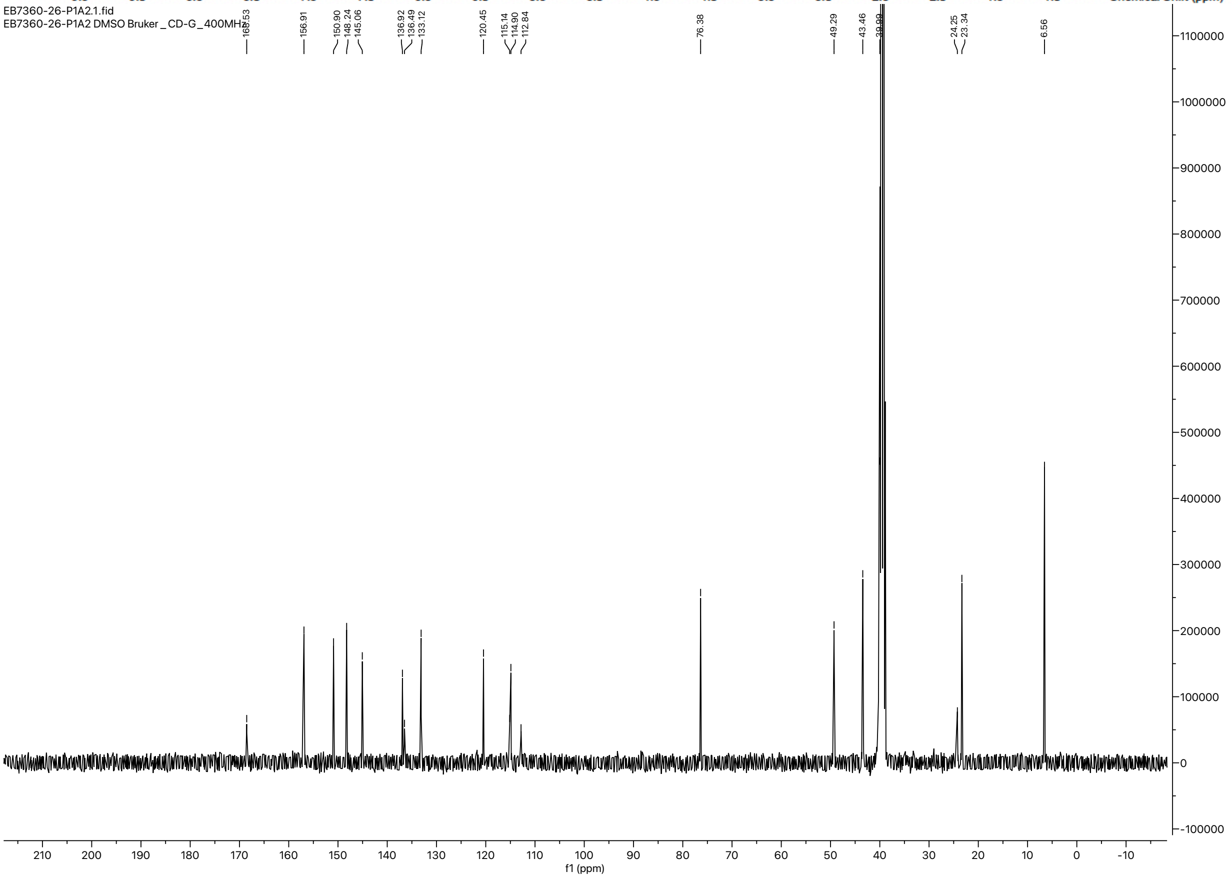

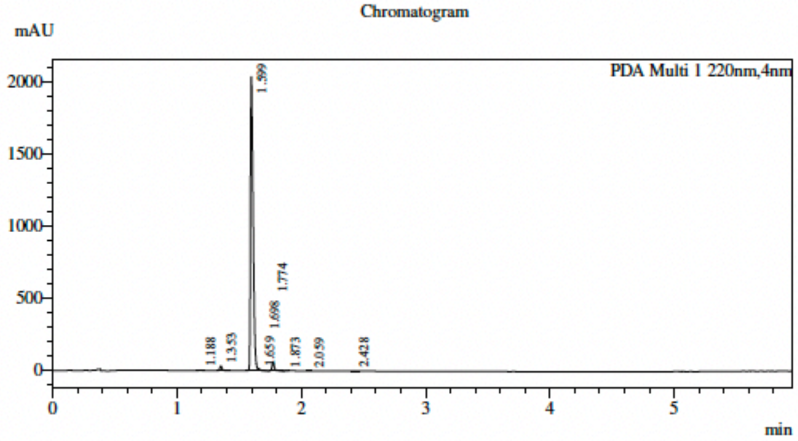


Integration Result

PDA Ch1 220nm

| Peak# | Ret. Time | USP Width | Height | Height% | Area | Area% |
|---|---|---|---|---|---|---|
| 1 | 1.188 | 0.029 | 2840 | 0.132 | 3255 | 0.102 |
| 2 | 1.353 | 0.031 | 31659 | 1.471 | 39398 | 1.233 |
| 3 | 1.599 | 0.040 | 2040631 | 94.802 | 3056550 | 95.644 |
| 4 | 1.659 | 0.346 | 10296 | 0.478 | 11494 | 0.360 |
| 5 | 1.698 | 0.154 | 2754 | 0.128 | 6020 | 0.188 |
| 6 | 1.774 | 0.031 | 59167 | 2.749 | 72616 | 2.272 |
| 7 | 1.873 | 0.033 | 2499 | 0.116 | 2954 | 0.092 |
| 8 | 2.059 | 0.030 | 1139 | 0.053 | 1324 | 0.041 |
| 9 | 2.428 | 0.037 | 1527 | 0.071 | 2137 | 0.067 |
| Total | | | 2152512 | 100.000 | 3195748 | 100.000 |

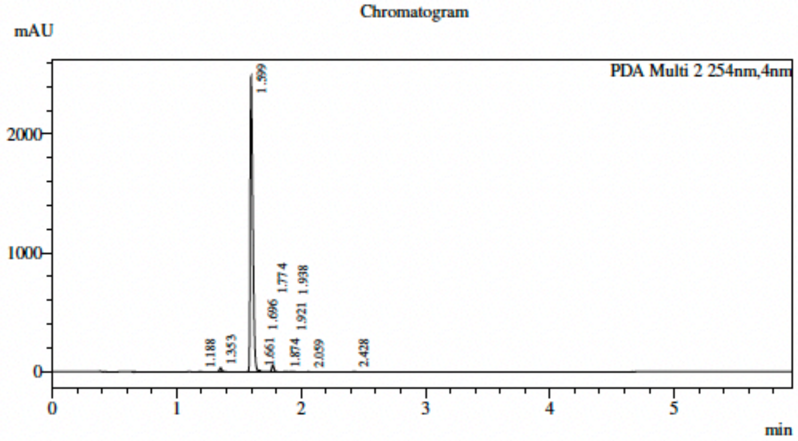


Integration Result

PDA Ch2 254nm

| Peak# | Ret. Time | USP Width | Height | Height% | Area | Area% |
|---|---|---|---|---|---|---|
| 1 | 1.188 | 0.029 | 2055 | 0.079 | 2348 | 0.061 |
| 2 | 1.353 | 0.031 | 33133 | 1.273 | 41448 | 1.082 |
| 3 | 1.599 | 0.040 | 2483078 | 95.403 | 3680072 | 96.064 |
| 4 | 1.661 | 0.132 | 13669 | 0.525 | 15936 | 0.416 |
| 5 | 1.696 | 0.103 | 3076 | 0.118 | 5824 | 0.152 |
| 6 | 1.774 | 0.031 | 57759 | 2.219 | 71882 | 1.876 |
| 7 | 1.874 | 0.038 | 2793 | 0.107 | 4013 | 0.105 |
| 8 | 1.921 | 0.071 | 2304 | 0.089 | 3326 | 0.087 |
| 9 | 1.938 | 0.178 | 2026 | 0.078 | 2223 | 0.058 |
| 10 | 2.059 | 0.031 | 1067 | 0.041 | 1231 | 0.032 |
| 11 | 2.428 | 0.038 | 1775 | 0.068 | 2539 | 0.066 |
| Total | | | 2602737 | 100.000 | 3830842 | 100.000 |

### NMR and HPLC spectra of 2i:

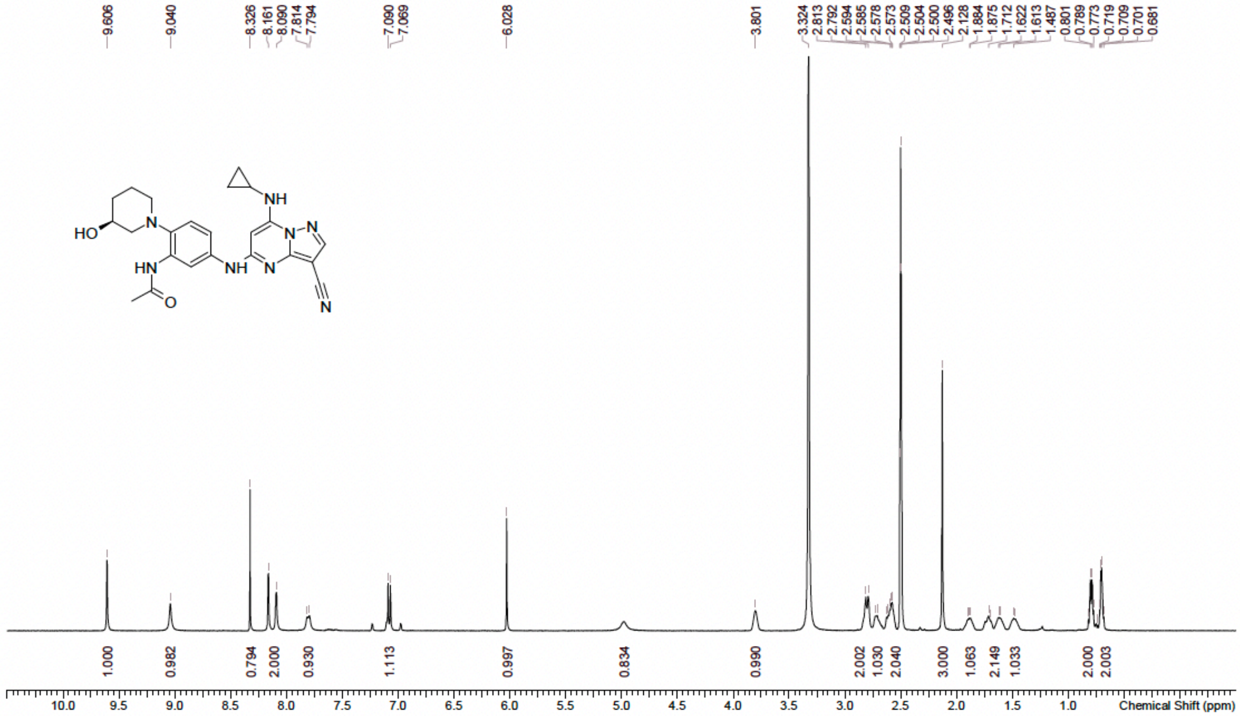


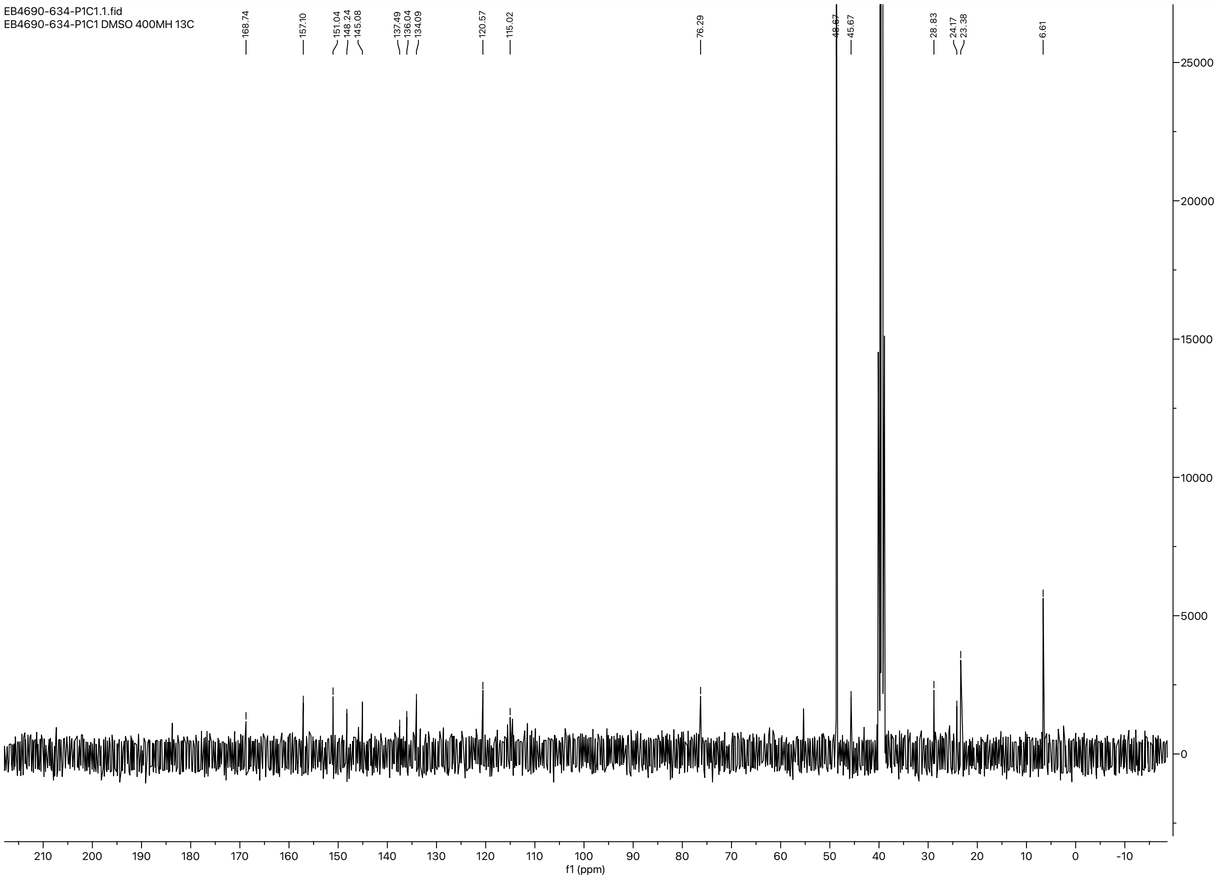

[illegible]

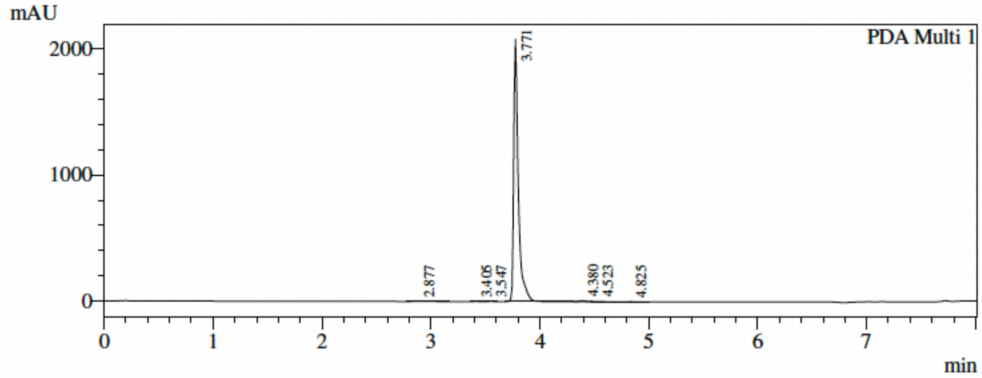


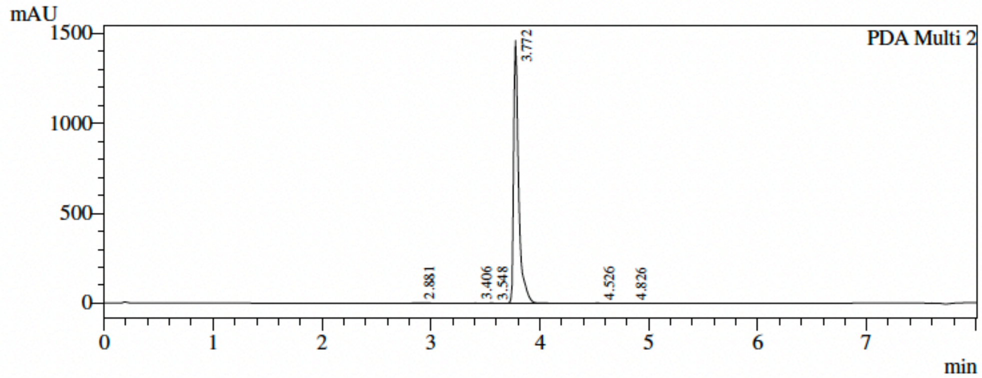


1 PDA Multi 1 / 220nm 4nm
2 PDA Multi 2 / 254nm 4nm

==========================================================================

Integration Result

==========================================================================

PeakTable

PDA Ch1 220nm 4nm

| Peak# | Ret. Time | Height | Height % | Area | Area % |
|---|---|---|---|---|---|
| 1 | 2.877 | 1163 | 0.056 | 12629 | 0.206 |
| 2 | 3.405 | 1255 | 0.060 | 3405 | 0.056 |
| 3 | 3.547 | 1153 | 0.055 | 3183 | 0.052 |
| 4 | 3.771 | 2077942 | 99.328 | 6071865 | 98.995 |
| 5 | 4.380 | 7986 | 0.382 | 30941 | 0.504 |
| 6 | 4.523 | 1775 | 0.085 | 8301 | 0.135 |
| 7 | 4.825 | 725 | 0.035 | 3158 | 0.051 |
| Total | | 2091998 | 100.000 | 6133482 | 100.000 |

PeakTable

PDA Ch2 254nm 4nm

| Peak# | Ret. Time | Height | Height % | Area | Area % |
|---|---|---|---|---|---|
| 1 | 2.881 | 1347 | 0.092 | 15542 | 0.330 |
| 2 | 3.406 | 768 | 0.052 | 2115 | 0.045 |
| 3 | 3.548 | 1061 | 0.072 | 2893 | 0.061 |
| 4 | 3.772 | 1459960 | 99.699 | 4685327 | 99.465 |
| 5 | 4.526 | 647 | 0.044 | 2310 | 0.049 |
| 6 | 4.826 | 584 | 0.040 | 2325 | 0.049 |
| Total | | 1464367 | 100.000 | 4710512 | 100.000 |

PeakTable

PDA Ch3

### NMR and HPLC spectra of 2j:

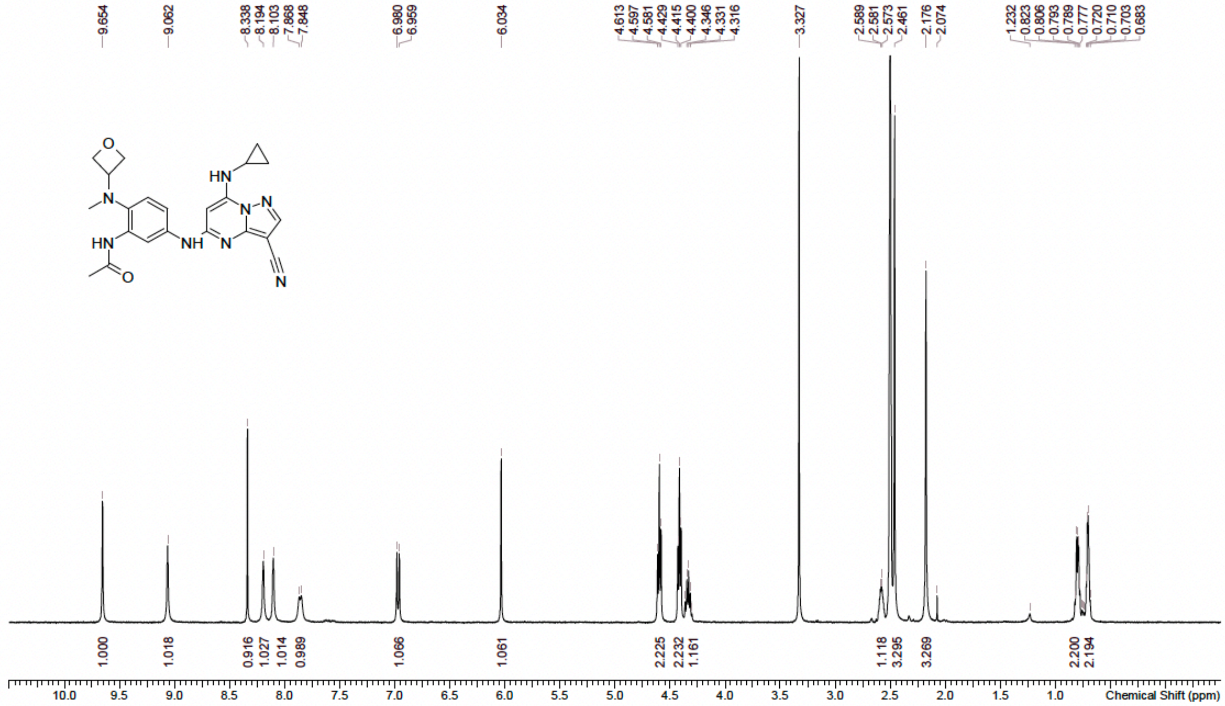


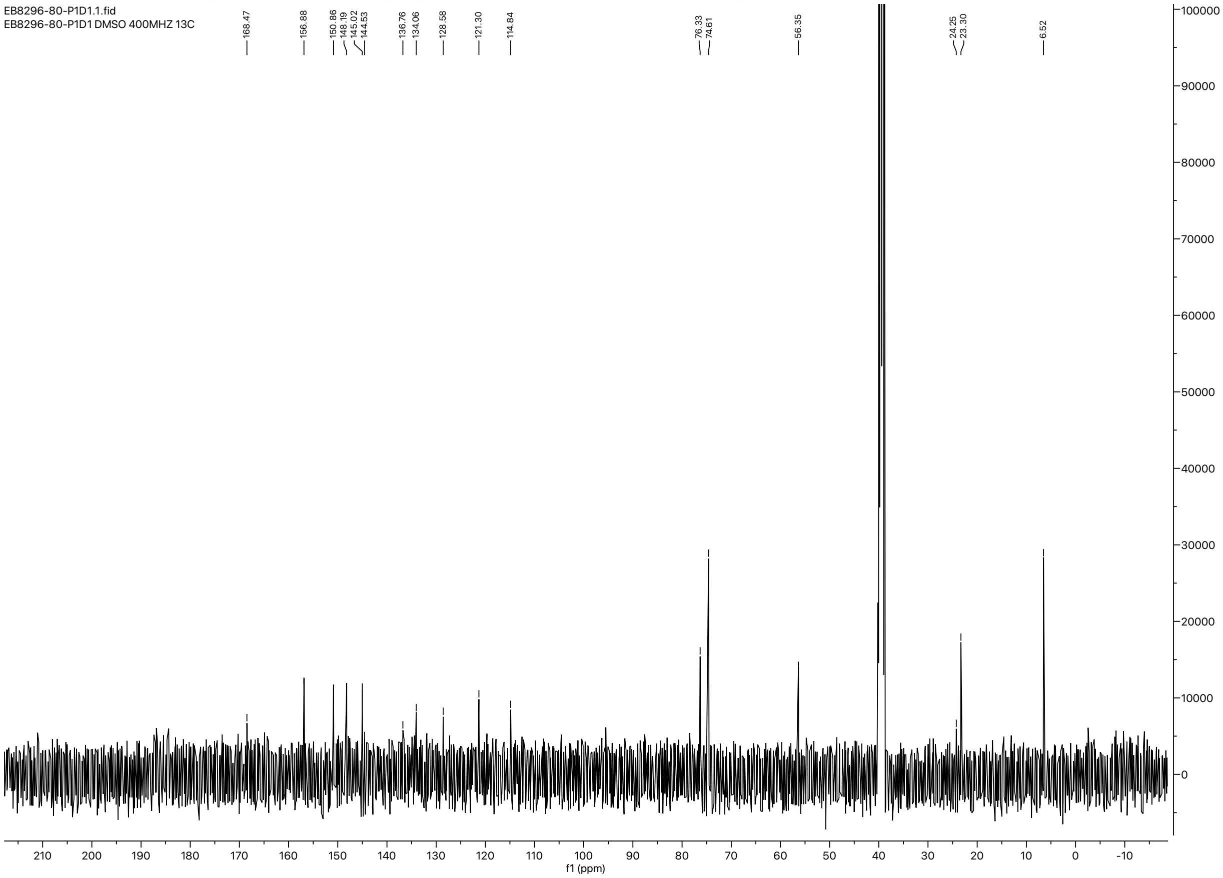

[illegible]

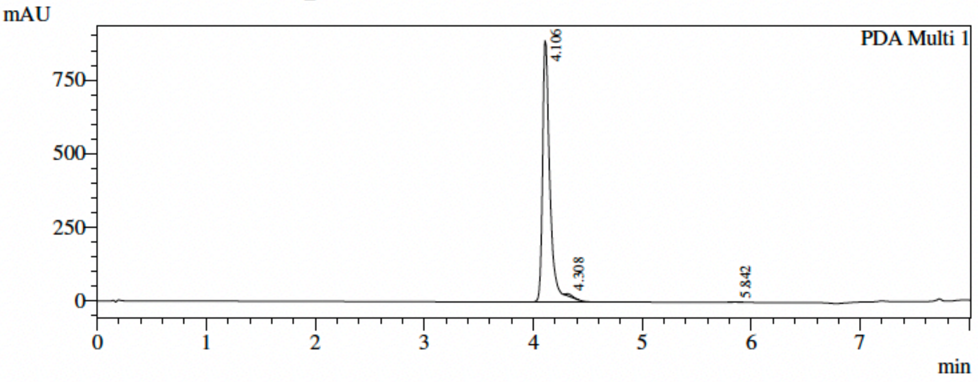


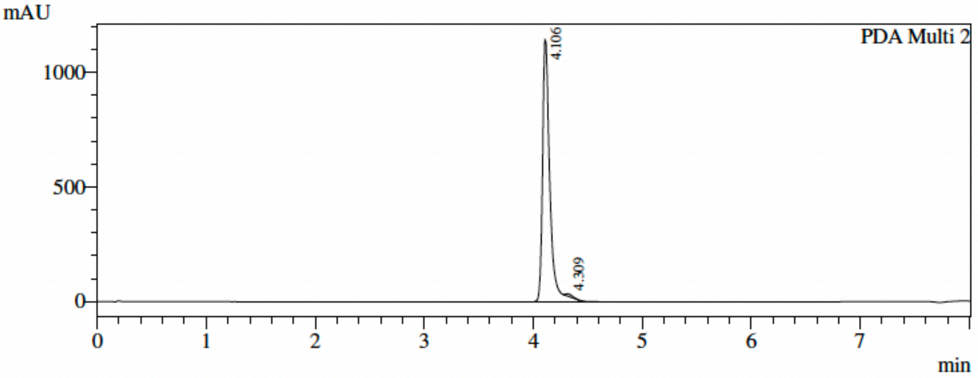


1 PDA Multi 1 / 220nm 4nm
2 PDA Multi 2 / 254nm 4nm

==========================================================================

Integration Result

==========================================================================

PeakTable

PDA Ch1 220nm 4nm

| Peak# | Ret. Time | Height | Height % | Area | Area % |
|---|---|---|---|---|---|
| 1 | 4.106 | 886833 | 99.264 | 4203036 | 99.335 |
| 2 | 4.308 | 5995 | 0.671 | 25736 | 0.608 |
| 3 | 5.842 | 583 | 0.065 | 2384 | 0.056 |
| Total | | 893411 | 100.000 | 4231157 | 100.000 |

PeakTable

PDA Ch2 254nm 4nm

| Peak# | Ret. Time | Height | Height % | Area | Area % |
|---|---|---|---|---|---|
| 1 | 4.106 | 1143554 | 99.264 | 5337382 | 99.318 |
| 2 | 4.309 | 8480 | 0.736 | 36639 | 0.682 |
| Total | | 1152034 | 100.000 | 5374021 | 100.000 |

PeakTable

PDA Ch3

### NMR and HPLC spectra of 2k:

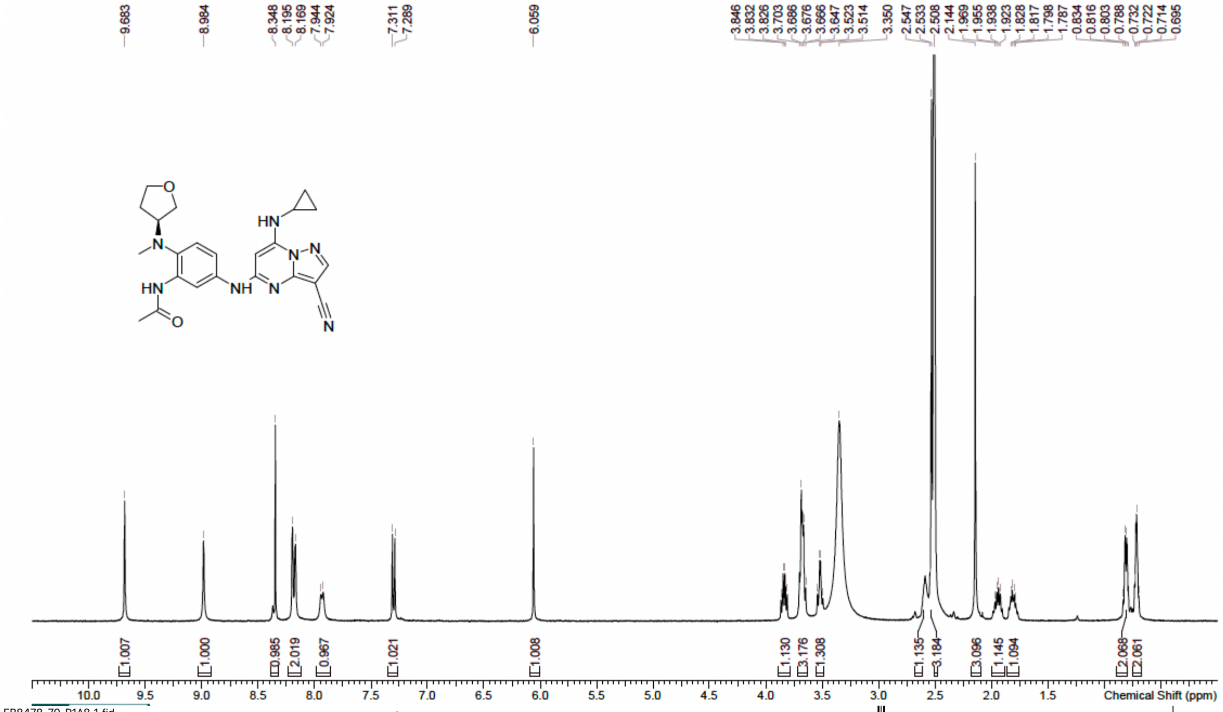

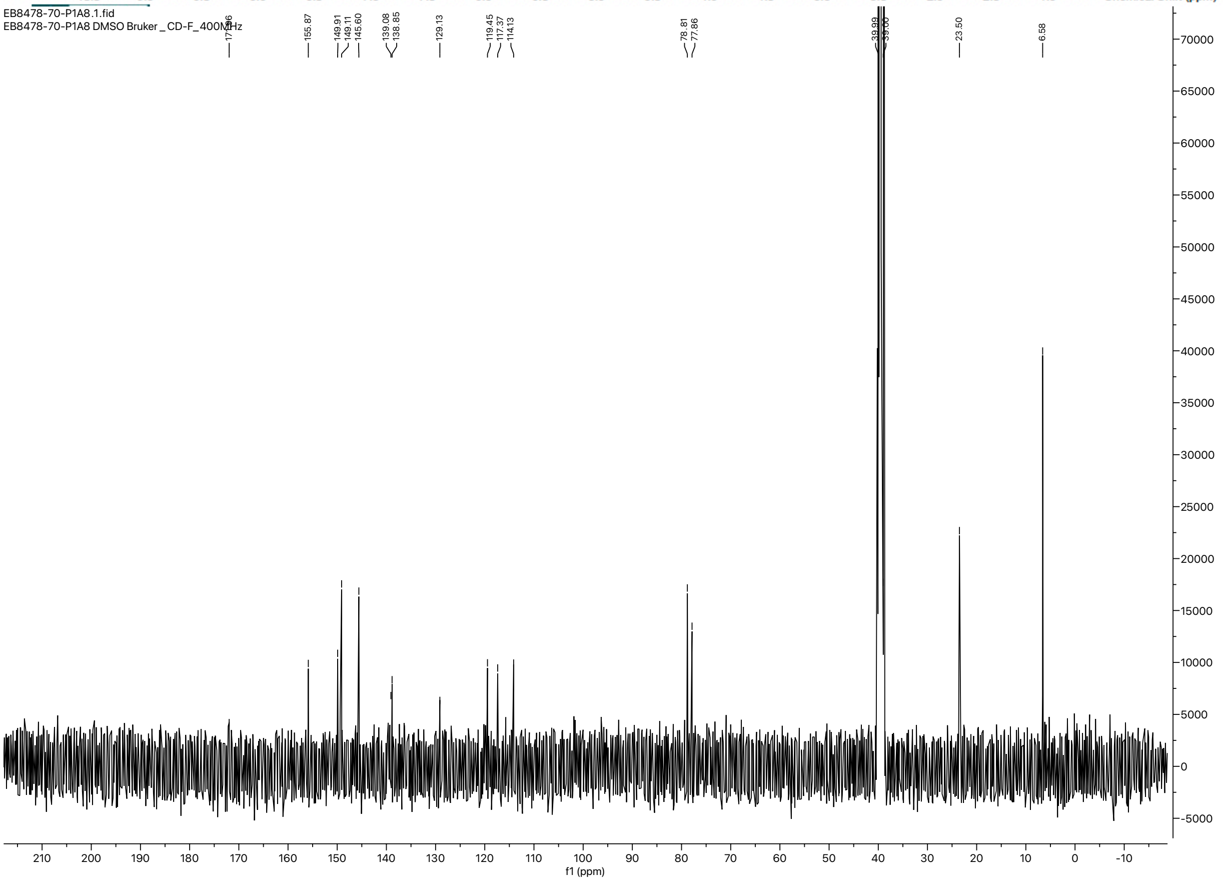

Chromatogram

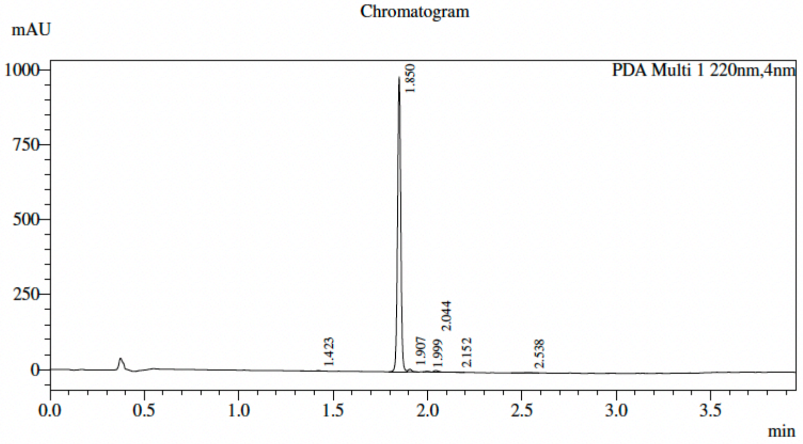


### Integration Result

PDA Ch1 220nm

| Peak# | Ret. Time | USP Width | Height | Height% | Area | Area% |
|---|---|---|---|---|---|---|
| 1 | 1.423 | 0.027 | 1466 | 0.146 | 1475 | 0.122 |
| 2 | 1.850 | 0.031 | 985161 | 97.960 | 1174514 | 97.374 |
| 3 | 1.907 | 0.037 | 8679 | 0.863 | 11739 | 0.973 |
| 4 | 1.999 | 0.032 | 2189 | 0.218 | 2584 | 0.214 |
| 5 | 2.044 | 0.043 | 5527 | 0.550 | 9142 | 0.758 |
| 6 | 2.152 | 0.109 | 1245 | 0.124 | 1800 | 0.149 |
| 7 | 2.538 | 0.105 | 1407 | 0.140 | 4939 | 0.409 |
| Total | | | 1005673 | 100.000 | 1206194 | 100.000 |

Chromatogram

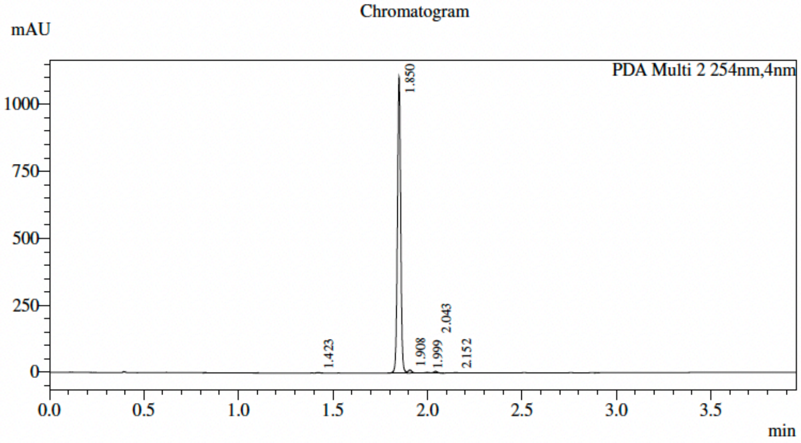


### Integration Result

PDA Ch2 254nm

| Peak# | Ret. Time | USP Width | Height | Height% | Area | Area% |
|---|---|---|---|---|---|---|
| 1 | 1.423 | 0.028 | 2106 | 0.187 | 2149 | 0.160 |
| 2 | 1.850 | 0.031 | 1106064 | 98.001 | 1316884 | 97.824 |
| 3 | 1.908 | 0.036 | 11203 | 0.993 | 15210 | 1.130 |
| 4 | 1.999 | 0.033 | 2199 | 0.195 | 2634 | 0.196 |
| 5 | 2.043 | 0.032 | 5955 | 0.528 | 7441 | 0.553 |
| 6 | 2.152 | 0.040 | 1102 | 0.098 | 1854 | 0.138 |
| Total | | | 1128629 | 100.000 | 1346173 | 100.000 |

**NMR and HPLC spectra of 2l:**

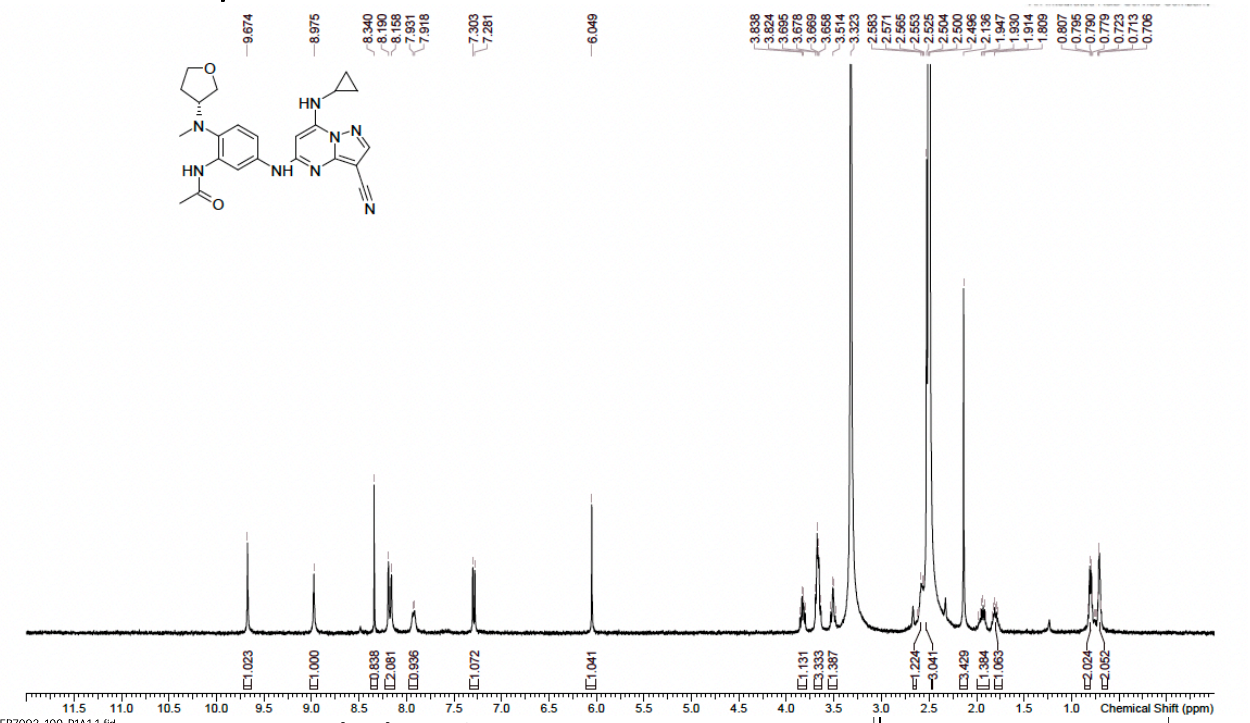


EB7093-100-P1A1.1.fid
EB7093-100-P1A1 DMSO Bruker _CD-C_400MHz 13C

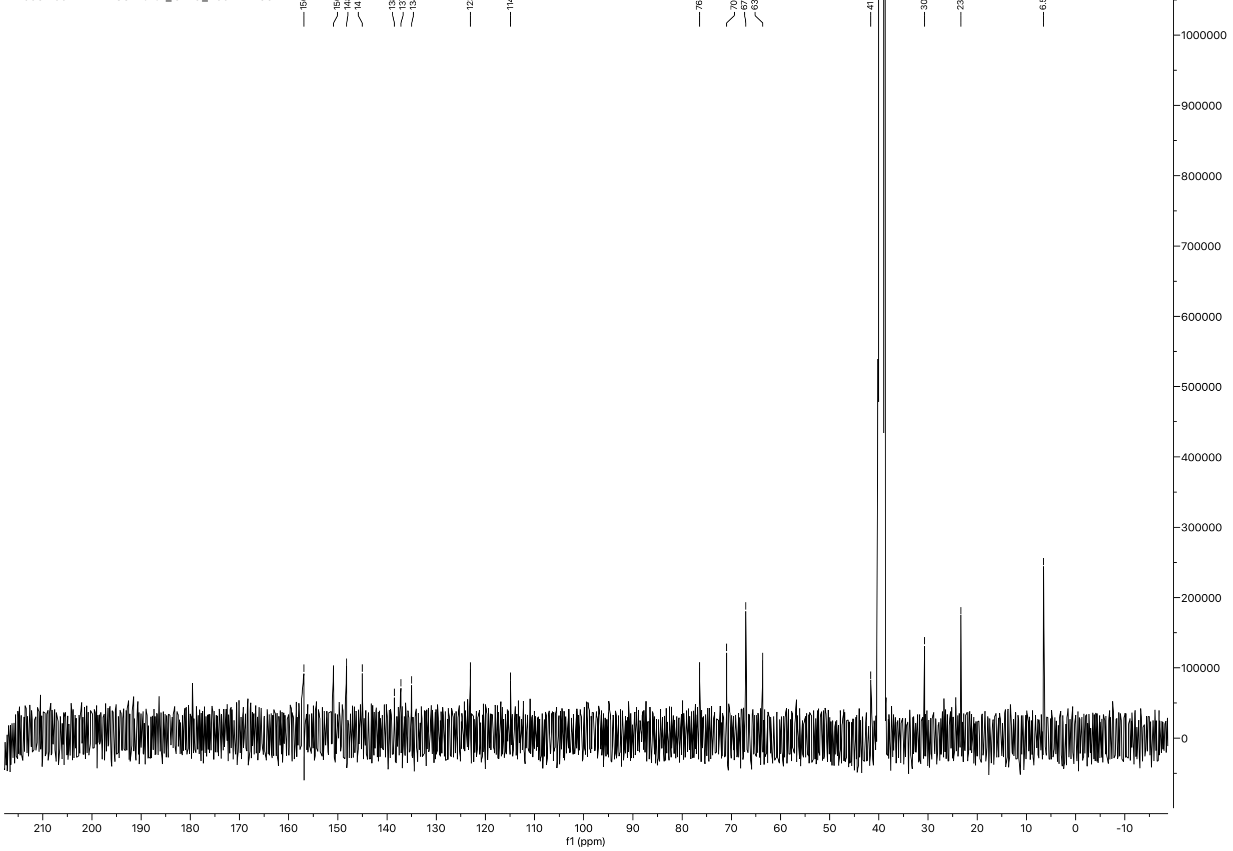

Chromatogram

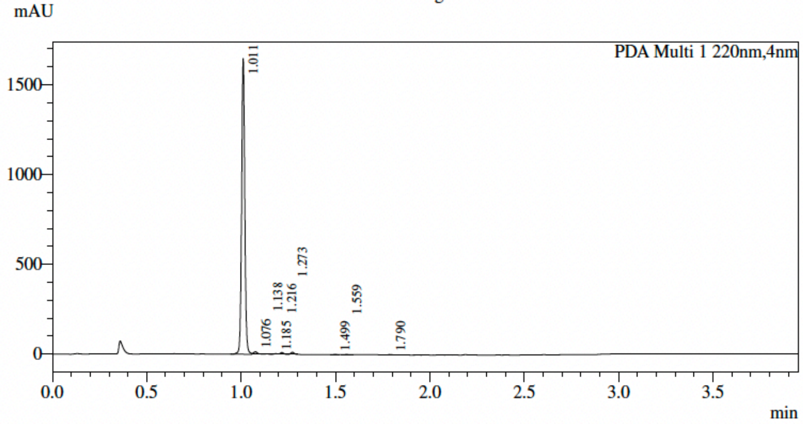


**Integration Result**

PDA Ch1 220nm

| Peak# | Ret. Time | USP Width | Height | Height% | Area | Area% |
|---|---|---|---|---|---|---|
| 1 | 1.011 | 0.030 | 1647850 | 97.064 | 1930550 | 96.685 |
| 2 | 1.076 | 0.040 | 13718 | 0.808 | 21639 | 1.084 |
| 3 | 1.138 | 0.035 | 2810 | 0.166 | 3740 | 0.187 |
| 4 | 1.185 | 0.036 | 4091 | 0.241 | 5236 | 0.262 |
| 5 | 1.216 | 0.034 | 10313 | 0.607 | 13133 | 0.658 |
| 6 | 1.273 | 0.033 | 11686 | 0.688 | 14086 | 0.705 |
| 7 | 1.499 | 0.032 | 2509 | 0.148 | 3055 | 0.153 |
| 8 | 1.559 | 0.032 | 2420 | 0.143 | 3133 | 0.157 |
| 9 | 1.790 | 0.025 | 2291 | 0.135 | 2180 | 0.109 |
| Total | | | 1697688 | 100.000 | 1996752 | 100.000 |

Chromatogram

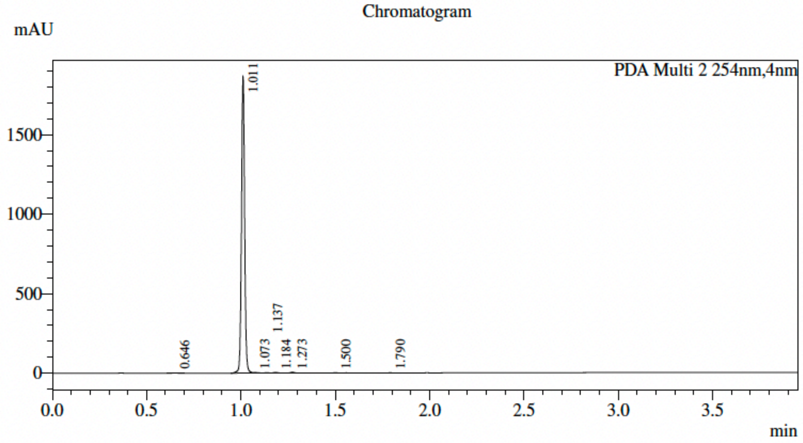


**Integration Result**

PDA Ch2 254nm

| Peak# | Ret. Time | USP Width | Height | Height% | Area | Area% |
|---|---|---|---|---|---|---|
| 1 | 0.646 | 0.040 | 1654 | 0.088 | 2754 | 0.125 |
| 2 | 1.011 | 0.030 | 1863169 | 98.883 | 2179383 | 98.756 |
| 3 | 1.073 | 0.061 | 4213 | 0.224 | 6923 | 0.314 |
| 4 | 1.137 | 0.030 | 2226 | 0.118 | 2587 | 0.117 |
| 5 | 1.184 | 0.032 | 4371 | 0.232 | 5208 | 0.236 |
| 6 | 1.273 | 0.033 | 5366 | 0.285 | 6454 | 0.292 |
| 7 | 1.500 | 0.031 | 2156 | 0.114 | 2586 | 0.117 |
| 8 | 1.790 | 0.024 | 1053 | 0.056 | 943 | 0.043 |
| Total | | | 1884208 | 100.000 | 2206839 | 100.000 |

### NMR and HPLC spectra of 2m:

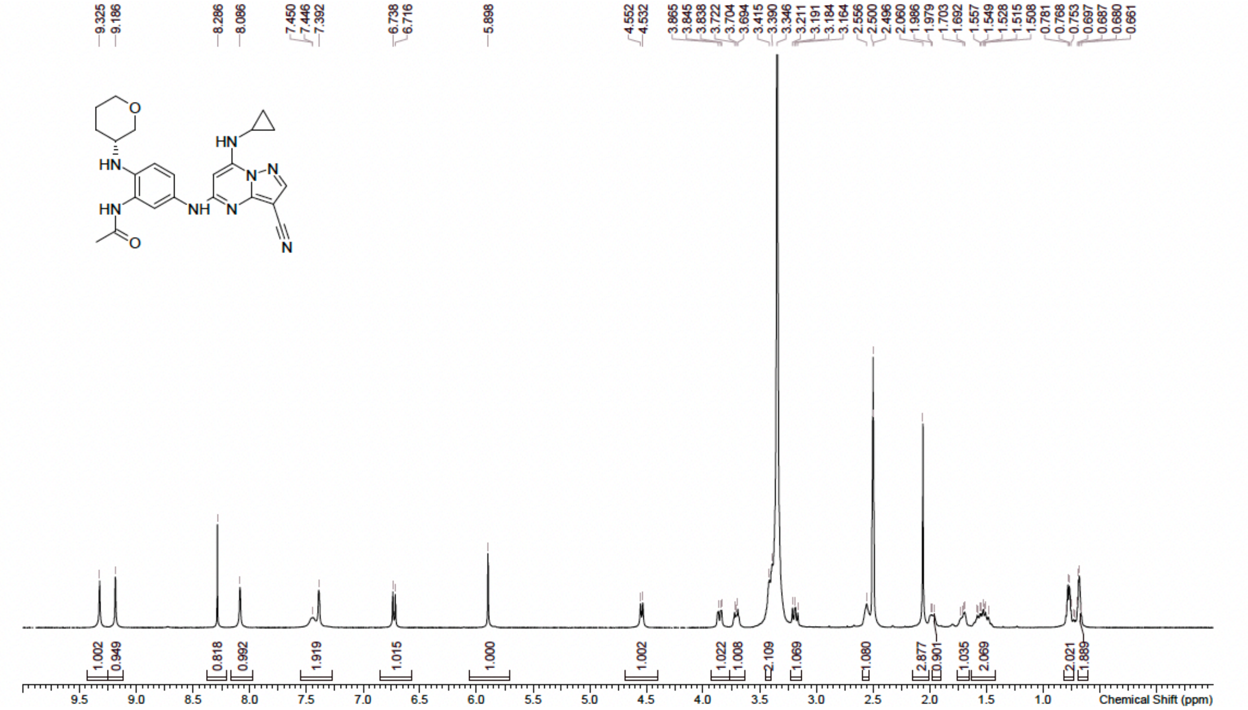


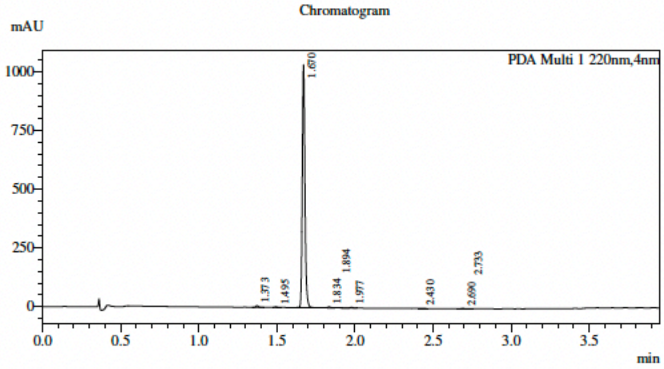


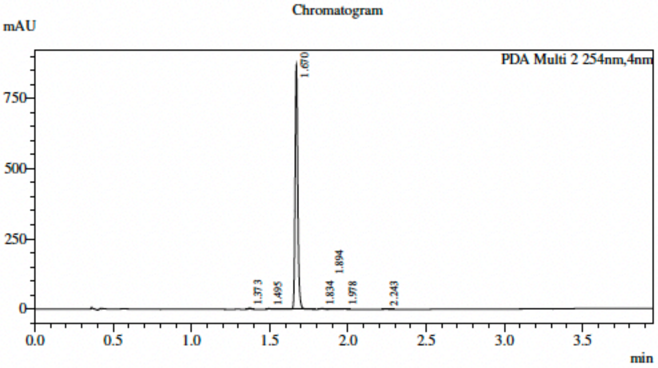


**Integration Result**

PDA Ch1 220nm

| Peak# | Ret. Time | USP Width | Height | Height% | Area | Area% |
|---|---|---|---|---|---|---|
| 1 | 1.373 | 0.028 | 7323 | 0.693 | 8110 | 0.653 |
| 2 | 1.495 | 0.031 | 2975 | 0.281 | 3892 | 0.314 |
| 3 | 1.670 | 0.030 | 1034702 | 97.890 | 1211514 | 97.599 |
| 4 | 1.834 | 0.032 | 4392 | 0.416 | 5782 | 0.466 |
| 5 | 1.894 | 0.041 | 1102 | 0.104 | 1807 | 0.146 |
| 6 | 1.977 | 0.034 | 2849 | 0.270 | 4428 | 0.357 |
| 7 | 2.430 | 0.042 | 1239 | 0.117 | 1969 | 0.159 |
| 8 | 2.690 | 0.037 | 1362 | 0.129 | 2147 | 0.173 |
| 9 | 2.733 | 0.041 | 1055 | 0.100 | 1674 | 0.135 |
| Total | | | 1057000 | 100.000 | 1241322 | 100.000 |

**Integration Result**

PDA Ch2 254nm

| Peak# | Ret. Time | USP Width | Height | Height% | Area | Area% |
|---|---|---|---|---|---|---|
| 1 | 1.373 | 0.028 | 4165 | 0.470 | 4449 | 0.427 |
| 2 | 1.495 | 0.031 | 2984 | 0.337 | 5124 | 0.492 |
| 3 | 1.670 | 0.030 | 871955 | 98.352 | 1021765 | 98.038 |
| 4 | 1.834 | 0.031 | 3351 | 0.378 | 4028 | 0.386 |
| 5 | 1.894 | 0.042 | 1278 | 0.144 | 2881 | 0.276 |
| 6 | 1.978 | 0.035 | 1812 | 0.204 | 2479 | 0.238 |
| 7 | 2.243 | 0.037 | 1024 | 0.116 | 1488 | 0.143 |
| Total | | | 886569 | 100.000 | 1042213 | 100.000 |

### NMR and HPLC spectra of 2n:

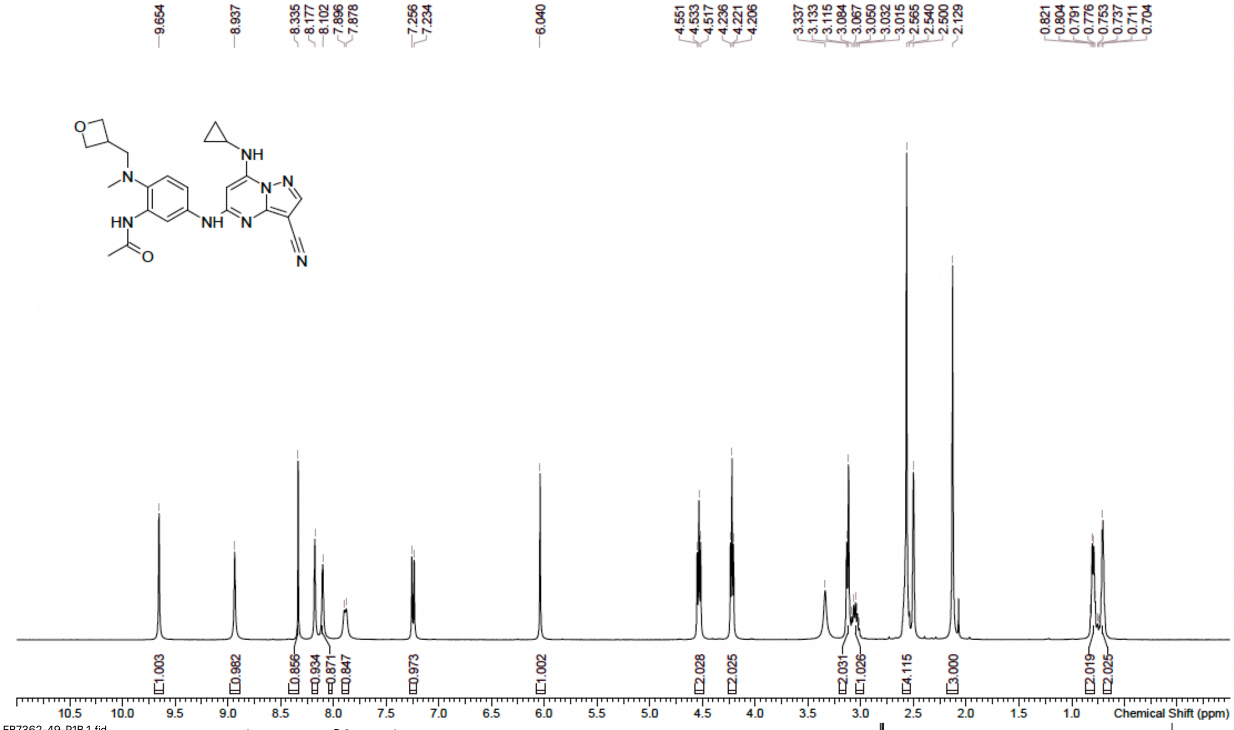

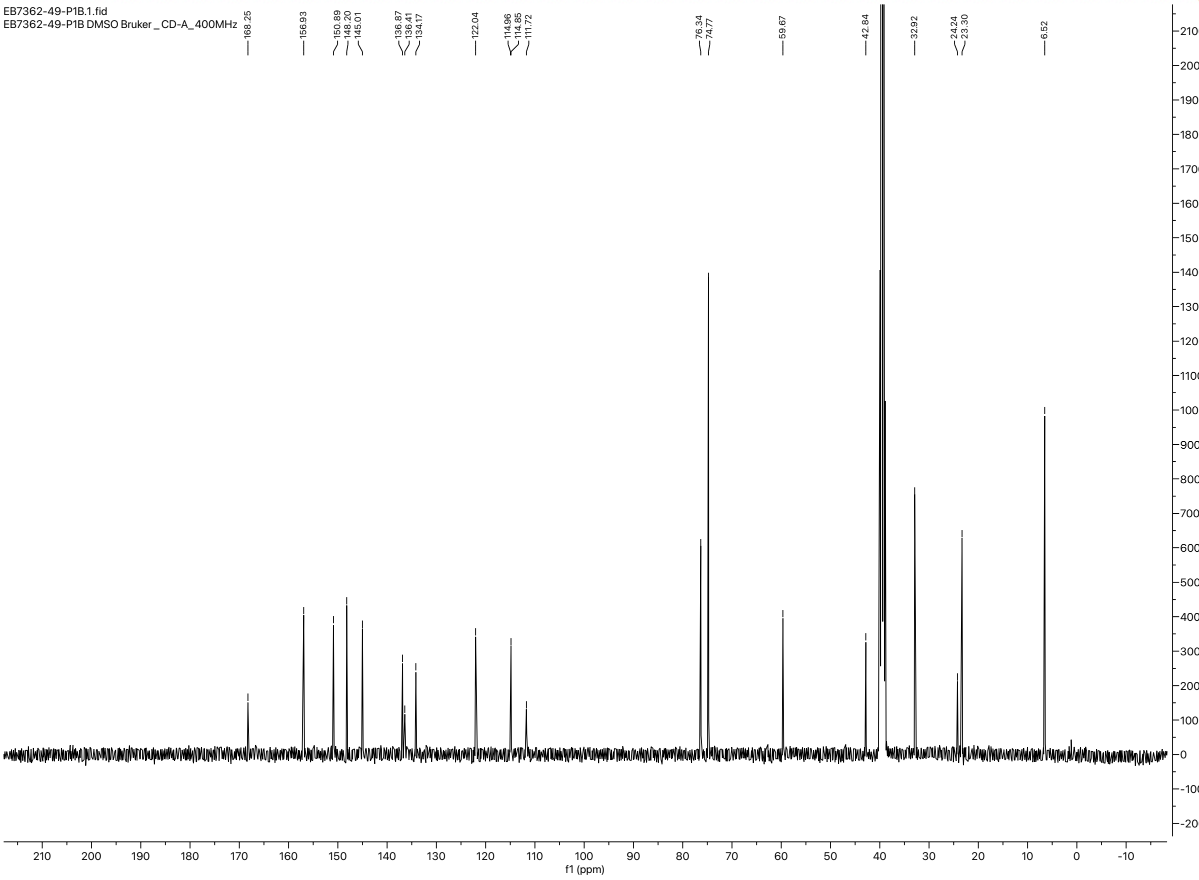

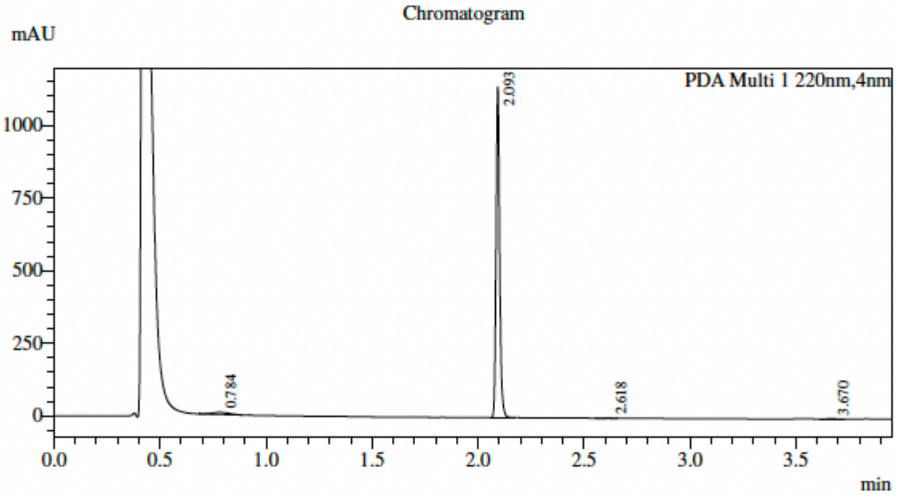


### Integration Result

PDA Ch1 220nm

| Peak# | Ret. Time | USP Width | Height | Height% | Area | Area% |
|---|---|---|---|---|---|---|
| 1 | 0.784 | 0.136 | 7387 | 0.643 | 40781 | 2.928 |
| 2 | 2.093 | 0.030 | 1139441 | 99.175 | 1346604 | 96.681 |
| 3 | 2.618 | 0.051 | 1021 | 0.089 | 2485 | 0.178 |
| 4 | 3.670 | 0.059 | 1071 | 0.093 | 2960 | 0.213 |
| Total | | | 1148920 | 100.000 | 1392830 | 100.000 |

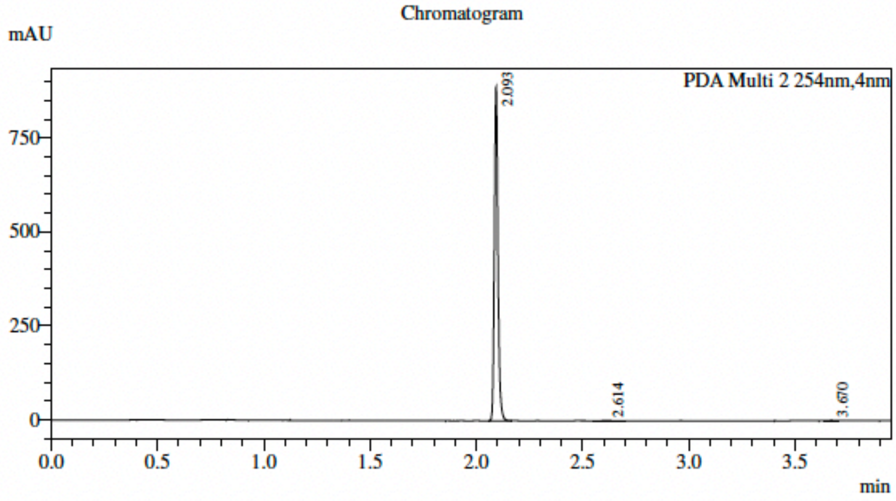


### Integration Result

PDA Ch2 254nm

| Peak# | Ret. Time | USP Width | Height | Height% | Area | Area% |
|---|---|---|---|---|---|---|
| 1 | 2.093 | 0.030 | 887654 | 99.743 | 1051968 | 99.517 |
| 2 | 2.614 | 0.051 | 1138 | 0.128 | 3058 | 0.289 |
| 3 | 3.670 | 0.044 | 1148 | 0.129 | 2043 | 0.193 |
| Total | | | 889941 | 100.000 | 1057069 | 100.000 |
